# Exclusion of the fittest predicts microbial community diversity in fluctuating environments

**DOI:** 10.1101/2020.07.22.216010

**Authors:** Shota Shibasaki, Mauro Mobilia, Sara Mitri

## Abstract

Microorganisms live in environments that inevitably fluctuate between mild and harsh conditions. As harsh conditions may cause extinctions, the rate at which fluctuations occur can shape microbial communities and their diversity, but we still lack an intuition on how. Here, we build a mathematical model describing two microbial species living in an environment where substrate supplies randomly switch between abundant and scarce. We then vary the rate of switching as well as different properties of the interacting species, and measure the probability of the weaker species driving the stronger one extinct. We find that this probability increases with the strength of demographic noise under harsh conditions and peaks at either low, high, or intermediate switching rates depending on both species’ ability to withstand the harsh environment. This complex relationship shows why finding patterns between environmental fluctuations and diversity has historically been difficult. In parameter ranges where the fittest species was most likely to be excluded, however, the beta diversity in larger communities also peaked. In sum, how environmental fluctuations affect interactions between a few species pairs predicts their effect on the beta diversity of the whole community.

## 1 Introduction

Natural environments are not static: temperature, pH, or availability of resources change over time. Many studies in microbiology, ecology and evolution have focused on responses to fluctuations in resource abundance in the regime of feast and famine periods (Hengge-Aronis, 1993; Vasi et al., 1994; Srinivasan and Kjelleberg, 1998; Xavier et al., 2005; Merritt and Kuehn, 2018; Himeoka and Mitarai, 2019). These models capture the dynamics within many natural ecosystems. For example, the gut microbiome of a host is exposed to fluctuating resources that depend on its host’s feeding rhythm, which may affect microbiota diversity (Cignarella et al., 2018; Li et al., 2017; Thaiss et al., 2014). In addition to their magnitude, environmental fluctuations can also differ in their time scales – for the gut microbiota, a host’s feeding rhythm may vary from hourly to daily, or even monthly if feeding depends on seasonal changes (Davenport et al., 2014; Smits et al., 2017) – or in the type of substrates taken up, which can be nutritious or harmful for the microbiota.

How environmental fluctuations (EFs) affect species diversity has been a highly contested topic in ecology. Here, EFs refer to changes that are not caused by the organisms themselves, but nevertheless affect their dynamics (e.g., abiotic resource supplies). The intermediate disturbance hypothesis argues that intermediate intensity and frequency of disturbances maximize species diversity (Connell, 1978; Grime, 1973) because highly competitive species dominate at a low level of disturbance, while only species that have adapted to the disturbance dominate at high disturbance (Grime, 1977). This hypothesis is controversial (Fox, 2013) and other relationships between disturbance and species diversity have been reported both empirically and theoretically (Mackey and Currie, 2001; Miller et al., 2011).

Another framework that is used to predict species diversity under EFs is the modern coexistence theory (Chesson, 1994), which explains the maintenance of diversity through species’ differing responses to and preferences for environmental conditions, which can vary over spatial and/or temporal scales through fluctuations (Amarasekare, 2019; Chesson, 2000a,b; Letten et al., 2018b; Barabás et al., 2018; Ellner et al., 2019). The modern coexistence theory divides environmental factors into those that are independent of species abundances (e.g., temperature) and those that depend on them (e.g., amounts of resources). The latter environmental factors mediate the sign and/or magnitude of interspecific interactions (Hoek et al., 2016; Piccardi et al., 2019; Zuñiga et al., 2019), and whether species tend to cooperate or compete can, in turn, drive community diversity and stability (Mougi and Kondoh, 2012; Coyte et al., 2015; Marsland III et al., 2019; Butler and O’Dwyer, 2018, 2020). Microbial communities often experience extreme environmental fluctuations that can alter interactions between species and hence affect species diversity (Rodríguez-Verdugo et al., 2019; Nguyen et al., 2020b).

Another important factor potentially influencing the outcome of interactions between species is demographic noise (DN) arising from randomness in birth and death events in finite populations. DN is negligible in large populations but strong in small ones, where it can lead to species extinction or fixation, and affect community composition (Roughgarden, 1979; Ewens, 2004). As EFs affect population sizes, they modulate the strength of demographic noise, leading to a coupling of EFs and DN. This interdependence has been understudied until recently, despite its potentially important consequences on eco-evolutionary dynamics (Wienand et al., 2017, 2018; West and Mobilia, 2020; Taitelbaum et al., 2020).

To understand how the interplay between EFs and DN affect community diversity, we set up a stochastic model of multiple species subject to a varying supply of nutrients and/or toxins. Our model then allows us to ask: How do EFs, coupled to DN, affect species interactions and diversity?

We include toxins in our model as they are important in natural communities, but often missed in similar models. Toxins that typically come to mind are pesticides or antibiotics (Pérez et al., 2005; Xu et al., 2011), but in principle they can be anything that inhibits microbial growth compared to some optimal condition. For example, oxygen is harmful to anaerobic microbes (Guittar et al., 2021) and bile acids are toxic to microbes in the human gastrointestinal tract (Ruiz et al., 2013; Molinero et al., 2019). These toxins can, however, be degraded by microbes. In the example of bile acids, *Lactobacillus* and *Bifidobacterium* strains degrade them by producing bile-salt hydolases and other extracellular enzymes (Ruiz et al., 2013; Molinero et al., 2019). Such degradation of toxic compound by microbes is expected to be quite common, as microbes will be selected to become resistant to toxins by diminishing environmental toxicity. In our model then, toxins contribute to environmental harshness and reduce population sizes, which can modulate the strength of DN. We also expect toxins to affect inter-species interactions, as species that absorb or degrade them can potentially facilitate the growth of others (Hoek et al., 2016; Piccardi et al., 2019; Zuñiga et al., 2019).

The next section describes our theoretical model, explaining how DN and EFs are implemented and how species interactions and diversity are measured (section 2). We first use it to explore how often one single species goes extinct due to environmental switching coupled with DN (section 3.1). We then add a second slower-growing species, still focusing on the behavior of the first, and ask how its extinction probability is affected by the slower-grower, and how different properties of the two species change the effects of the switching rate (section 3.2). We find that sensitivity to toxins increases the strength of DN because it decreases species abundances. At toxin sensitivities that are high enough to increase DN but low enough that either species is likely to survive, the slower-growing species can outcompete the fast-growing one (we call this phenomenon *exclusion of the fittest*, sections 3.3 and 3.4), a result that is a direct consequence of coupling DN and EFs. Finally, we expand our model to larger communities and show that our first analysis of the exclusion of the fittest species predicts beta diversity patterns in larger communities (section 3.5): community composition is the most diverse when strong species are most likely to be excluded. Section 4 discusses the importance of coupling EFs and DN on species interactions and diversity. Finally, we compare our results with the intermediate disturbance hypothesis and the modern coexistence theory. Technical and computational details are given in a series of Appendices.

## 2 Model & Methods

In order to investigate the interdependence of EFs and DN on species interactions and diversity, we study an idealized chemostat model that combines DN and environmental switching (Fig. 1). The chemostat is meant to represent natural ecosystems subject to in- and outflow, such as a gut or a river. Within the chemostat, we consider a well-mixed population consisting of *N* species of bacteria, and *N/*2 types of resources as well as *N/*2 types of toxins (without loss of generality, *N* is assumed to be even). Our system can exhibit different levels of species richness at equilibrium depending on parameter values: only one species may persist as species interact with all resources and toxins, or all species may coexist as the environment has as many limiting factors (resources and toxins) as the number of species. The special case of a model with a single resource and a single toxin in a mono-culture is considered for completeness in Section 3.1. Although many models ignore the role of toxins, they play two important roles in our model: first, sensitivities to toxins changes the strength of DN because population sizes of sensitive species are low. Second, toxins can enable a pair of species to facilitate each other (Piccardi et al., 2019) because toxin absorbance or degradation decreases the death rate of other species. In this manuscript, we consider toxins that are not released from cells when they die, e.g., antibiotics that inhibit DNA or RNA synthesis (Cangelosi and Meschke, 2014), although the release of toxins from dead cells could be introduced with a small modification of our model (see Huang et al. (2013) for example). The absence of toxins is similar to having a low toxin sensitivity (Fig. A.22).

**Figure 1:**
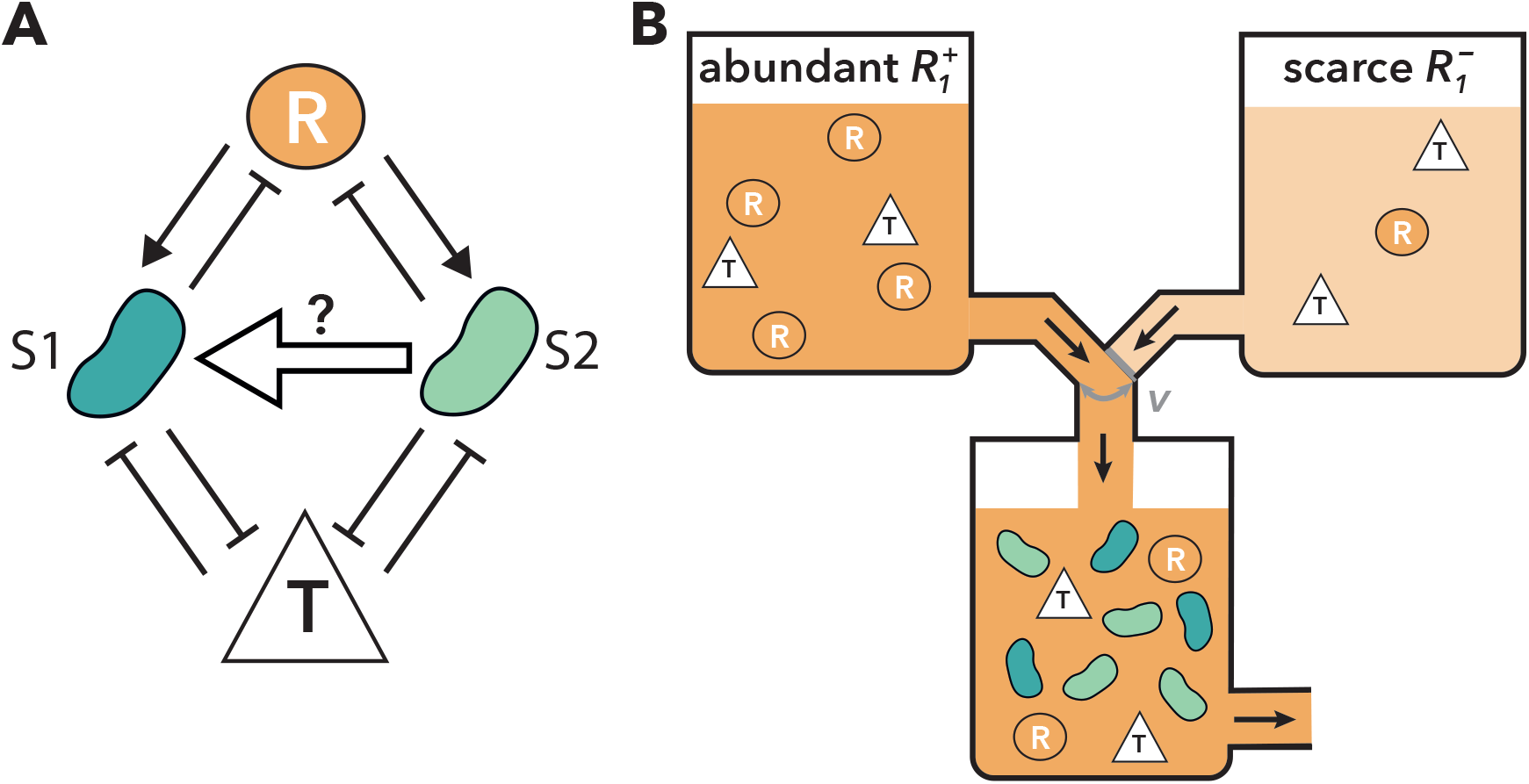
Schematic illustration of the model. A chemostat model with environmental switching. A: Interaction network when *N* = 2. A → B represents that A increases B while A ⊣ B represents that A decreases B. Two species compete for the same resource (R in a circle) but are killed by the same toxic compound (T in a triangle). As a proxy for species interactions, we follow the net effect of the slower-growing species 2 on species 1 (large arrow from species 2 to 1). B: An example of a chemostat model with environmental switching and *N* = 2. Environmental switching is realized by changing the media flowing into a chemostat. In this example, the current environmental condition is abundant resource supply 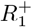.

In our model, communities follow a continuous-time multi-variate birth-and-death process (see, e.g. Novozhilov et al. (2006); Allen (2010)) in a time-fluctuating binary environment (see, *e*.*g*., Wienand et al. (2017, 2018); West and Mobilia (2020); Taitelbaum et al. (2020)). More specifically, we consider that at time *σ* the community consists of an amount *r*_*i*_(*σ*) of resources of type *i* (*i* = 1,. .., *N/*2), an amount *t*_*j*_(*σ*) of toxin of type *j* (*j* = 1,. .., *N/*2), and an abundance *s*_*k*_(*σ*) of microbial species *k* (*k* = 1,. .., *N*). Here, resources are assumed to be nutrients for all species, allowing them to grow at different rates, while the toxins kill all species at different rates when species have different toxin sensitivities. hence the species extinction probability.

In a static environment (no EFs), this chemostat model evolves according to the birth-and-death process defined by the following “birth” and “death” reactions:

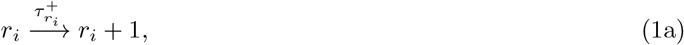

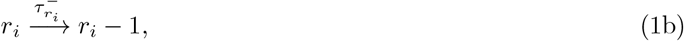

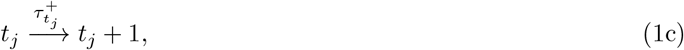

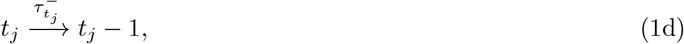

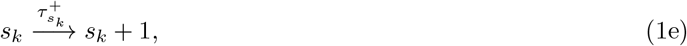

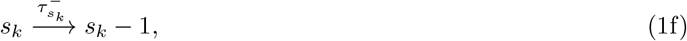

occurring with transition rates:

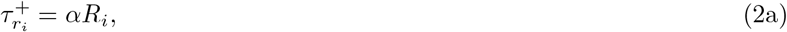

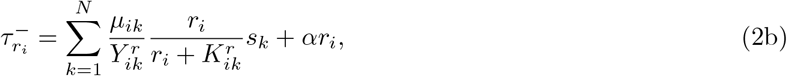

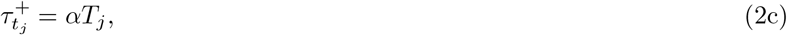

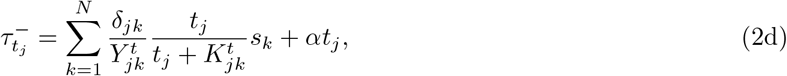

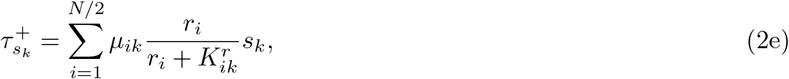

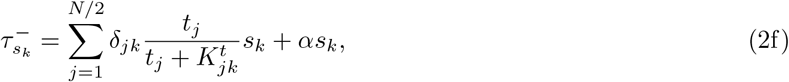

where *α* is the dilution rate of the chemostat, *ξ* = ±1 (see below) represents changing environmental conditions in terms of in-flowing resource and/or toxin amount. *R*_*i*_(*ξ*) (*T*_*j*_(*ξ*)) is resource *i*’s (toxin *j*’s) supply under the environmental condition 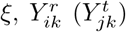 is species *k*’s biomass yield for resource *i* (toxin *j*), *μ*_*ik*_ is the maximum growth rate of species *k* by resource *i, δ*_*jk*_ is the maximum death rate of species *k* by toxin *j* (species *k*’s sensitivity to toxin *j*), and 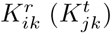is the amount of resource *i* (toxin *j*) that gives the half-maximum growth (death) rate for species *k* (see also Table A.1). These transition rates hence reflect that (i) the amounts of resources and toxins increase depending on the product of their in-flow concentrations and the dilution rate, (ii) the amounts of resources and toxins decrease with the dilution rate and with the consumption/absorption by species, (iii) the growth and death rates depend in Monod functional forms on the amounts of resources and toxins, respectively, (iv) the dilution rate *α* sets the time scale at which the environment attains the state of abundance or scarcity (after a time *∼* 1*/α*), see below and Appendix 2.2 and (v) all effects are additive. When *R*_*i*_ and *T*_*j*_ are constant, the environment is static and the birth-and-death process defined by Eqs (1a) – (2f) naturally accounts for the DN arising in the population, which is the sole source of noise.

We model EFs by considering a time-fluctuating binary environment resulting in *R*_*i*_ and/or *T*_*j*_ to be *dichotomous random variables*, i.e., *R*_*i*_ = *R*_*i*_(*ξ*(*σ*)) and/or *T*_*j*_(*ξ*(*σ*)), where *ξ*(*σ*) = ±1 represents the binary state of the environment (*ξ* = 1 represents a mild environment while *ξ* = –1 represent a harsh environment). We hence assume that *R*_*i*_ and/or *T*_*j*_ switch between two values at a rate *ν*, according to a time-continuous colored dichotomous Markov noise (random telegraph noise) (Bena, 2006; Horsthemke and Lefever, 2006)

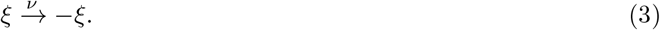

We call *ν* a “switching rate” because we implement a symmetrically switching environment as a simple example of EFs (but see Appendix 2.2 for a further discussion of this choice and Appendix 5 for other forms of EFs). Although we use arbitrary units in the model, the unit of the switching rate applies to all events. For example, if the unit for the dilution rate is per hour, *ν* = 10^0^ means that the environment switches every hour on average. We investigate three environmental switching scenarios, where either or both resource and toxin supplies fluctuate over time, see Table 1. In the main text, we focus on the scenario where only resource supplies switch between abundant and scarce supplies (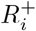 and 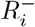, respectively, such that 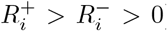 while the amounts of toxin supplies remain constant over time 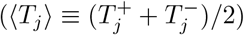; see Appendix 3 for the remaining scenarios. The initial resource supply in the main text is 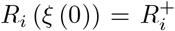 with probability 0.5 and otherwise 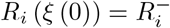.

**Table 1:** Different scenarios of environmental switching. 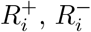, and ⟨*R*_*i*_⟩ represent abundant, scarce, and mean resource supplies, respectively. 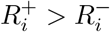 and 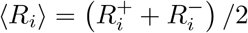 for *i* = 1,. .., *N/*2. Similar representation and relations hold for toxin supply *T*_*j*_. In each condition, *ξ* = 1 (*ξ* = −1) means mild (harsh) environment, respectively.

| Scenario | $R_i (\xi = 1)$ | $R_i (\xi = -1)$ | $T_j (\xi = 1)$ | $T_j (\xi = -1)$ |
| --- | --- | --- | --- | --- |
| 1 | $R_i^+$ | $R_i^-$ | $\langle T_j \rangle$ | $\langle T_j \rangle$ |
| 2 | $\langle R_i \rangle$ | $\langle R_i \rangle$ | $T_j^-$ | $T_j^+$ |
| 3 | $R_i^+$ | $R_i^-$ | $T_j^-$ | $T_j^+$ |

We assume that *ξ* switches symmetrically between the states ±1 (see Taitelbaum et al. (2020) and Appendix 5 for the cases of asymmetric switching). In all our simulations, *ξ* is stationary and thus *ξ* has zero mean, ⟨*ξ*(*σ*) ⟩ = 0, autocorrelation ⟨*ξ*(*σ*)*ξ*(*σ′*) ⟩ = exp (–2*ν*| *σ − σ′*|), where ⟨*·*⟩ denotes the ensemble average over the environmental noise, and finite correlation time 1*/*(2*ν*). A great advantage of dichotomous noise is its simplicity: *ξ* is bounded (*R*_*i*_ and *T*_*j*_ are always well defined) and straightforward to simulate. This choice allows us to model suddenly changing environmental conditions, which reflect situations in microbial life such as exposure to resource or toxin oscillations that can be reproduced in lab-controlled experiments (Sunya et al., 2013). Other forms of EFs are also possible, e.g. *ξ* could be a Gaussian random variable, but then *R*_*i*_ and *T*_*j*_ would be unbounded and vary continuously and could take unrealistic values. Modeling EFs with Eq (3) is arguably the simplest biologically-motivated choice to couple fluctuations in resource/toxin supplies with demographic noise, and allows us to investigate questions that are not specific to dichotomous noise, see Appendix 2.2. Interestingly, the analysis of the long-term dynamics of the two-species, one-resource-one-toxin model under symmetric dichotomous noise in Appendix 2.2 reveals that this simple form of environmental variability leads to distributions of the total population size *n* (i.e., the sum of species, resources and toxins abundances *n* ≡ Σ_*k*_ *s*_*k*_ + Σ_*i*_ *r*_*i*_ + Σ_*j*_ *t*_*j*_) that varies greatly with the rate of switching relative to the dilution rate: when *ν/α* ≫ 1 (fast switching), the total population size *n* is unimodal; when *ν/α* ≪ 1 (slow switching), *n* is bimodal and fluctuates between two very different values; intermediate scenarios interpolating between unimodal and bimodal distribution of *n* arise when *ν/ α* ∼ 1 (intermediate switching), see Wienand et al. (2017, 2018); West and Mobilia (2020); Taitelbaum et al. (2020) and Fig. A.6. This results in an explicit coupling of DN to EFs in multi-species communities, via the modulation of the DN intensity by *ν/ α*. This is a distinctive feature of our model comparing previous studies. For example, some models (Leigh, 1981; Engen and Lande, 1996; Kalyuzhny et al., 2015) do not couple DN and EFs, while other models (Engen and Lande, 1996; Kamenev et al., 2008; Chisholm et al., 2014; Fung et al., 2015) study the coupling of DN and EFs in single species scenarios. See Appendix 2.2 for more detailed discussion and analysis.

The master equation for this model is defined by combining the dynamics of the amounts of resources and toxins, abundances of species, with the environmental switching (Eqs (1a) - (3)):

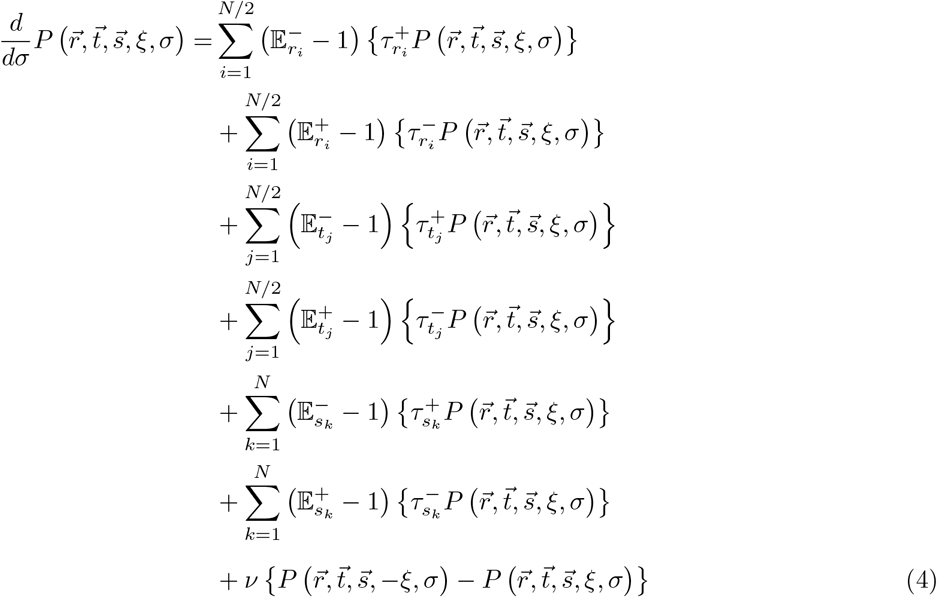

Where 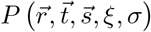 gives the probability to find the population in state 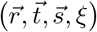 at time *σ*, with 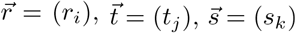. Here, 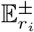 is a shift operator such that

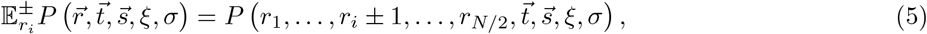

and 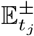 and 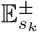 are the equivalent shift operators for *t*_*j*_ and *s*_*k*_, respectively. Note that 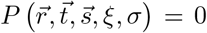 whenever any of *r*_*i*_, *t*_*j*_, *s*_*k*_ *<* 0. The first to sixth lines on the right-hand-side of Eq (4) represent the birth-and-death processes (1), while the last line accounts for environmental switching.

The master equation (4) fully describes the model dynamics and can be simulated exactly with the Gillespie algorithm (Gillespie, 1977). Owing to the stochastic nature of the model, after a time that diverges exponentially with the community size, DN will cause the eventual collapse of the population (Spalding et al., 2017; Assaf and Meerson, 2017). This phenomenon is practically unobservable when resource levels remain sufficiently large, i.e. if 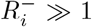, and the population settles in a long-lived quasi-stationary distribution. Here we focus on the quasi-stationary regime that is attained when distributions of species abundances appear to be stationary for a long time (see supplementary video): the distributions of two species’ abundances change little from time *σ* = 130 onward when the parameter values are as shown in Table A.1 with *ν* = 10^*-*1^ and *δ* = 0.2, which are typical parameter values used in this study. It would be reasonable then to set *σ*_*end*_ = 200 expecting that species’ abundances reach a quasi-stationary state in many of our chosen parameter values.

### 2.1 Evaluating species interactions

First, we analyze how the net effect of species interactions (i.e., resource competition and facilitation via detoxification) can change under DN and EFs by measuring the extinction probabilities in the presence or absence of another species at time *σ*_*end*_ (see Appendix 1.1 for details). We begin with two species (*N* = 2) where the sign and magnitude of species interactions can change because the amounts of resources and toxins can change the net effect of resource competition and facilitation via detoxification (Piccardi et al., 2019). We used parameter values such that species 1, if it persists, always outcompetes species 2 in the absence of DN (i.e., species 1 grows faster than species 2, see Appendix 2.1 for analysis and Table A.1 for parameter values) to identify the effects of DN. Importantly, EFs alone do not change species 1’s extinction probabilities in mono-versus co-culture in the absence of DN. Under DN coupled with EFs in our chosen parameter range, either of the two species or both species tend to go extinct. As a proxy for interactions, we focus on the net effect of species 2 on species 1, which is defined by the extinction probability of species 1 in mono-culture minus that in co-culture with species 2 (the so-called *difference in extinction probability*, see also Eq (A.1)): species 2 has a negative (positive) effect on species 1 if species 1 more (less) frequently goes extinct in co-culture with species 2 than in mono-culture. When species 2 has a negative effect on species 1, one can consider two possibilities in co-culture: (i) species 2 outcompetes species 1 (see Fig. 4) or (ii) both species 1 and 2 go extinct. As explained in section 3.3, we focus on the former probability, the so-called *probability of exclusion of the fittest* to understand the net effect of species 2 on species 1. We performed 10^5^ simulations for each switching rate and toxin sensitivity in mono- and co-cultures. We calculated 95% of highest posterior density intervals (HPDI) to measure the uncertainty of species 1’s extinction probabilities (see Appendix 1.1) but these intervals are too small to be visible on our plots due to the large number of simulations we ran.

### 2.2 Evaluating species diversity

To explore how species diversity changes with the switching rate, we ran simulations at different numbers of species ranging from *N* = 2 species to *N* = 10 species and different mean toxin sensitivities (see Appendix 1.3 for details). For each condition (one number of species and one mean toxin sensitivity), we sampled 100 sets of parameters from certain probability distributions. These sets of parameters represented 100 communities composed of *N* species. In this analysis (section 3.5) we consider a more general scenario and all previous simplifying assumptions on the parameter values are relaxed: all species have randomly drawn (see Appendix 1.3), and hence typically different, growth and death rates.

The dynamics of each community were independently simulated 100 times to see whether the species composition was robust against DN and EFs. These 100 replicate runs can be seen as 100 independent “patches”that initially consist of the same species set, but no species migrate from one patch to another. We measured the beta diversity of each community (Jost, 2007; Chao et al., 2012) and species richness (number of surviving species) as functions of the environmental switching rate in the quasi-stationary distribution (at time *σ*_*end*_). Beta diversity accounts for the heterogeneity of each community across 100 replicates For example, if beta diversity is larger than one but species richness is one in all replicates, different species fixate in each replicate. In contrast, beta diversity is one if all communities show identical species compositions; for example, in the two-species scenario with parameter values shown by Table A.1 and in the absence of DN, species 1 always outcompetes species 2 and thus beta diversity is one. This baseline corresponds to a perfectly deterministic scenario. One could instead consider using another baseline corresponding to a perfectly stochastic or neutral scenario where all species’ parameter values are identical (e.g., Fig. A.21), in which case species compositions are determined only by demographic noise coupled with environmental fluctuations. We choose to focus on the deterministic baseline, as beta diversity is always = 1 regardless of the number of species, their parameter values, and the environmental switching rate. In contrast, beta diversity in the neutral scenario changes with these parameter values, making it more difficult to compare across conditions.

We compared the patterns of beta diversity of two- or ten-species communities with the probability of exclusion of the fittest in species pairs sampled from these communities (see also Appendix 7). The sampled species pairs may stably coexist, in which case the fittest species in a pair is the one that is more abundant in the absence of any noise. If both species go extinct in the absence of noise, either of the two species is randomly labeled as the fittest. This labeling generalizes what is used in the species interaction analysis above where species 1 (the fittest species) always grows faster than species 2, such that their long term coexistence is impossible in the absence of DN.

### 2.3 Statistical analysis

Statistical analysis was performed with Python 3.7.6 incorporating Scipy 1.4.1. and pymc3 3.10.0. For statistical tests of Pearson’s correlation and Spearman’s rank-order correlation, scipy.stats.pearsonr and scipy.stats.spearmanr were used, respectively. For calculation of HPDIs, pymc3.stats.hpd was used.

## 3 Results

### 3.1 Toxin sensitivity determines single species’ response to coupled DN and EFs

To establish our intuition on how the coupling of EFs and DN affects the dynamics, we first analyse the extinction probabilities of a single species (species 1) in mono-culture with one type of resource and toxin (Fig. 2A). As the switching rate decreases, the duration of the harsh or mild environments become longer. In the presence of DN, this duration determines whether or not the species goes extinct, together with its sensitivity to toxins in the environment, which can be seen to modulate environmental harshness. When the switching rate is very low (*ν →* 0), the species is exposed to the static environment with either abundant or scarce resources (with probability 0.5, respectively) depending on the initial environmental condition *ξ*(0): it mostly goes extinct under scarce resource supplies even when their sensitivity to toxins is low (Fig. 2B). On the other hand, abundant resource supplies maintain species 1 with some probability even if it is very sensitive to the toxin (Fig. 2D). Over many simulations, low fluctuation rates therefore result in a bimodal distribution of the species’ abundance (e.g., Fig. 2B). At the other extreme, very high switching rates (*ν → ∞*) expose the bacteria to an environment with mean abundance of resources (Wienand et al., 2017, 2018; West and Mobilia, 2020; Taitelbaum et al., 2020). This is enough to rescue the species with a low toxin sensitivity (Fig. 2B) but not with a high sensitivity (Fig. 2D). At an intermediate toxin sensitivity (Fig. 2C), the worst situation lies in the intermediate fluctuation rate: the duration of the harsh environment is long enough to drive them extinct, but the time with abundant resource supply is not long enough to rescue them fully. In sum, even when only a single species is present, we see non-trivial patterns in its response to EFs coupled with DN, which depends on its sensitivity to toxins.

**Figure 2:**
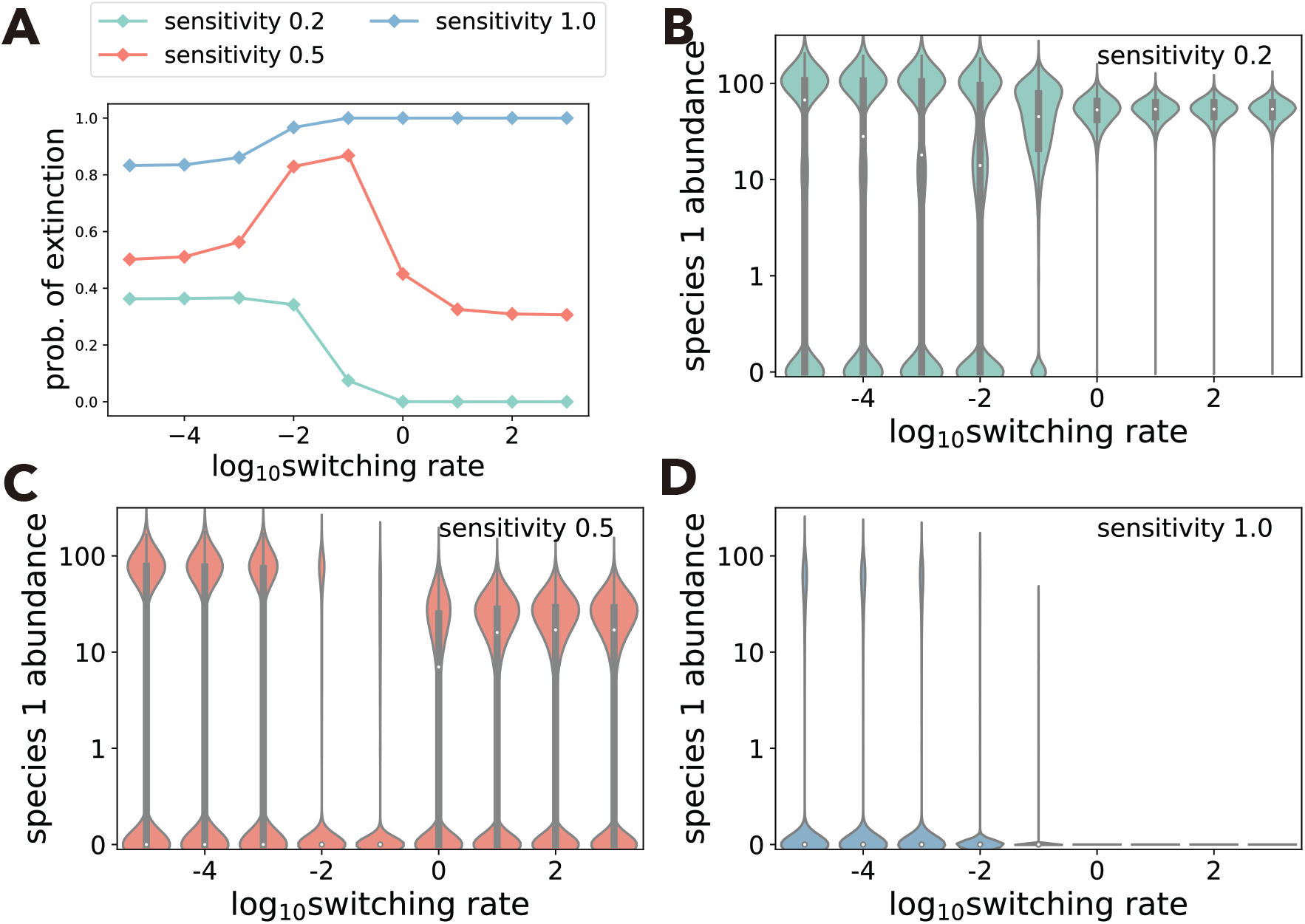
Extinction probability and species abundance in mono-culture. A: Extinction probability of species 1 in mono-culture when the toxin sensitivity is low (green), moderate (orange), or high (blue). 95% HDPIs are too small to see. B–D: violin plots of species 1’s abundance at the end of 10^5^ simulations with a low (B), moderate (C), or high (D) toxin sensitivity. White dots and black bars represent median values of the abundances and their interquartile ranges, respectively.

### 3.2 Toxin sensitivity changes how switching rate affects two-species competition

Next, we add another species (species 2) that grows slower than species 1 into the environment and ask how it interacts with species 1 in our model. Rather than measuring interactions through the effect of each species on the other’s abundances, we focus on species 1 and analyze how its extinction probability is affected by the presence of species 2, compared to mono-culture (Fig. 2). Our reasoning is that (i) species 1 should always out-compete species 2 in the absence of DN, and that (ii) we already know the extinction probability of species 1 under EFs and DN in mono-culture; measuring any deviation from the mono-culture outcome allows us to quantify how likely it is for the fitter species to be lost in a given community. Such species loss events can be seen as ecological drift. We again explore changes in the species’ toxin sensitivity, as we learned above that it affects species abundances via DN (Fig. 2), but also because we expect it to affect species interactions (Piccardi et al., 2019). For now, we varied sensitivity to toxins simultaneously for both species, an assumption that we relax later.

When both species were highly sensitive to the toxin, species 2 had a positive effect on species 1, reducing its extinction probability. This occurs because in the simulations, toxic compounds are degraded more quickly in co-culture than in mono-culture due to the larger initial number of individuals (*s*_1_(0) + *s*_2_(0) *> s*_1_(0)). This effect can be recapitulated by a mono-culture with larger initial abundance (Fig. A.20). A larger total initial species abundance in co-culture decreases death rates, which outweighs competition for nutrients in toxic environments (Piccardi et al., 2019). However, for most parameter values in Fig. 3A, species 2 has a negative effect on species 1 by increasing its extinction probability. We therefore focus on competitive interactions in the main text and consider positive interactions in Appendix 4.

**Figure 3:**
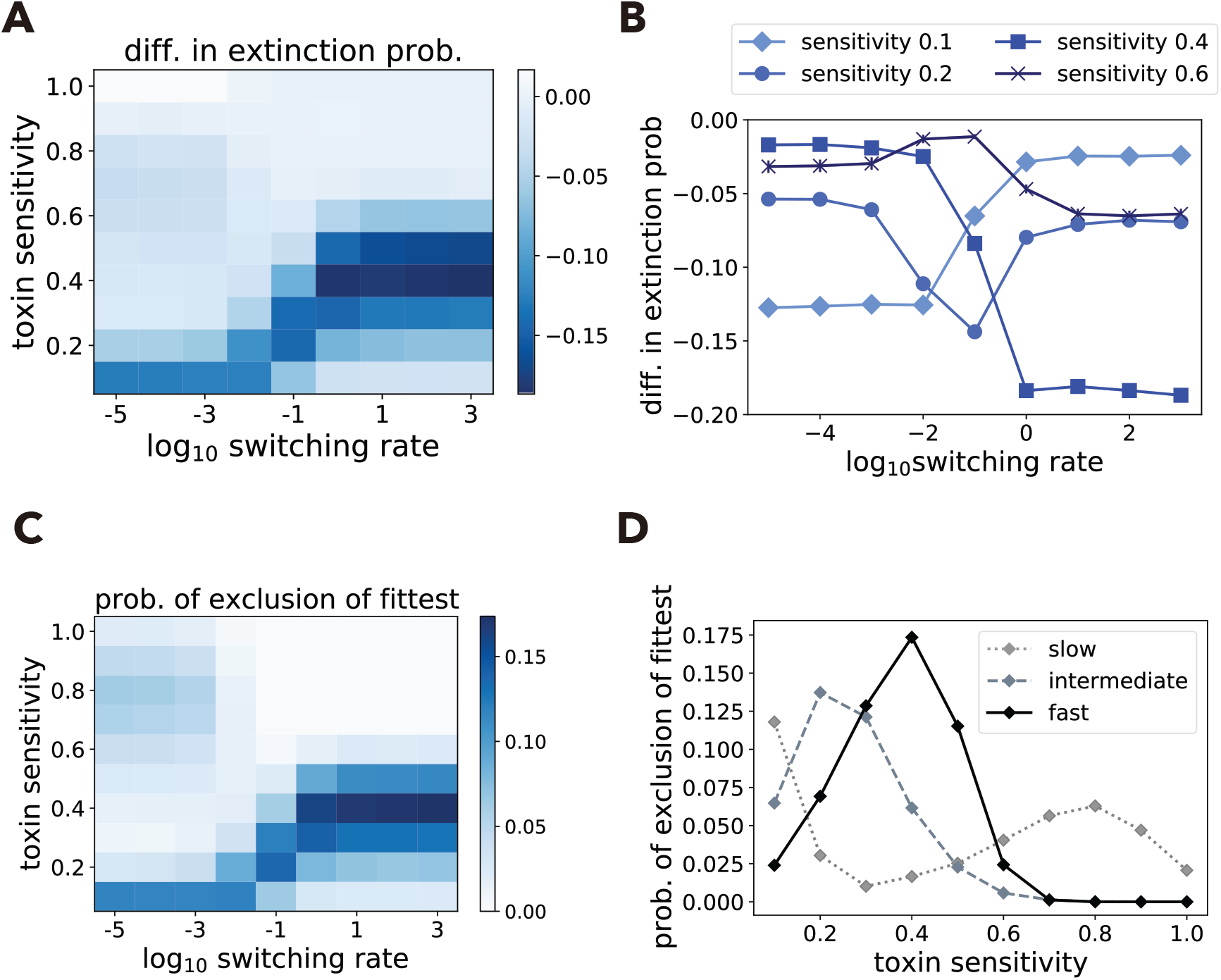
Species interaction strength changes differently over the switching rate. A: Difference in species 1’s extinction probability in mono-culture minus co-culture with species 2 (Eq A.1) changes over the switching rate *ν* and the two species’ identical toxin sensitivities. In most of the parameter space, species 2 has a negative effect on species 1 (i.e., species 2 increases the extinction probability of species 1). B: Some illustrative examples from panel A plotted differently to show how species 2’s effect on species 1 changes over the switching rate at given toxin sensitivities. The difference in extinction probability can monotonically increase (toxin sensitivity 0.1), monotonically decrease (toxin sensitivity 0.4), or non-monotonically change with a minimum (toxin sensitivity 0.2) or a maximum (toxin sensitivity 0.6) value at an intermediate switching rate. C: Probability that species 2 persists but species 1 goes extinct (i.e., exclusion of the fittest) over the switching rate and the toxin sensitivity. D: probabilities of exclusion of the fittest over the toxin sensitivity, when the environmental switching rate is slow (*ν* = 10^*–*5^), intermediate (*ν* = 10^*–*1^), or fast (*ν* = 10^3^). In panels B and D, 95% HDPIs are too small to see.

**Figure 4:**
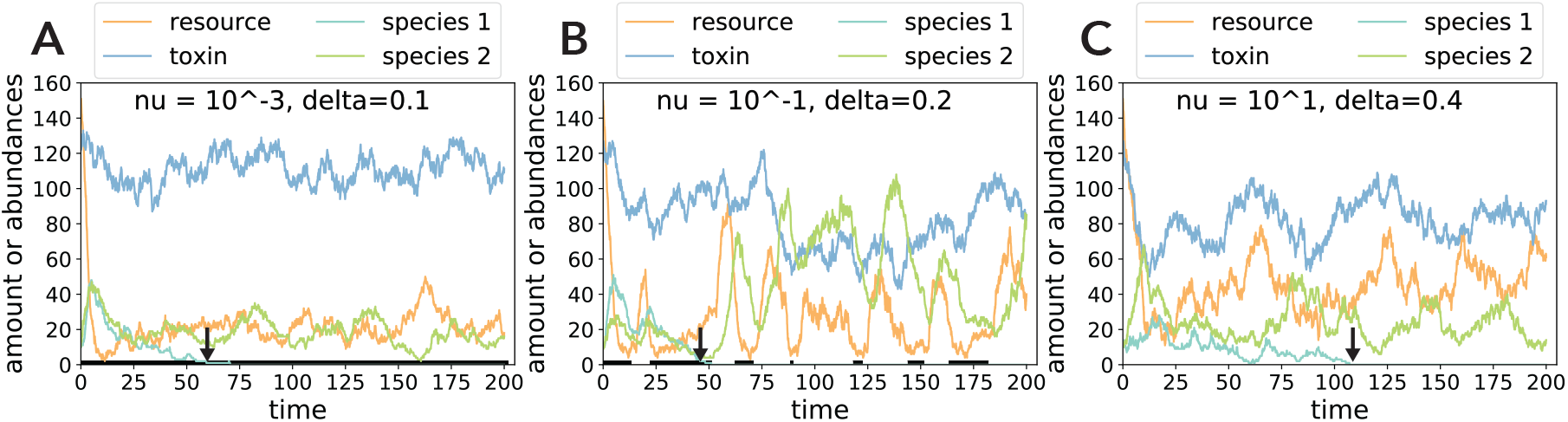
Examples of the dynamics with exclusion of the fittest. In these examples (A: *ν* = 10^*–*3^, and *δ* = 0.1, A: *ν* = 10^*–*1^, and δ = 0.2, C: *ν* = 10^1^, and = 0.4), species 1 goes extinct but species 2 survives at the end of simulation *σ*_*end*_ = 200 due to the DN-EFs coupling. Black lines on the *x*-axis represents times when the resource supply is scarce (*ξ*(*t*) = *-*1) while white lines represent times when the resource supply is abundant (*ξ*(*t*) = 1). In panels A and B, species 1 decreases its abundance and goes extinct during the scarce resource supply condition (pointed by arrows). In panel C, the environmental conditions are not shown because they are not visible due to the fast switching.

As in the single species case (Fig. 2), the extinction of species 1 was highly dependent on the toxin sensitivity of the two species, as we varied the fluctuation rate: monotonically increasing, monotonically decreasing, or non-monotonically changing with a minimum or maximum value at an intermediate switching rate (Fig. 3B). Interestingly, this pattern does not match the single-species behavior (compare Fig. 2A and 3AB).

### 3.3 Behavior at extreme switching rates explains non-monotonic changes in exclusion of the fittest

To better understand why the faster-grower goes extinct in the observed parameter ranges (Fig. 3B), we decompose species 2’s effect on species 1 into the probability that species 2 persists but species 1 goes extinct (hereafter, called *probability of exclusion of the fittest*) and the probability of any other outcome (e.g., extinction of both species, see Eq (A.2)). We focus on the probability of exclusion of the fittest, as it explains the variation in species interaction strength in most cases (compare Figs. 3A and C) and investigate how it changes over the switching rate and depends on toxin sensitivity.

We again let the two extreme switching rates guide our intuition (see also Fig. A.4). At a very slow rate (*ν →* 0), the resource supply remains either scarce or abundant, while a very fast environmental switching rate (*ν →* ∞) drives resource supply to the mean concentration. We explore system behavior at these three constant resource supplies (scarce, abundant or mean) and different toxin sensitivities, which together represent how harsh the environment is. When the environment is harsh – due to resource scarcity, high toxin supplies, or high toxin sensitivity –, both species are most likely to go extinct rather than to outcompete each other (Figs. 5A – C). As toxin sensitivity goes down and species survival becomes more likely, DN becomes more important and we see a higher probability that even the fitter species (species 1) will be excluded. The more resources are available, the more likely it is that species survive – in particular, that species 1 out-competes species 2, and the peak of competitive exclusion moves to a higher toxin sensitivity (see arrows in Figs. 5A – C). When it is easy for both species to survive (toxin sensitivity is low and/or resources are abundant), DN no longer plays an important role and the faster-growing species 1 is unlikely to be excluded. ntuitively then, the exclusion of the fittest is caused by the coupling of DN with EFs: Harsh environments, where both species’ abundances are positive but low (see Appendix 2.2), lead to stronger DN and a higher probability of extinction of the fittest species compared to a static or mild environment (Fig. 4). It is important to stress that the exclusion of the fittest never occurs without DN, regardless of EFs (see Appendix 2).

**Figure 5:**
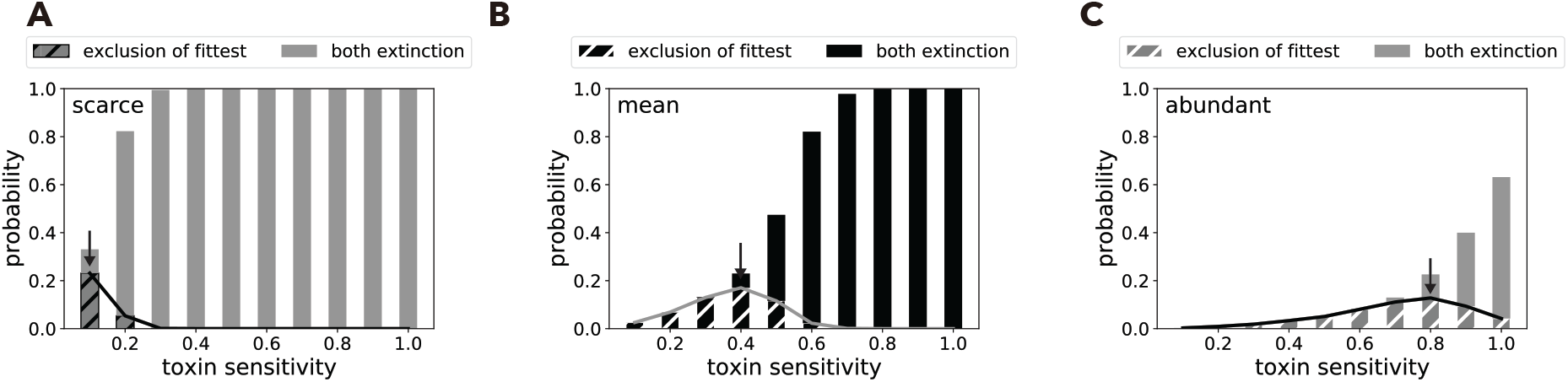
Exclusion of the fittest explains how DN changes with toxin sensitivity and resource supply. Analysis of exclusion of the fittest predicts at which toxin sensitivity DN is strongest. A – C: In the absence of environmental switching, the probability of exclusion of the fittest (i.e., probability that species 2 excludes species 1 and survives, shown by solid lines and hatched bars) is uni-modal over the toxin sensitivity while the probability of both species extinction (bars) monotonically increases. The toxin sensitivities giving the peak values of probabilities of exclusion of the fittest (critical toxin sensitivities, pointed by black arrows) depend on the resource supply: scarce 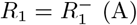, mean *R*_1_ = ⟨*R*_1_⟩ (B), or abundant 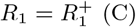. 95% HDPIs are not observable as they are very small.

Hereafter, we refer to the toxin sensitivity that maximizes the probability of exclusion of the fittest in the absence of environmental switching as the “critical toxin sensitivity” (arrows in Figs. 5A – C). We see two critical toxin sensitivities (at 0.1 and 0.8 in Figs. 3C and D, Figs. 5A and C) at *ν →* 0 that correspond to the long time spent with either scarce or abundant resources. Instead, at *ν →* ∞, where resources remain at mean abundance, there is a single critical toxin sensitivity (at 0.4 in Figs. 3C and D) where exclusion of the fittest is most likely (Fig. 5B). Toxin sensitivities between these critical values can show a maximum or minimum probability of exclusion of the fittest at an intermediate switching rate, resulting in the rugged landscape of Fig. 3C (see Appendix 1.2 for more detail).

### 3.4 Competition strength changes non-monotonically under different scenarios

We have shown that using a given set of parameters, the rugged landscape shown in Fig. 3C causes the competitive exclusion of a faster-growing species to either increase, decrease or vary non-monotonically across switching rates, depending on toxin sensitivity. We next explore the generality of this finding. In the appendix, we explore scenarios where (i) switching occurs in toxin rather than resource supplies, where (ii) both resource and toxin supplies switch (Table 1, see Appendix 3), or where (iii) we change the amounts of scarce and abundant resource supplies (Appendix 4).

In all these scenarios, the landscapes of species 1’s difference in extinction probability and probability of exclusion of the fittest are qualitatively similar (Figs.3 C and D, A.7, and A.9). However, each scenario differs in the three critical toxin sensitivities and likelihood that the difference in extinction probability non-monotonically changes over the switching rate. Accordingly, we asked whether the distances between critical toxin sensitivities might predict the probability of observing non-monotonic behavior. In Table A.2 and Fig. A.14, we show that the distance between critical sensitivities under harsh and mean environments (i.e., very fast environmental switching) correlates positively with the likelihood of observing non-monotonic effects of the switching rate on competition (Fig. A.14, black circles; Spearman’s *ρ* = 0.77, P-value: 0.043), but no significant correlation was found with the distance between the critical toxin sensitivities under the mean and mild, or the harsh and mild environments (Fig. A.14, grey diamonds and cross marks; Spearman’s *ρ* = –0.22, P-value: 0.64, and *ρ* = 0.42, P-value: 0.35, respectively). Therefore, the non-monotonic change of the difference in extinction probability is likely when there is a large difference in the critical toxin sensitivities under harsh and mean conditions.

### 3.5 Beta diversity changes similarly to exclusion of the fittest

In the previous sections, we focused on interactions between two species and the conditions under which one may drive the other extinct. Ultimately, however, our interest is to predict how whole communities comprised of tens, hundreds or even thousands of species are affected by fluctuations in the environment.

We set up a model of 100 communities composed of between 2 and 10 species each. Species within each community were defined by parameter values that were randomly sampled from the same distributions with the exception of toxin sensitivity *δ*, which was sampled from beta distributions with different means, ranging from 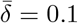 to 1. We generated a new set of 100 communities with different numbers of species and different fluctuation rates as above, and ran 100 replicate simulations for each of the 100 communities in each set. We then measured beta diversity across the 100 replicate simulations per community and final species richness (number of surviving species) over all 100 runs for the 100 communities (total: 10’000). In this model design, the 100 replicate runs represent independent “patches” without migration. Their beta diversity then indicates how different the species compositions were across all patches in a given environment (e.g. a given fluctuation rate), and we repeat the exercise 100 times with different species sampled from the same distributions to see generality of the results. A high beta diversity would then indicate that we have different community compositions in each patch, while a beta diversity of 1 would tell us that all patches have the same species composition.

In two-species communities, we obtained qualitatively similar patterns of exclusion of the fittest to those in the species interaction analysis (compare column A in Fig. 6 with Fig. 3C), suggesting that our results in Fig. 3C are unlikely to be specific to the choice of species parameter values in Table A.1. Beta diversity changes over the environmental switching rate similarly to the probability of exclusion of the fittest for four out of the five tested mean toxin sensitivities (columns A and B in Fig. 6, see also Fig. A.15): both monotonically decrease (mean toxin sensitivity 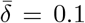 or 1.0), or non-monotonically change with maximum 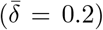 or minimum 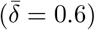 values at intermediate switching rates. This similarity can be explained as follows: ignoring extinction of both species, a small probability of exclusion of the fittest indicates that the stronger species 1 fixates in most simulations, a homogeneous outcome with small beta diversity. In contrast, when the exclusion of the fittest is more likely, the weaker species 2 is more likely to fixate, leading to significant heterogeneity in the simulation results and large beta diversity (species 2 survives alone in a fraction of the runs, and species 1 in the rest). At one mean toxin sensitivity 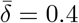, the patterns of the probability of exclusion of the fittest and beta diversity over the switching rate do not match. At this toxin sensitivity, beta diversity remains high at low switching rates (*ν* = 10^*-*5^, 10^*-*4^, 10^*-*3^) because both species go extinct in 50% of the runs but they can coexist in about 7%, as illustrated by our measure of final species richness (column C in Fig. 6). Beta diversity ignores the cases of both-species extinction but increases in cases of coexistence, see Eq (A.6). Overall, looking at final species richness (column C in Fig. 6), we see that complete extinction is more likely to occur as sensitivity increases. At the highest toxin sensitivity, the only way a species can survive is if the switching rate is really low and they can benefit from abundant resources for a long time, while at lower toxin sensitivity, complete extinctions only occur at low switching rates, because there is long term exposure to scarce resources.

**Figure 6:**
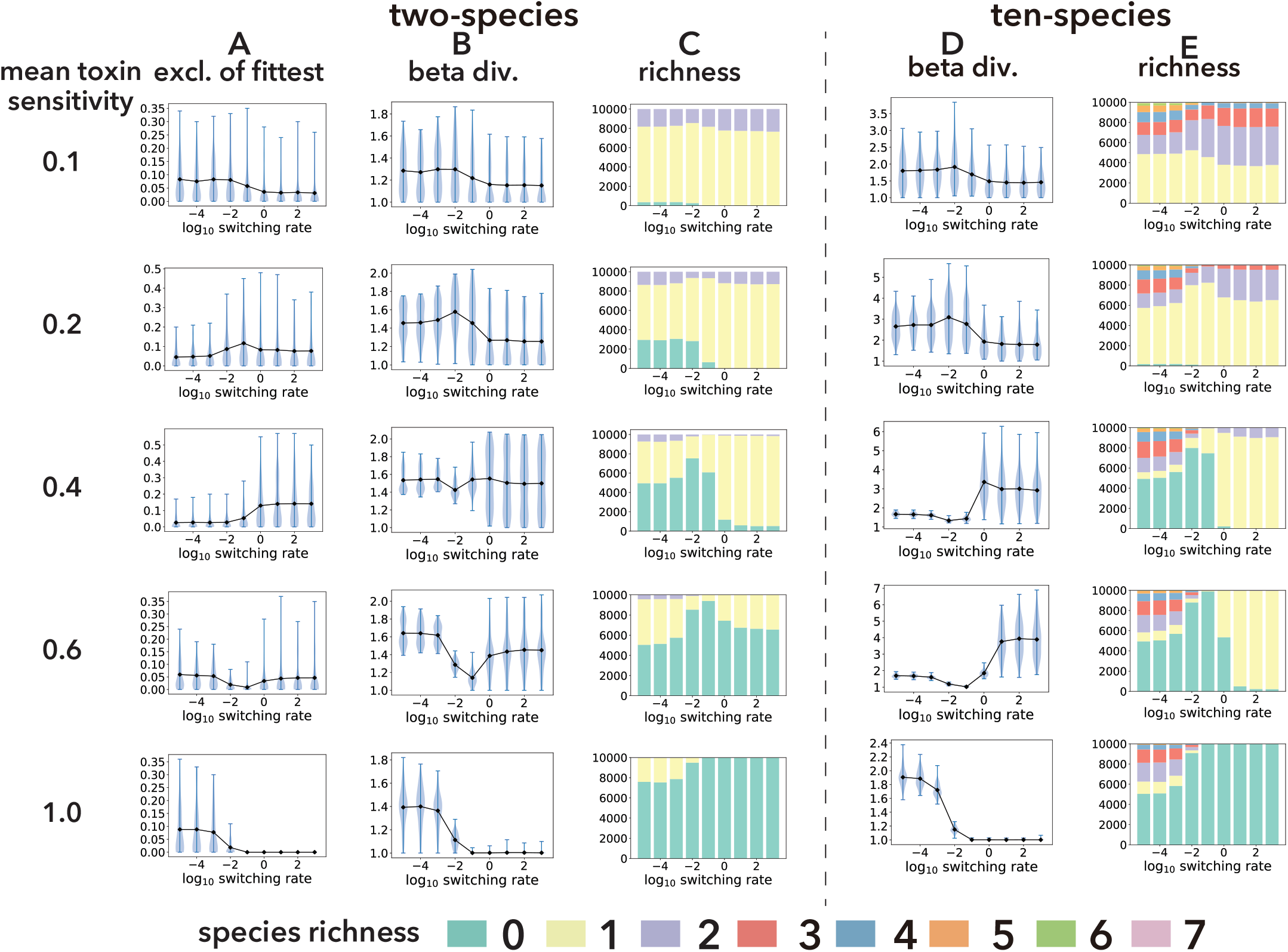
Comparison of exclusion of the fittest and diversity. Probabilities of exclusion of the fittest (column A), beta diversities (column B), and species richness (column C) over the switching rate and the mean toxin sensitivities in two-species communities. Here, competitive exclusion refers the event where the slower-growing species excludes the faster-growing species. Columns D and E show the beta diversity and species richness in ten-species communities, respectively. In the plots of probability of exclusion of the fittest and beta diversity, the black lines show the means and blue areas represent the probability distributions calculated from 10’000 simulations (100 beta diversity and each of them from 100 replicate runs). Each color in the species richness plots represents the proportion of 10’000 runs where at the end of the run there were that number of species surviving. Each row represents the mean toxin sensitivity of communities in those runs.

In communities with ten species (see Figs. A.18 and A.19 for intermediate community sizes), we observe similar patterns between beta diversity (column D in Fig. 6) and the probability of exclusion of the fittest (column A in Fig. 6) Studying interactions between species pairs can therefore predict the behavior of a ten-species community. To explore whether it matters which two species one selects for the interaction analysis, we next repeatedly sub-sampled species pairs from each ten-species community and compared the exclusion of the fittest in the pairs with the beta diversity of the whole community (Fig. A.16). Naturally, the more species pairs one samples, the more accurately we can predict the pattern of beta diversity, but this accuracy appears to saturate at around five species pairs (Appendix 7, Fig. A.17), which is approximately 11% of all possible species pairs. In addition, large beta diversity does not necessarily reflect a large variation in species richness; large beta diversity with small species richness (see mean toxin sensitivity 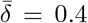 or 0.6 at switching rate *ν ≥* 10^0^ in columns D and E of Fig. 6) indicate that different species fixate in each run, which also supports the observed relationship between the exclusion of the fittest and beta diversity. In sum, estimating the probability of exclusion of the fittest between a few randomly selected species pairs (section 3.3) is a good predictor for the beta diversity of larger communities under those same environmental conditions (see Appendix 7 for detailed discussion). This similarity is not coincidental: as for the probability of exclusion of the fittest, beta diversity is also maximized when DN is the strongest, such that different species survive in each patch.

## 4 Discussion

Understanding how species diversity in microbial communities arises and is maintained is a central question in microbial ecology and evolution. While many theoretical and experimental studies have addressed this question in static environments, community diversity is expected to respond to fluctuations between benign and harsh environmental conditions, which can alter the abundance of different species and the interactions between them (see e.g., Rodríguez-Verdugo et al. (2019)). Strong drops in population sizes caused by harsh conditions can increase the strength of DN, which, coupled with EFs may lead to non-trivial outcomes (Wienand et al., 2017, 2018; West and Mobilia, 2020; Taitelbaum et al., 2020). Here we have analysed a mathematical model representing a biological scenario such as a gut or soil microbial community that experiences fluctuations between benign and harsh conditions, such as feast and famine, or drought and rain. The model shows how community diversity – mediated by inter-specific interactions and DN – changes with environmental fluctuation rates.

Our study is centred on two main findings. First, we show that the rate at which resource supplies switch changes the ability of a slower-growing species to drive a fast-growing one extinct (Fig. 3). While the fitter species will never be excluded in the absence of DN, in a fluctuating environment with DN, harsh conditions (e.g. scarce resources) strengthen DN to sometimes drive species extinction in spite of their greater fitness. We see this as a form of ecological drift, wherein selection by the environment is not strong enough to maintain the fittest species and thus the fittest species may go extinct due to DN. By changing the length of time spent in the harsh environment, the environmental switching rate affects such competitive exclusion. In addition, the species’ ability to withstand environmental harshness can also strengthen DN and lead to stronger ecological drift. We confirmed the generality of these results by exploring various forms of EFs (e.g., asymmetric switching environments and cyclically changing environments in Appendix 5). When we consider many replicate communities with identical initial species compositions, the increased stochasticity resulting from DN means that which species go extinct under these harsh conditions is less dependent on their relative competitiveness, and species composition will be more random in each replicate, leading to greater beta diversity. Recent studies corroborate this finding: smaller communities show larger varieties in species composition due to DN (Gilbert and Levine, 2017), while species composition is robust against EFs when species are insensitive to them (Dedrick et al., 2021).

Although we now understand that beta diversity increases when the probability of exclusion of the fittest is high, precisely when such ecological drift is maximized will be difficult to predict in practice, as it is a function of multiple factors: the form of the EFs (Appendix 3 and Appendix 5), the magnitude of EFs (Appendix 4), the rate of EFs and the sensitivity of species to environmental harshness. Yet, our simulation approach allows us to investigate these different effects efficiently.

Our second main finding is that a good way to predict how beta diversity will respond to EFs is to measure the probability of exclusion of the fittest across fluctuation rates in few pairs of species randomly sampled from a focal community and use it as an indicator for how the beta diversity of the whole community will behave (Fig. 6, Appendix 7). In two species communities, explaining the similarity between the probability of exclusion of the fittest and beta diversity is straightforward: the latter accounts for the probabilities that both or either of the two species persist. We verified that this similarity also holds in neutral scenarios (i.e., two species are identical except for their labels and thus the fittest species is arbitrarily chosen, Fig. A.21). In larger communities, the similarity is not straightforward but our first main finding helps to understand this result. EFs affecting DN strength could exclude species that would persist in a community without noise. If the “fittest” species goes extinct with some probability due to DN, community compositions at time *σ*_*end*_ are heterogeneous and beta diversity increases. Therefore, our two findings emphasize the importance of coupling EFs and DN: competition results can differ from those in static environments and this affects beta diversity.

This brings us to a hypothesis that has been debated at length in ecology: the intermediate disturbance hypothesis (Connell, 1978; Grime, 1973), which states that intermediate intensity and frequency of disturbance maximize species diversity. Fox (2013) argues that the intermediate disturbance hypothesis should be abandoned because many examples disagree with it (Mackey and Currie, 2001; Miller et al., 2011). In our model, fluctuations in resource and toxin can be regarded as disturbances. In agreement with Mackey and Currie (2001) and Miller et al. (2011) then, an intermediate intensity (i.e., toxin sensitivity) or disturbance frequency (environmental switching rate) does not always maximize beta diversity: our analysis shows that intermediate frequencies of disturbance maximize beta diversity only when mean toxin sensitivity is within a certain range. Mean toxin sensitivities at the two thresholds of this range show that beta diversity monotonically decreases or increases over the switching rate. These thresholds depend on scenarios of environmental switching and amounts of resource supplies because these parameters change the probability of competitive exclusion (see Appendix 3 and Appendix 4). High beta diversity at intermediate disturbances is then a consequence of a change in environmental conditions and not expected to apply generally.

The relationship between EFs and species diversity is also an important question in the modern coexistence theory, which predicts that fluctuations will affect species coexistence by changing species growth rates when rare (Chesson, 2000a,b; Barabás et al., 2018; Ellner et al., 2019; Letten et al., 2018a). Compared to the approach taken in this study – where we ask how many and which species persist at the end of a long but fixed time frame (i.e., for a quasi-stationary distribution, Fig. 6) – the modern coexistence theory allows one to analyze whether or for how long a set of species will all coexist (Schreiber et al., 2020). An interesting future direction would be to apply the modern coexistence theory to investigate how environmental fluctuation rates and toxin sensitivities affect the duration of all-species coexistence. This approach would help to propose biological mechanisms behind species coexistence in our setup.

Of course, our model makes some simplifying assumptions and has some limitations. First, we used arbitrary time units, which in practice can be considered to be hours, corresponding to typical bacterial growth rates in relevant experiments (Novick and Szilard, 1950; Lin et al., 2002; Zhao and Lin, 2003). This implies that species interactions and beta diversity will vary when environmental switching ranges from hourly (*ν* = 10^0^) to about once every four days (*ν* = 10^*–*2^) on average, which is shorter than in some experimental studies (Benneir and Lenski, 1999; Rodríguez-Verdugo et al., 2019; Chen and Zhang, 2020) but not impractical. That said, under this assumption, changing environmental switching from a daily to an hourly scale, for example, would show different species compositions or diversity.

Second, our model focuses on competitive exclusion but other types of interactions can also affect diversity (Rodríguez-Verdugo et al., 2019). Positive interactions between pairs of species (e.g., cross-feeding), for example, might increase alpha and gamma diversities, because such interactions enable species to coexist (Sun et al., 2019). This could result in an increase in beta diversity because the extinction of one species increases its partner species’ extinction probability (Dunn et al., 2009; Goldberg and Friedman, 2020).

Third, one can consider more complex species-resource (and species-toxin) interaction functions. For example, species’ growth rates can be limited by the resource with the smallest amount when the resources are complementary (León and Tumpson, 1975). In addition, absorption rates of toxin might correlate with species’ growth rate if the toxin targets cell metabolism. These functional forms may enable more species to coexist, collapsing the similarity between beta diversity and the probability of exclusion of the fittest. Our study would also be more general if species could change their growth rates depending on resource concentrations (Nguyen et al., 2020a) or types (Balakrishnan et al., 2021). Introducing such plasticity would affect species’ extinction probability and diversity. One could also consider building species-compounds interaction networks. In the current manuscript, we assume that each resource (toxin) has a positive (negative) effect on each species. However, we know that some compounds, (e.g., pesticides) can be resources for some species but toxic to others (Muturi et al., 2017), and other compounds (e.g., that affect pH (Ratzke et al., 2020) and osmolarity (Larsen, 1986; Oren, 2008)) can have either positive or negative effects on growth depending on their concentrations. The way in which species and compounds interact could affect exclusion of the fittest and species diversity.

Although we focused on beta diversity to measure the heterogeneity of communities in this manuscript, other metrics of species diversity could be considered. For example, Fung et al. (2015) and Kalyuzhny et al. (2015) analyze how demographic noise and environmental fluctuations affect species abundance distributions (SAD). Grilli (2020) instead calculate the mean abundance distribution (MAD), which is defined as the distribution of mean species abundances over communities and follows a log-normal distribution. While SAD characterizes species diversity within a single community, it does not explain the heterogeneity of species compositions across communities, which is what we can capture with beta diversity. Combining multiple SADs or probability distributions of SADs is possible, but would not be as easy to analyze. Similarly, MAD ignores the variation of species’ abundances across communities and may therefore not capture the heterogeneity of communities caused by demographic noise and environmental fluctuations.

Finally, our community analysis considers up to ten microbial species, which is orders of magnitude below the size of natural microbial communities, according to genomic sampling (Gans, 2005; Roesch et al., 2007). However, it may also be reasonable to assume that species live in structured environments where they cannot possibly interact with more than a handful of other genotypes (Tecon et al., 2019). This suggests that a 10-species community may already be biologically meaningful.

In conclusion, the time scale of environmental fluctuations changes the importance of species fitness for survival and thus community beta diversity. This occurs in our model when EFs affect the strength of DN, leading to the occasional exclusion of strong species. Predicting how the strength of DN changes is not simple because it is affected by both environmental and species’ parameters (resource and/or toxin supplies and toxin sensitivities in our model). This may be one explanation as to why the intermediate disturbance hypothesis does not always hold, but rather there are many relations between diversity and disturbance (Mackey and Currie, 2001; Miller et al., 2011). Nevertheless, we found similarities between how competitive exclusion plays out between species pairs and beta diversity at the community level. In the event that we would like to predict how the diversity of a given ecosystem, such as a soil community or a bioremediation ecosystem, responds to environmental fluctuations, it may be sufficient to isolate a few culturable species and analyze their interactions over different fluctuation rates. This approach promises to greatly facilitate our ability to study large and complex natural communities and their response to harsh conditions.

## Author Contributions

SS, MM and SM designed the study, SS performed simulations, analyzed data, and wrote the first draft, and all authors contributed to revisions.

## Acknowledgement

We thank four anonymous referees for their helpful comments in the earlier version of the manuscript.

## Data accessibility

The programming codes and simulation data in csv files for this manuscript are available in Github.

## References

Allen, L. J. S. An Introduction to Stochastic Processes with Applications to Biology. Chapman and Hall/CRC, New York, 2nd edition, 2010. ISBN 9780429184604. doi: 10.1201/b12537. URL https://www.taylorfrancis.com/books/9781439894682.

Amarasekare, P. The evolution of coexistence theory. Theoretical Population Biology, 133:49–51, 2019. ISSN 10960325. doi: 10.1016/j.tpb.2019.09.005. URL https://doi.org/10.1016/j.tpb.2019.09.005.

Assaf, M. and Meerson, B. WKB theory of large deviations in stochastic populations. Journal of Physics A: Mathematical and Theoretical, 50(26):263001, 6 2017. ISSN 1751-8113. doi: 10.1088/1751-8121/aa669a. URL http://dx.doi.org/10.1088/1751-8121/aa669a.

Balakrishnan, R., Hwa, T., and Cremer, J. Suboptimal proteome allocation during changing environments constrains bacterial response and growth recovery. bioRxiv, 2021. doi: https://doi.org/10.1101/2021.04.28.441780.

Barabás, G., D’Andrea, R., and Stump, S. M. Chesson’s coexistence theory. Ecological Monographs, 0(0):1–27, 2018. ISSN 00129615. doi: 10.1002/ecm.1302. URL http://doi.wiley.com/10.1002/ecm.1302.

Bena, I. Dichotomous Markov noise: Exact results for out-of-equilibrium systems. A review. International Journal of Modern Physics B, 20(20):2825–2888, 6 2006. ISSN 02179792. doi: 10.1142/S0217979206034881. URL http://arxiv.org/abs/cond-mat/0606116.

Benneir, A. F. and Lenski, R. E. Experimental evolution and its role in evolutionary physiology. American Zoologist, 39(2):346–362, 1999. ISSN 00031569. doi: 10.1093/icb/39.2.346.

Butler, S. and O’Dwyer, J. Cooperation and Stability for Complex Systems in Resource-Limited Environments. Theoretical Ecology, 13:239–250, 2020. doi: 10.1101/514018. URL https://doi.org/10.1007/s12080-019-00447-5.

Butler, S. and O’Dwyer, J. P. Stability criteria for complex microbial communities. Nature Communications, 9(1):2970, 12 2018. ISSN 2041-1723. doi: 10.1038/s41467-018-05308-z. URL http://www.nature.com/articles/s41467-018-05308-z.

Cangelosi, G. A. and Meschke, J. S. Dead or Alive: Molecular Assessment of Microbial Viability. Applied and Environmental Microbiology, 80(19):5884–5891, 10 2014. ISSN 0099-2240. doi: 10.1128/AEM.01763-14. URL https://journals.asm.org/doi/10.1128/AEM.01763-14.

Chao, A., Chiu, C. H., Hsieh, T. C., and Inouye, B. D. Proposing a resolution to debates on diversity partitioning. Ecology, 93(9):2037–2051, 9 2012. ISSN 00129658. doi: 10.1890/11-1817.1. URL http://doi.wiley.com/10.1890/11-1817.1.

Chen, P. and Zhang, J. Antagonistic pleiotropy conceals molecular adaptations in changing environments. Nature Ecology & Evolution, 2 2020. ISSN 2397-334X. doi: 10.1038/s41559-020-1107-8. URL http://www.nature.com/articles/s41559-020-1107-8.

Chesson, P. Multispecies Competition in Variable Environments. Theoretical Population Biology, 45:227–276, 1994.

Chesson, P. Mechanisms of Maintenance of Species Diversity. Annual Review of Ecology and Systematics, 31(1):343–366, 11 2000a. ISSN 0066-4162. doi: 10.1146/annurev.ecolsys.31.1.343. URL http://www.annualreviews.org/doi/10.1146/annurev.ecolsys.31.1.343.

Chesson, P. General theory of competitive coexistence in spatially-varying environments. Theoretical Population Biology, 58(3):211–237, 2000b. ISSN 00405809. doi: 10.1006/tpbi.2000.1486.

Chisholm, R. A., Condit, R., Rahman, K. A., Baker, P. J., Bunyavejchewin, S., Chen, Y.-Y., Chuyong, G., Dattaraja, H. S., Davies, S., Ewango, C. E. N., Gunatilleke, C. V. S., Nimal Gunatilleke, I. A. U., Hubbell, S., Kenfack, D., Kiratiprayoon, S., Lin, Y., Makana, J.-R., Pongpattananurak, N., Pulla, S., Punchi-Manage, R., Sukumar, R., Su, S.-H., Sun, I.-F., Suresh, H. S., Tan, S., Thomas, D., and Yap, S. Temporal variability of forest communities: empirical estimates of population change in 4000 tree species. Ecology Letters, 17(7): 855–865, 7 2014. ISSN 1461023X. doi: 10.1111/ele.12296. URL http://doi.wiley.com/10.1111/ele.12296.

Cignarella, F., Cantoni, C., Ghezzi, L., Salter, A., Dorsett, Y., Chen, L., Phillips, D., Weinstock, G. M., Fontana, L., Cross, A. H., Zhou, Y., and Piccio, L. Intermittent Fasting Confers Protection in CNS Autoimmunity by Altering the Gut Microbiota. Cell Metabolism, 27(6):1222–1235, 2018. ISSN 19327420. doi: 10.1016/j.cmet.2018.05.006. URL https://doi.org/10.1016/j.cmet.2018.05.006.

Connell, J. H. Diversity in Tropical Rain Forests and Coral Reefs. Science, 199(4335):1302–1309, 1978.

Coyte, K. Z., Schluter, J., and Foster, K. R. The ecology of the microbiome: Networks, competition, and stability. Science, 350(6261):663–666, 2015. ISSN 0036-8075. doi: 10.1126/science.aad2602. URL http://www.sciencemag.org/cgi/doi/10.1126/science.aad2602.

Davenport, E. R., Mizrahi-Man, O., Michelini, K., Barreiro, L. B., Ober, C., and Gilad, Y. Seasonal variation in human gut microbiome composition. PLoS ONE, 9(3), 2014. ISSN 19326203. doi: 10.1371/journal.pone.0090731.

Dedrick, S., Akbari, M. J., Dyckman, S. K., Zhao, N., Liu, Y.-Y., and Momeni, B. Impact of Temporal pH Fluctuations on the Coexistence of Nasal Bacteria in an in silico Community. Frontiers in Microbiology, 12 (February):1–12, 2021. ISSN 1664302X. doi: 10.3389/fmicb.2021.613109.

Dunn, R. R., Harris, N. C., Colwell, R. K., Koh, L. P., and Sodhi, N. S. The sixth mass coextinction: are most endangered species parasites and mutualists? Proceedings of the Royal Society B: Biological Sciences, 276(1670):3037–3045, 9 2009. ISSN 0962-8452. doi: 10.1098/rspb.2009.0413. URL https://royalsocietypublishing.org/doi/10.1098/rspb.2009.0413.

Ellner, S. P., Snyder, R. E., Adler, P. B., and Hooker, G. An expanded modern coexistence theory for empirical applications. Ecology Letters, 22(1):3–18, 1 2019. ISSN 1461-023X. doi: 10.1111/ele.13159. URL https://onlinelibrary.wiley.com/doi/abs/10.1111/ele.13159.

Engen, S. and Lande, R. Population Dynamic Models Generating Species Abundance Distributions of the Gamma Type. Journal of Theoretical Biology, 178(3):325–331, 2 1996. ISSN 00225193. doi: 10.1006/jtbi.1996.0028. URL https://linkinghub.elsevier.com/retrieve/pii/S0022519396900284.

Ewens, W. J. Mathematical population genetics. Springer, 2004. doi: 10.1007/978-0-387-21822-9.

Fox, J. W. The intermediate disturbance hypothesis should be abandoned. Trends in Ecology & Evolution, 28 (2):86–92, 2 2013. ISSN 01695347. doi: 10.1016/j.tree.2012.08.014. URL https://linkinghub.elsevier.com/retrieve/pii/S0169534712002091.

Fung, T., Villain, L., and Chisholm, R. A. Analytical formulae for computing dominance from species-abundance distributions. Journal of Theoretical Biology, 386:147–158, 2015. ISSN 10958541. doi: 10.1016/j.jtbi.2015.09.011. URL http://dx.doi.org/10.1016/j.jtbi.2015.09.011.

Gans, J. Computational Improvements Reveal Great Bacterial Diversity and High Metal Toxicity in Soil. Science, 309(5739):1387–1390, 8 2005. ISSN 0036-8075. doi: 10.1126/science.1112665. URL https://www.sciencemag.org/lookup/doi/10.1126/science.1112665.

Gilbert, B. and Levine, J. M. Ecological drift and the distribution of species diversity. Proceedings of the Royal Society B: Biological Sciences, 284(1855):20170507, 5 2017. ISSN 0962-8452. doi: 10.1098/rspb.2017.0507. URL https://royalsocietypublishing.org/doi/10.1098/rspb.2017.0507.

Gillespie, D. T. Exact stochastic simulation of coupled chemical reactions. Journal of Physical Chemistry, 81 (25):2340–2361, 1977. ISSN 00223654. doi: 10.1021/j100540a008.

Goldberg, Y. and Friedman, J. Positive interactions within and between populations decrease the likelihood of evolutionary rescue. bioRxiv, 2020. ISSN 26928205. doi: 10.1101/2020.08.06.239608. URL http://dx.doi.org/10.1371/journal.pcbi.1008732.

Grilli, J. Macroecological laws describe variation and diversity in microbial communities. Nature Communications, 11(1):1–11, 2020. ISSN 20411723. doi: 10.1038/s41467-020-18529-y. URL http://dx.doi.org/10.1038/s41467-020-18529-y.

Grime, J. P. Competitive Exclusion in Herbaceous Vegetation. Nature, 242(5396):344–347, 3 1973. ISSN 0028-0836. doi: 10.1038/242344a0. URL http://www.nature.com/doifinder/10.1038/242344a0.

Grime, J. P. Evidence for the Existence of Three Primary Strategies in Plants and Its Relevance to Ecological and Evolutionary Theory. The American Naturalist, 111(982):1169–1194, 1977. ISSN 0003-0147. doi: 10.1086/283244.

Guittar, J., Koffel, T., Shade, A., Klausmeier, C. A., and Litchman, E. Resource Competition and Host Feedbacks Underlie Regime Shifts in Gut Microbiota. The American Naturalist, 198(1):000–000, 5 2021. ISSN 0003-0147. doi: 10.1086/714527. URL https://www.journals.uchicago.edu/doi/10.1086/714527.

Hengge-Aronis, R. Survival of hunger and stress: The role of rpoS in early stationary phase gene regulation in E. coli. Cell, 72(2):165–168, 1 1993. ISSN 00928674. doi: 10.1016/0092-8674(93)90655-A. URL https://linkinghub.elsevier.com/retrieve/pii/009286749390655A.

Himeoka, Y. and Mitarai, N. Dynamics of bacterial populations under the feast-famine cycles. arXiv, pages 1–39, 10 2019. URL http://arxiv.org/abs/1910.05673.

Hoek, T. A., Axelrod, K., Biancalani, T., Yurtsev, E. A., Liu, J., and Gore, J. Resource Availability Modulates the Cooperative and Competitive Nature of a Microbial Cross-Feeding Mutualism. PLOS Biology, 14(8): e1002540. 8 2016. ISSN 1545-7885. doi: 10.1371/journal.pbio.1002540. URL http://dx.plos.org/10.1371/journal.pbio.1002540.

Horsthemke, W. and Lefever, R. Noise-Induced Transitions, volume 15 of Springer Series in Synergetics. Springer Berlin Heidelberg, 2nd edition, 4 2006. ISBN 978-3-540-11359-1. doi: 10.1007/3-540-36852-3. URL http://link.springer.com/10.1007/3-540-36852-3.

Huang, Q., Parshotam, L., Wang, H., Bampfylde, C., and Lewis, M. A. A model for the impact of contaminants on fish population dynamics. Journal of Theoretical Biology, 334:71–79, 2013. ISSN 00225193. doi: 10.1016/j.jtbi.2013.05.018. URL http://dx.doi.org/10.1016/j.jtbi.2013.05.018.

Jost, L. PARTITIONING DIVERSITY INTO INDEPENDENT ALPHA AND BETA COMPONENTS. Ecology, 88(10):2427–2439, 10 2007. ISSN 0012-9658. doi: 10.1890/06-1736.1. URL http://doi.wiley.com/10.1890/06-1736.1.

Kalyuzhny, M., Kadmon, R., and Shnerb, N. M. A neutral theory with environmental stochasticity explains static and dynamic properties of ecological communities. Ecology Letters, 18(6):572–580, 6 2015. ISSN 1461023X. doi: 10.1111/ele.12439. URL http://doi.wiley.com/10.1111/ele.12439.

Kamenev, A., Meerson, B., and Shklovskii, B. How Colored Environmental Noise Affects Population Extinction. Physical Review Letters, 101(26):268103, 12 2008. ISSN 0031-9007. doi: 10.1103/PhysRevLett.101.268103. URL https://link.aps.org/doi/10.1103/PhysRevLett.101.268103.

Larsen, H. Halophilic and halotolerant microorganisms-an overview and historical perspective. FEMS Microbiology Reviews, 39(April):3–7, 1986.

Leigh, E. G. The average lifetime of a population in a varying environment. Journal of Theoretical Biology, 90(2):213–239, 5 1981. ISSN 00225193. doi: 10.1016/0022-5193(81)90044-8. URL https://linkinghub.elsevier.com/retrieve/pii/0022519381900448.

León, J. A. and Tumpson, D. B. Competition between two species for two complementary or substitutable resources. Journal of Theoretical Biology, 50(1):185–201, 1975. ISSN 10958541. doi: 10.1016/0022-5193(75)90032-6.

Letten, A. D., Dhami, M. K., Ke, P. J., and Fukami, T. Species coexistence through simultaneous fluctuation-dependent mechanisms. Proceedings of the National Academy of Sciences of the United States of America, 115(26):6745–6750, 2018a. ISSN 10916490. doi: 10.1073/pnas.1801846115.

Letten, A. D., Dhami, M. K., Ke, P. J., and Fukami, T. Species coexistence through simultaneous fluctuation-dependent mechanisms. Proceedings of the National Academy of Sciences of the United States of America, 115(26):6745–6750, 2018b. ISSN 10916490. doi: 10.1073/pnas.1801846115.

Li, G., Xie, C., Lu, S., Nichols, R. G., Tian, Y., Li, L., Patel, D., Ma, Y., Brocker, C. N., Yan, T., Krausz, K. W., Xiang, R., Gavrilova, O., Patterson, A. D., and Gonzalez, F. J. Intermittent Fasting Promotes White Adipose Browning and Decreases Obesity by Shaping the Gut Microbiota. Cell Metabolism, 26(4):672–685, 2017. ISSN 19327420. doi: 10.1016/j.cmet.2017.08.019. URL http://dx.doi.org/10.1016/j.cmet.2017.08.019.

Lin, Y. H., Bayrock, D. P., and Ingledew, W. M. Evaluation of Saccharomyces cerevisiae grown in a multistage chemostat environment under increasing levels of glucose. Biotechnology Letters, 24(6):449–453, 2002. doi: 10.1023/A:1014501125355.

Mackey, R. L. and Currie, D. J. The diversity-disturbance relationship: Is it generally strong and peaked? Ecology, 82(12):3479–3492, 2001. ISSN 00129658. doi: 10.1890/0012-9658(2001)082[3479:TDDRII]2.0.CO;2.

Marsland III, R., Cui, W., Goldford, J., Sanchez, A., Korolev, K., and Mehta, P. Available energy fluxes drive a transition in the diversity, stability, and functional structure of microbial communities. PLOS Computational Biology, 15(2):e1006793. 2 2019. doi: 10.1371/journal.pcbi.1006793. URL http://dx.plos.org/10.1371/journal.pcbi.1006793.

Merritt, J. and Kuehn, S. Frequency-and Amplitude-Dependent Microbial Population Dynamics during Cycles of Feast and Famine. Physical Review Letters, 121(9):098101, 8 2018. ISSN 0031-9007. doi: 10.1103/PhysRevLett.121.098101. URL https://link.aps.org/doi/10.1103/PhysRevLett.121.098101.

Miller, A. D., Roxburgh, S. H., and Shea, K. How frequency and intensity shape diversity-disturbance relationships. Proceedings of the National Academy of Sciences of the United States of America, 108(14):5643–5648, 2011. ISSN 10916490. doi: 10.1073/pnas.1018594108.

Molinero, N., Ruiz, L., Sánchez, B., Margolles, A., and Delgado, S. Intestinal bacteria interplay with bile and cholesterol metabolism: Implications on host physiology. Frontiers in Physiology, 10(MAR):1–10, 2019. ISSN 1664042X. doi: 10.3389/fphys.2019.00185.

Mougi, A. and Kondoh, M. Diversity of Interaction Types and Ecological Community Stability. Science, 337 (6092):349–351, 2012. ISSN 0036-8075. doi: 10.1126/science.1220529. URL http://www.sciencemag.org/cgi/doi/10.1126/science.1220529.

Muturi, E. J., Donthu, R. K., Fields, C. J., Moise, I. K., and Kim, C.-H. Effect of pesticides on microbial communities in container aquatic habitats. Scientific Reports, 7(1):44565, 4 2017. ISSN 2045-2322. doi: 10.1038/srep44565. URL http://www.nature.com/articles/srep44565.

Nguyen, J., Fernandez, V., Pontrelli, S., Sauer, U., Ackermann, M., and Stocker, R. A distinct growth physiology enhances bacterial growth under rapid nutrient fluctuations. bioRxiv, page 2020.08.18.256529, 2020a. URL https://www.biorxiv.org/content/10.1101/2020.08.18.256529v1.

Nguyen, J., Lara-Gutiérrez, J., and Stocker, R. Environmental fluctuations and their effects on microbial communities, populations, and individuals. FEMS Microbiology Reviews, 12 2020b. ISSN 0168-6445. doi: 10.1093/femsre/fuaa068. URL https://academic.oup.com/femsre/advance-article/doi/10.1093/femsre/fuaa068/6041721.

Novick, A. and Szilard, L. Experiments with the Chemostat on Spontaneous Mutations of Bacteria. Proceedings of the National Academy of Sciences, 36(12):708–719, 12 1950. ISSN 0027-8424. doi: 10.1073/pnas.36.12.708. URL http://www.pnas.org/cgi/doi/10.1073/pnas.36.12.708.

Novozhilov, A. S., Karev, G. P., and Koonin, E. V. Biological applications of the theory of birth-and-death processes. Briefings in Bioinformatics, 7(1):70–85, 3 2006. ISSN 1467-5463. doi: 10.1093/bib/bbk006. URL https://academic.oup.com/bib/article/7/1/70/263777.

Oren, A. Microbial life at high salt concentrations: Phylogenetic and metabolic diversity. Saline Systems, 4(1): 1–13, 2008. ISSN 17461448. doi: 10.1186/1746-1448-4-2.

Pérez, S., Eichhorn, P., and Aga, D. S. Evaluating the biodegradability of sulfamethazine, sulfamethoxazole, sulfathiazole, and trimethoprim at different stages of sewage treatment. Environmental Toxicology and Chemistry, 24(6):1361, 2005. ISSN 0730-7268. doi: 10.1897/04-211R.1. URL http://doi.wiley.com/10.1897/04-211R.1.

Piccardi, P., Vessman, B., and Mitri, S. Toxicity drives facilitation between 4 bacterial species. Proceedings of the National Academy of Sciences, 116(32):15979–15984, 8 2019. ISSN 0027-8424. doi: 10.1073/pnas.1906172116. URL http://www.pnas.org/lookup/doi/10.1073/pnas.1906172116.

Ratzke, C., Barrere, J., and Gore, J. Strength of species interactions determines biodiversity and stability in microbial communities. Nature Ecology and Evolution, 4(3):376–383, 2020. ISSN 2397334X. doi: 10.1038/s41559-020-1099-4. URL http://dx.doi.org/10.1038/s41559-020-1099-4.

Rodríguez-Verdugo, A., Vulin, C., and Ackermann, M. The rate of environmental fluctuations shapes ecological dynamics in a two-species microbial system. Ecology Letters, 22(5):838–846, 5 2019. ISSN 1461-023X. doi: 10.1111/ele.13241. URL http://doi.wiley.com/10.1111/ele.13241.

Roesch, L. F., Fulthorpe, R. R., Riva, A., Casella, G., Hadwin, A. K., Kent, A. D., Daroub, S. H., Camargo, F. A., Farmerie, W. G., and Triplett, E. W. Pyrosequencing enumerates and contrasts soil microbial diversity. ISME Journal, 1(4):283–290, 2007. ISSN 17517362. doi: 10.1038/ismej.2007.53.

Roughgarden, J. Theory of population genetics and evolutionary ecology : an introduction. Macmillan, New York, 1979. ISBN 0024031801.

Ruiz, L., Margolles, A., and Sánchez, B. Bile resistance mechanisms in Lactobacillus and Bifidobacterium. Frontiers in Microbiology, 4(DEC):1–8, 2013. ISSN 1664302X. doi: 10.3389/fmicb.2013.00396.

Schreiber, S., Levine, J., Godoy, O., Kraft, N., and Hart, S. Does deterministic coexistence theory matter in a finite world? bioRxiv, 2020. doi: 10.1101/290882.

Smits, S. A., Leach, J., Sonnenburg, E. D., Gonzalez, C. G., Lichtman, J. S., Reid, G., Knight, R., Manjurano, A., Changalucha, J., Elias, J. E., Dominguez-Bello, M. G., and Sonnenburg, J. L. Seasonal cycling in the gut microbiome of the Hadza hunter-gatherers of Tanzania. Science, 357(6353):802–805, 2017. ISSN 10959203. doi: 10.1126/science.aan4834.

Spalding, C., Doering, C. R., and Flierl, G. R. Resonant activation of population extinctions. Physical Review E, 96(4):042411, 10 2017. ISSN 2470-0045. doi: 10.1103/PhysRevE.96.042411. URL http://dx.doi.org/10.1103/PhysRevE.96.042411.

Srinivasan, S. and Kjelleberg, S. Cycles of famine and feast: The starvation and outgrowth strategies of a marine Vibrio. Journal of Biosciences, 23(4):501–511, 1998. ISSN 02505991. doi: 10.1007/BF02936144.

Sun, Z., Koffel, T., Stump, S. M., Grimaud, G. M., and Klausmeier, C. A. Microbial cross-feeding promotes multiple stable states and species coexistence, but also susceptibility to cheaters. Journal of Theoretical Biology, 465:63–77, 2019. ISSN 00225193. doi: 10.1016/j.jtbi.2019.01.009. URL https://linkinghub.elsevier.com/retrieve/pii/S0022519319300104.

Sunya, S., Bideaux, C., Molina-Jouve, C., and Gorret, N. Short-term dynamic behavior of Escherichia coli in response to successive glucose pulses on glucose-limited chemostat cultures. Journal of Biotechnology, 164 (4):531–542, 2013. ISSN 01681656. doi: 10.1016/j.jbiotec.2013.01.014. URL http://dx.doi.org/10.1016/j.jbiotec.2013.01.014.

Taitelbaum, A., West, R., Assaf, M., and Mobilia, M. Population Dynamics in a Changing Environment: Random versus Periodic Switching. Physical Review Letters, 125(4):048105, 7 2020. ISSN 0031-9007. doi: 10.1103/PhysRevLett.125.048105. URL https://link.aps.org/doi/10.1103/PhysRevLett.125.048105.

Tecon, R., Mitri, S., Ciccarese, D., Or, D., van der Meer, J. R., and Johnson, D. R. Bridging the Holistic-Reductionist Divide in Microbial Ecology. mSystems, 4(1):17–21, 2019. ISSN 2379-5077. doi: 10.1128/msystems.00265-18.

Thaiss, C. A., Zeevi, D., Levy, M., Zilberman-Schapira, G., Suez, J., Tengeler, A. C., Abramson, L., Katz, M. N., Korem, T., Zmora, N., Kuperman, Y., Biton, I., Gilad, S., Harmelin, A., Shapiro, H., Halpern, Z., Segal, E., and Elinav, E. Transkingdom control of microbiota diurnal oscillations promotes metabolic homeostasis. Cell, 159(3):514–529, 2014. ISSN 10974172. doi: 10.1016/j.cell.2014.09.048. URL http://dx.doi.org/10.1016/j.cell.2014.09.048.

Vasi, F., Travisano, M., and Lenski, R. E. Long-term experimental evolution in Escherichia coli. II.Changes in life-history traits during adaptation to a seasonal environment. American Naturalist, 144(3):432–456, 1994. ISSN 00030147. doi: 10.1086/285685. URL https://www.jstor.org/stable/2462954.

West, R. and Mobilia, M. Fixation properties of rock-paper-scissors games in fluctuating populations. Journalof Theoretical Biology, 491:110135, 4 2020. ISSN 00225193. doi: 10.1016/j.jtbi.2019.110135. URL http://dx.doi.org/10.1016/j.jtbi.2019.110135.

Wienand, K., Frey, E., and Mobilia, M. Evolution of a Fluctuating Population in a Randomly Switching Environment. Physical Review Letters, 119(15):158301, 10 2017. ISSN 0031-9007. doi: 10.1103/PhysRevLett.119.158301. URL https://link.aps.org/doi/10.1103/PhysRevLett.119.158301.

Wienand, K., Frey, E., and Mobilia, M. Eco-evolutionary dynamics of a population with randomly switching carrying capacity. Journal of the Royal Society Interface, 15(145):20180343, 8 2018. ISSN 17425662. doi: 10.1098/rsif.2018.0343. URL https://royalsocietypublishing.org/doi/10.1098/rsif.2018.0343.

Xavier, J. B., Picioreanu, C., and Van Loosdrecht, M. C. M. A framework for multidimensional modelling of activity and structure of multispecies biofilms. Environmental Microbiology, 7(8):1085–1103, 2005. ISSN 14622912. doi: 10.1111/j.1462-2920.2005.00787.x.

Xu, B., Mao, D., Luo, Y., and Xu, L. Sulfamethoxazole biodegradation and biotransformation in the water-sediment system of a natural river. Bioresource Technology, 102(14):7069–7076, 2011. ISSN 09608524. doi: 10.1016/j.biortech.2011.04.086. URL http://dx.doi.org/10.1016/j.biortech.2011.04.086.

Zhao, Y. and Lin, Y. H. Growth of Saccharomyces cerevisiae in a chemostat under high glucose conditions. Biotechnology Letters, 25(14):1151–1154, 2003. ISSN 01415492. doi: 10.1023/A:1024577414157.

Zuñiga, C., Li, C.-T., Yu, G., Al-Bassam, M. M., Li, T., Jiang, L., Zaramela, L. S., Guarnieri, M., Betenbaugh, M. J., and Zengler, K. Environmental stimuli drive a transition from cooperation to competition in synthetic phototrophic communities. Nature Microbiology, 4(12):2184–2191, 12 2019. ISSN 2058-5276. doi: 10.1038/s41564-019-0567-6. URL http://www.nature.com/articles/s41564-019-0567-6.

