## Appendices for "Exclusion of the fittest predicts microbial community diversity in fluctuating environments"

### Supplementary information

#### Appendix 1 Details of analysis

##### Appendix 1.1 Species interaction analysis

Here, we summarize the details of the simulations and parameter values used for species interaction analysis in the main text. In the analysis of species interactions, we used the minimal model ($N = 2$): two species compete for one resource and absorb and are killed by one toxin. We assumed that species 1 grows faster than species 2 but the other parameter values are identical (Table A.1). In each run, the initial condition is either $(r_1(0), t_1(0), s_1(0), s_2(0)) = (150, 100, 10, 0)$ or $(150, 100, 10, 10)$, where species 2 is absent or present, respectively. The initial environmental condition is $\xi = 1$ with a probability of 0.5; otherwise $\xi = -1$. Each simulation continues until at time $\sigma_{end} = 200$ when the distributions of species abundances converge to a quasi-stationary distribution and do not change for a long time. The extinction probability was estimated by running $10^5$ simulations for each condition.

In the main text, to investigate how species 2's effect on species 1 changes over the switching rate, we began by analyzing the difference in the extinction probability of species 1 in the presence/absence of species 2 as a proxy for species interactions:

$$\Delta P\left(s_1(\sigma_{end}) = 0\right) \equiv P\left(s_1(\sigma_{end}) = 0; s_2(0) = 0\right) - P\left(s_1(\sigma_{end}) = 0; s_2(0) > 0\right). \tag{A.1}$$

Then, we moved to using the probability of exclusion of the fittest (probability that species 2 excludes species 1) instead:

$$\Delta P\left(s_1(\sigma_{end}) = 0\right) = -\underbrace{P\left(s_1(\sigma_{end}) = 0, s_2(\sigma_{end}) > 0; s_2(0) > 0\right)}_{\text{exclusion of the fittest}} + \left\{ \underbrace{P\left(s_1(\sigma_{end}) = 0; s_2(0) = 0\right)}_{\text{sp 1 goes extinct in mono-culture}} - \underbrace{P\left(s_1(\sigma_{end}) = 0, s_2(\sigma_{end}) = 0; s_2(0) > 0\right)}_{\text{both species go extinct}} \right\}. \tag{A.2}$$

Although Fig.3A and C show that these two measures give similar results, Fig. A.2 confirms this conclusion as the ratio of these two measures is around 1. In other words, the second line on the right-hand-side of Eq (A.2) is ignorable. Intuitively, this is because the environment is very harsh when both species are likely to go extinct (i.e., small resource supplies and high toxin sensitivity); then species 1 is also likely to go extinct in mono-culture under such harsh environment. For this reason, the competitive exclusion probability (the first line on the right-hand-side of Eq (A.2)) changes similarly to difference in extinction probability ($\Delta P\left(s_1\left(\sigma_{end}\right) = 0\right)$) over environmental switching rates and toxin sensitivities (Fig. 3C), although they differ in their signs.

We calculated 95% of highest posterior density intervals (HDPIs) to see the uncertainty of species '1 extinction probabilities or the probability of exclusion of the fittest. As we were interested in whether species 1 goes extinct (or is excluded by species 2) or not in each simulation run, it is reasonable to assume these probabilities follow beta distributions. As a prior distribution, we assumed the following beta distribution

$$\mathrm{Beta}\,(1, 1)\,, \tag{A.3}$$

which is equivalent to a uniform distribution between 0 and 1. After running $10^5$ simulations and observing species 1's extinction (or exclusion) $X$ times, the posterior probability distribution of species 1's extinction probability (or the probability of exclusion of the fittest) is given as follows:

$$\mathrm{Beta}\left(X + 1, 10^5 - X + 1\right). \tag{A.4}$$

To calculate the 95% HDPIs, we sampled 10,000 samples from the posterior distributions and used pymc3.stats.hpd function. However, the 95% HDPIs are very small and not observable in the main text due to the large number of simulations.

#### Appendix 1.2 Landscapes of exclusion of the fittest

To understand how the switching rate and toxin sensitivity affect the probability of the exclusion of the fittest, we analyzed three cases in the absence of EFs (Fig. 5), deriving three critical toxin sensitivities where the probability of exclusion of the fittest is maximized. This analysis clarifies what happens at the two extreme switching rates. We see two critical toxin sensitivities (at 0.1 and 0.8 in Figs. 3C and D) at $\nu \to 0$ that correspond to the long time spent with either scarce or abundant resources, while at $\nu \to \infty$, where resources remain at mean abundance, there is a single critical toxin sensitivity (at 0.4 in Figs. 3C and D). As the switching rate increases from one extreme to the other, the form of the probability of exclusion of the fittest in Fig. 3C changes from bi-modal to uni-modal.

We now see that the landscape of competitive exclusion in Fig. 3C contains two "mountain ranges". The first includes two peaks corresponding to the critical toxin sensitivities under scarce and mean resources $(0.1, 0.4$, respectively). The peak at toxin sensitivity 0.1 converges to the peak at 0.4 (Fig. 3D) with increasing environmental switching. The second mountain range has a single peak corresponding to the critical toxin sensitivity under abundant resources (0.8), which vanishes by increasing the switching rate (Fig. 3D). At toxin sensitivities between the critical values under scarce and mean resource supplies ($\delta = 0.2, 0.3$), the probability of exclusion of the fittest changes in a humped shape over the switching rate (species 1's difference in extinction probability changes in a U shape, see Fig. 3B), as it passes over the first mountain range. When the toxin sensitivity is between the critical values under mean and abundant resources (e.g. $\delta = 0.6$), the probability of exclusion of the fittest instead has a "valley" over the switching rate (the difference in extinction probability changes in a humped shape, see Fig. 3B). At toxin sensitivities larger than the abundant resource critical value ($\delta > 0.8$), exclusion of the fittest is very unlikely because both species frequently go extinct (Fig. 5C).

In summary, the transition between the two extreme switching rates results in a highly rugged landscape. This means that the stochastic exclusion of the fittest species can be very high or very low at an intermediate switching rate (either in agreement or in contradiction with the intermediate disturbance hypothesis), depending on species' toxin sensitivities, our proxy for environmental harshness. This makes predicting the outcome at a given switching rate very difficult, as it is dependent on precise parameter values.

#### Appendix 1.3 Species diversity analysis

In the community diversity analysis, we changed the number of species from $N = 2$ to $N = 10$. Some parameter values were not fixed in this analysis, and we sampled them from the following probability density functions:

$$\mu_{ik} \sim \mathcal{N}\left(1, 0.1^2\right), \tag{A.5a}$$

$$\delta_{jk} \sim \text{Beta}\left(100\bar{\delta}, 100\left(1-\bar{\delta}\right)\right), \tag{A.5b}$$

$$K^r_{ik}, K^t_{jk} \sim \mathcal{N}\left(100, 10^2\right). \tag{A.5c}$$

Here, each function is uni-modal with a mean of 1.0 for $\mu_{ik}$, $\bar{\delta}$ for $\delta_{jk}$, and 100 for $K^r_{ik}, K^t_{jk}$. For $\mu_{ik}, K^r_{ik}$, and $K^t_{jk}$, the mean values are the same as in Table A.1 and they are rarely negative due to the small variances. We set the mean of $\mu_{ik}$ as 1 so that $\mu_{ik}$ is likely to be larger than $\delta_{jk}$ unless $\bar{\delta} = 0.99$: species would easily go extinct when $\mu_{ik} < \delta_{jk}$. The means of $K^r_{ik}$ and $K^t_{jk}$ are chosen so that amounts of resource and/or toxin inflows are larger (smaller) than these values under the abundant (scarce) supply condition. We expected that the growth and/or death rates of species would largely change depending on the environmental conditions with this setting. We sampled $\delta_{jk}$ from a beta distribution so that $0 \leq \delta_{jk} \leq 1$. $\delta_{jk}$ should be non-negative by definition and not be larger than 1 because a large $\delta_{jk}$ is likely to drive species $k$ extinct. Beta distribution satisfies these requirements regardless value of mean $\bar{\delta}$. We systematically vary the value of the mean toxin sensitivity $\bar{\delta}$ to be $\bar{\delta} = 0.1, 0.2, 0.4, 0.6$ or 0.99 in each set of simulations. We used $\bar{\delta} = 0.99$ instead of $\bar{\delta} = 1.0$ because the beta distribution did not generate different values of $\delta_{jk}$ with $\bar{\delta} = 1.0$. For each number of species $N$ (2, 4, 6, 8 or 10) and $\bar{\delta}$, 100 sets of parameter values are sampled. With each parameter set, we performed the simulation 100 times until $\sigma_{end} = 200$ at each value of the switching rate $\nu$. Then, we calculated the species richness (the number of surviving species) and beta diversity.

In each run, initial resource amounts, species abundances, and toxin amounts are given by $(r_i(0), t_j(0), s_k(0)) = (150, 150, 10)$ for any $i, j, k$. As an environmental switching scenario, we chose scenario 1 (Table 1) with $R_i^+ = 200$, $R_i^- = 50$, and $\langle T_k \rangle = 125$ for any $i$ and $k$. The initial environmental condition is $\xi = 1$ with probability of 0.5; otherwise $\xi = -1$.

Then, at a quasi-stationary distribution ($\sigma_{end}$), we evaluated beta diversity and species richness. Beta diversity is calculated as follows:

$${}^1D_\beta\left(\sigma_{end}\right) \equiv \frac{{}^1D_\gamma\left(\sigma_{end}\right)}{{}^1D_\alpha\left(\sigma_{end}\right)}, \tag{A.6}$$

with alpha and gamma diversities defined as below:

$$
{}^{1}D_{\alpha}\left(\sigma_{end}\right) \equiv \exp\left(-\sum_{l=1}^{100}\sum_{k=1}^{N} w_l p_{lk}\left(\sigma_{end}\right)\ln p_{ln}\left(\sigma_{end}\right)\right), \tag{A.7}
$$

$$
{}^{1}D_{\gamma}\left(\sigma_{end}\right) \equiv \exp\left(-\sum_{k=1}^{N} \bar{p}_k \ln \bar{p}_k\left(\sigma_{end}\right)\right). \tag{A.8}
$$

$w_l$ is a weight for community $l$ calculated by size of community $l$ (sum of species abundances in community $l$ relative to the sum of community sizes over $l$), $p_{lk}$ is the relative abundance of species $k$ in community $l$ (i.e., in community $l$, $p_{lk}\left(\sigma_{end}\right) = s_k\left(\sigma_{end}\right)/\sum_k s_k\left(\sigma_{end}\right)$), and $\bar{p}_k = \sum_l w_l p_{lk}$ is the mean relative abundance of species $k$ among communities $l = 1, \ldots, 100$. If all species go extinct in community $l$, it does not affect alpha, beta and gamma diversities as $w_l = 0$. If all species go extinct in all communities, beta diversity becomes ${}^{1}D_{\beta}\left(\sigma_{end}\right) = 1$. Fig. A.3 represents how alpha, beta, and gamma diversities change over the switching rate and mean toxin sensitivities.

### Appendix 2 Analysis excluding noise

#### Appendix 2.1 Deterministic scenarios (absence of DN and EFs)

In this section, we analyse the equilibrium state of a two-species, one-resource, and one-toxin model, in the absence of environmental switching and demographic noise. Although an equilibrium state cannot be analytically obtained, we shall see that there exists at most one stable and feasible equilibrium state. By removing demographic noise and environmental switching from Eq (4) in the main text, the dynamics are governed by the following ordinary differential equations:

$$
\dot{r_1} = \alpha\left(R_1 - r_1\right) - \sum_{k=1,2} \frac{\mu_{1k}}{Y_{1k}^{r}} \frac{r_1}{r_1 + K_{1k}^{r}} s_k, \tag{A.9a}
$$

$$
\dot{t_1} = \alpha\left(T_1 - t_1\right) - \sum_{k=1,2} \frac{\delta_{1k}}{Y_{1k}^{t}} \frac{t_1}{t_1 + K_{1k}^{t}} s_k, \tag{A.9b}
$$

$$
\dot{s_k} = \left(\mu_{1k}\frac{r_1}{r_1 + K_{1k}^{r}} - \delta_{1k}\frac{t_1}{t_1 + K_{1k}^{t}} - \alpha\right) s_k, \tag{A.9c}
$$

where the dot denotes the time derivative. In spite of the simplicity of the model, we learn from (A.9a) -(A.9c) that, in the absence of consumption by the species, the resources and toxins relax towards the in-flowing values $R_1$ and $T_1$ on a timescale of order $1/\alpha$. Furthermore, we infer from (A.9a) -(A.9c) that there are feedback loops between species abundances and resource and toxin concentrations (see Fig. 1A): as $s_k$ increases, both $r_1$ and $t_1$ decrease, see (A.9a) and (A.9b), which in turn can result either in a decrease (negative feedback loop) or in an increase of $s_k$ (positive feedback loop), depending on the sign of the parenthesis on the right-hand-side of (A.9c).

When only species $k$ persists in the system, a feasible[1] equilibrium state of Eqs (A.9a) -(A.9c), $(r_1, t_1, s_k) =$

[1] Here, feasibility means $r_{1k}^{*}, t_{1k}^{*}, s_k^{*} > 0$

$(r^*_{1k}, t^*_{1k}, s^*_k)$, should satisfy below:

$$\alpha\left(R_1 - r^*_{1k}\right) = \frac{\mu_{1k}}{Y^r_{1k}} \frac{r^*_{1k}}{r^*_{1k} + K^r_{1k}} s^*_k, \tag{A.10a}$$

$$\alpha\left(T_1 - t^*_{1k}\right) = \delta_{1k} Y^t_{1k} \frac{t^*_{1k}}{t^*_{1k} + K^t_{1k}} t^*_k, \tag{A.10b}$$

$$\mu_{1j} \frac{r^*_{1k}}{r^*_{1k} + K^r_{1k}} = \delta_{1k} \frac{t^*_{1k}}{t^*_{1k} + K^t_{1k}} + \alpha. \tag{A.10c}$$

By rearranging the first and second equations, we see that they represent quadratic functions of $r^*_{1k}$ and $t^*_{1k}$:

$$-r^{*2}_{1k} + \left\{R_1 - K^r_{1k} - \mu_{1k} s^*_k / (\alpha Y^r_{1k})\right\} r^*_{1k} + K^r_{1k} R_1 = 0, \tag{A.11}$$

$$-t^{*2}_{1k} + \left\{T_1 - K^t_{1k} - \delta_{1k} s^*_k / (\alpha Y^t_{1k})\right\} t^*_{1k} + K^t_{1k} T_1 = 0. \tag{A.12}$$

Notice that $K^r_{1k} R_1$ and $K^t_{1k} T_1$ are positive. This implies that the above equations always have a unique positive root. In other words, we can obtain unique solutions for $r^*_{1k}$ and $t^*_{1k}$ once we obtain $s^*_k$.

By substituting the positive roots of Eqs (A.11) and (A.12) into Eq (A.10c), we obtain the following equation whose positive root is $s^*_k$:

$$f(s) = \frac{1}{2\alpha} \left\{ \left(-\mu_{1k} + 2\alpha + \delta_{1k}\right) s + \sqrt{Q_1(s)} - \sqrt{Q_2(s)} + \alpha\left(-K^r_{1k} Y^r_{1k} + K^t_{1k} Y^t_{1k} - Y^r_{1k} R_1 + Y^t_{1k} T_1\right) \right\} \tag{A.13}$$

where $Q_1(s)$ and $Q_2(s)$ are quadratic functions of $s$:

$$Q_1(s) = \left\{\mu_{1k} s + \alpha Y^r_{1k}\left(K^r_{1k} - R_1\right)\right\}^2 + 4\alpha Y^r_{1k} K^r_{1k} R_1 > 0 \tag{A.14a}$$

$$Q_2(s) = \left\{\delta_{1k} s + \alpha Y^t_{1k}\left(K^t_{1k} - T_1\right)\right\}^2 + 4\alpha Y^t_{1k} K^t_{1k} T_1 > 0. \tag{A.14b}$$

Notice that $f(s)$ always has a root $s = 0$ because

$$\begin{aligned} f(0) &= \frac{1}{2\alpha} \left\{ \sqrt{Q_1(0)} - \sqrt{Q_2(0)} + \alpha\left(-K^r_{1k} Y^r_{1k} + K^t_{1k} Y^t_{1k} - Y^r_{1k} R_1 + Y^t_{1k} T_1\right) \right\} \\ &= \frac{1}{2\alpha} \left\{ \alpha Y^r_{1k}\left(K^r_{1k} + R_1\right) - \alpha Y^t_{1k}\left(K^t_{1k} + T_1\right) + \alpha\left(-K^r_{1k} Y^r_{1k} + K^t_{1k} Y^t_{1k} - Y^r_{1k} T_1 + Y^t_{1k} T_1\right) \right\} \\ &= 0. \end{aligned} \tag{A.15}$$

Although Newton's method numerically provides a root of $f(s)$, this root depends on the initial value used in Newton's method. In addition, as $f(s)$ has root $s = 0$, Newton's method may provide this root with various initial values, which does not always mean that $f(s)$ does not have positive roots (i.e., $s^*_k$). In other words, it is recommended to investigate how many positive roots $f(s)$ has before using Newton's method.

To investigate the number of $f(s)$'s positive roots, it is useful to obtain $df/ds$:

$$\frac{df}{ds} = \frac{1}{2\alpha} \left\{ \left(-\mu_{1k} + 2\alpha + \delta_{1k}\right) + \frac{dQ_1/ds}{2\sqrt{Q_1(s)}} - \frac{dQ_2/ds}{2\sqrt{Q_2(s)}} \right\}. \tag{A.16}$$

Although it is analytically difficult to obtain the solution(s) of $df/ds$, we can obtain the maximum number of positive roots of $f(s)$ by analyzing the number of $df/ds$'s sign changes. Notice that $dQ_1/ds$ and $dQ_2/ds$ are linear functions of $s$:

$$\frac{dQ_1}{ds} = 2\mu_{1k}\left\{\mu_{1k}s + \alpha Y^r_{1k}\left(K^r_{1k} - R_1\right)\right\}, \tag{A.17a}$$

$$\frac{dQ_2}{ds} = 2\delta_{1k}\left\{\delta_{1k}s + \alpha Y^t_{1k}\left(K^t_{1k} - T_1\right)\right\}. \tag{A.17b}$$

In addition, $Q_1(s)$ and $Q_2(s)$ are always positive. Then, the second and third terms of Eq (A.16) change their sign at most once by increasing $s$. The maximum number of $df/ds$'s sign change is, therefore, two. This implies that the maximum number of positive roots of $f(s)$ is also two. To obtain the exact number of positive roots of $f(s)$, it is necessary to numerically calculate the root(s) of $df/ds$. Substituting the root(s) into Eq (A.13) and calculating the sign of $f(s)$, it is possible to obtain the exact value of $s^*_k$.

Once we have obtained a feasible equilibrium state $(r^*_{1k}, s^*_k, t^*_{1k})$, it is necessary to analyze the stability of this equilibrium state. Although the stability analysis requires the evaluation of the Jacobian matrix at each equilibrium state, we can see that there exists at most one feasible and stable equilibrium state without such a stability analysis. Notice that:

$$\begin{aligned} \dot{s_j} &\lesseqgtr 0 \\ \Leftrightarrow \mu_{1k}\frac{r^*_{1k}}{r^*_{1k} + K^r_{1k}} &\lesseqgtr \delta_{1k}\frac{t^*_{1k}}{t^*_{1k} + K^t_{1k}} + \alpha \\ \Leftrightarrow f(s) &\gtreqless 0. \end{aligned} \tag{A.18}$$

These inequalities imply that a stable equilibrium state satisfies the following inequality:

$$\left.\frac{df}{ds}\right|_{s=s^*_k} > 0. \tag{A.19}$$

Although there can exist at most two feasible equilibria, only one of them satisfies the above inequality (Fig. A.1). The number of feasible and stable equilibria is, therefore, one at most.

For the sake of simplicity but without loss of generality, in the main text, we assumed that the maximum growth rate of species 1 is larger than species 2 but the remaining parameter values are identical, resulting in a *per-capita* growth rate of species 1 always larger than species 2. In this setting, in the absence of DN, species 2 always goes extinct after a finite time while species 1 persists if it has a feasible and stable equilibrium state. It is worth noting that this feature also characterizes the two-species, one-resource and one-toxin model in the presence of environmental switching without DN: in this case species 2 always goes extinct in a finite time, and at equilibrium one has either $s^*_1 = s^*_2 = 0$ (extinction of both species) or $s^*_1 > 0, s^*_2 = 0$ (survival of species 1, extinction of species 2), see Fig. A.5.

#### Appendix 2.2 With environmental fluctuations alone

It is also instructive to analyze the long-time dynamics of the two-species, one-resource, one-toxin model in the presence of EFs and without DN. This describes the dynamics of a sufficiently large community in which DN is always negligible and whose time evolution is described by Eqs (A.9a)-(A.9c), but now with $(R_1, T_1) = (R_1(\xi), T_1(\xi))$ switching randomly with rate $\nu$ between two states $\xi = \pm 1$, which is applicable to all three scenarios in Table 1. Here, for the sake of simplicity and concreteness, we consider the scenario 1 of Table 1 with $1 \ll R_1^- < \langle R_1 \rangle < R_1^+$, while $T_1 = \langle T_1 \rangle$ does not vary with the environment. In this case, Eqs. (A.9a)-(A.9c) are a system of stochastic differential equations (SDEs). In order to appreciate the effect of EFs on DN, it is useful to study the total population size $n \equiv r_1 + t_1 + s_1 + s_2$ which, according to (A.9a)-(A.9c), obeys

$$\dot{n} = \alpha \left( R_1(\xi) + \langle T_1 \rangle - n \right) - 2 \sum_{k=1,2} \delta_{1,k} \, \frac{t_1 s_k}{t_1 + K_{1,k}^t}, \tag{A.20}$$

where, as in the main text, $Y_{ik}^r = Y_{jk}^t = 1$. Importantly, $\sqrt{n}$ gives the intensity of DN in a total population of size $n$. Eq (A.20) is, however, difficult to analyze, due to the nonlinear coupling of $t_1$ with $s_k$ whose dynamics, according to Eqs (A.9b)-(A.9c), also depend on $\xi$. In order to obtain some insight into the stationary probability density $p(n)$ of the population, we have thus we analyzed $\hat{n} \equiv r_1 - t_1 + s_1 + s_2$ which is related to the total population size, since $\hat{n} = n - 2t_1$. In this simple example $\hat{n}$, according to Eqs. (A.9a)-(A.9c), obeys the following linear SDE

$$\dot{\hat{n}} = \alpha \left( R_1(\xi) - \langle T_1 \rangle - \hat{n} \right). \tag{A.21}$$

As seen above, the system reaches its equilibrium and $(s_1, s_2, t_1) \to (s_1^*, s_2^* = 0, t_1^*)$ after a finite time while the environment varies endlessly .The SDE (A.21) defines a simple piecewise deterministic Markov process (PDMP) (Davis, 1984) whose (marginal) probability density $p_{\nu/\alpha}(\hat{n})$ can be obtained analytically, see Bena (2006) and Horsthemke and Lefever (2006):

$$q_{\nu/\alpha}(\hat{n}) = \mathcal{Z} \; \left[ \left\{ R_1^+ - \langle T_1 \rangle - \hat{n} \right\} \left\{ \hat{n} - R_1^- + \langle T_1 \rangle \right\} \right]^{\frac{\nu}{\alpha} - 1}, \tag{A.22}$$

where $\mathcal{Z}$ is the normalization constant. Hence, in the absence of DN, with environmental switching as the sole source of randomness, the dynamics leads to a population consisting of only individuals of species 1 (species 2 is wiped out) and toxin and resources, with respective abundances $s_1, r_1, t_1$, in a fluctuating population whose scaled size $\hat{n} = s_1 + r_1 - t_1$ is distributed according to $q_{\nu/\alpha}(\hat{n})$ of finite support $\left[ R_1^- - \langle T_1 \rangle, R_1^+ - \langle T_1 \rangle \right]$ (i.e. $R_1^- - \langle T_1 \rangle \leq n \leq R_1^+ - \langle T_1 \rangle$). While this PDMP probability density ignores DN, it is known to generally provide a useful approximate description of how the quasi-stationary (at finite time $\gtrsim \sigma_{end}$) distribution varies with the environment in a large yet finite system, see, *e.g.*, (Wienand et al., 2017, 2018; West and Mobilia, 2020;

Taitelbaum et al., 2020). Here, Eq. (A.22) readily sheds light on the effect of $\nu$ and $\alpha$ on $\hat{n} = n - 2t_1$. It is clear from Eqs. (A.9a)-(A.9c), that $t_1$ varies with $\xi$ and thus the marginal probability density of population size, $p(n)$, cannot be immediately obtained from (A.22). Yet, we can obtain useful information about how the population varies with $\nu$ and $\alpha$ by focusing on the fast and slow switching regimes (see Fig. A.6):

- when $\nu/\alpha \ll 1$ (slow varying environment), $q_{\nu/\alpha}(\hat{n})$ is bimodal and $\hat{n}$ is as likely to fluctuate about $R_1^+ - \langle T_1 \rangle$ or $R_1^- - \langle T_1 \rangle$. Hence, when in this slowly switching regime, the population size fluctuates with the same probability either about $R_1^- - \langle T_1 \rangle + 2t_1^*(\xi = -1)$ or $R_1^+ - \langle T_1 \rangle + 2t_1^*(\xi = +1)$ (see, the first column of Fig. A.6).

- when $\nu/\alpha \gg 1$ (fast changing environment), $q_{\nu/\alpha}(\hat{n})$ is unimodal and $\hat{n}$ fluctuates about its average, i.e. $\hat{n} \approx \langle R_1 \rangle - \langle T_1 \rangle$. Hence, in this fast switching regime, the population size fluctuates about $n \approx \hat{n} + 2\langle t_1 \rangle = \langle R_1 \rangle - \langle T_1 \rangle + 2\langle t_1 \rangle$, where $\langle t_1 \rangle = (t_1^*(\xi = 1) + t_1^*(\xi = -1))/2$ and $t_1^*(\xi = \pm 1)$ are obtained from the equilibria of Eqs. (A.9a)-(A.9c) with $R_1 = R_1(\xi = \pm 1)$ (see, the third column of Fig. A.6).

This picture is confirmed by the simulation results reported in Fig. A.6, where we see that the marginal probability densities $p(n)$ and $q_{\nu/\alpha}(\hat{n})$ have qualitatively the same features: both are bimodal and have two well-separated sharp peaks when $\nu/\alpha \ll 1$, and a single pronounced peak when $\nu/\alpha \gg 1$. At intermediate values of $\nu/\alpha$, the probability densities of $n$ and $\hat{n}$ are much broader with a generally flat profile (exhibiting one or two "bumps"), see the second column of Fig. A.6. This analysis, which can be readily extended to $N > 2$ species and to other scenarios of environmental fluctuations, clearly shows that the population size can greatly vary as the environment changes. In particular, Eq (A.22) shows that when $\nu$ is low or $\alpha$ is high, half of the simulation runs lead to communities of "small sizes" where the effect of DN is expected to be significantly larger than in communities obtained in the faster switching regime ($\nu/\alpha \gg 1$) when $R_1^+ \gg R_1^-$.

It is also worth noting from Eqs (A.9a)-(A.9c) and Eq (A.21) that if, say, only the maximum growth rates were subject to environmental switching, i.e. $\mu_{ik} = \mu_{ik}(\xi)$ with all other parameters kept constant, we always obtain a large constant population size (if $R_1 \gg T_1$) and thus no DN-EFs coupling because the distributions of $n$ and $\hat{n}$ are independent on $\mu_{ik}$. On the other hand, if only the maximum death rates were subject to environmental switching, i.e. $\delta_{jk} = \delta_{jk}(\xi)$, the DN-EFs coupling would result from a complicated set of coupled stochastic differential equations obtained from (A.9a)-(A.9c) with $\delta_{jk} \to \delta_{jk}(\xi)$. These analyses clarify the significant difference of our model from others. Some previous studies (e.g., Leigh (1981); Kalyuzhny et al. (2015) and multi-species model of Engen and Lande (1996)) do not include DN-EFs coupling because they assume that EFs affect species' growth rates, but total species abundances or maximum population sizes do not change. Other studies (e.g., Kamenev et al. (2008); Chisholm et al. (2014); Fung et al. (2015) and the single species model of Engen and Lande (1996)) include a form of DN-EF coupling because EFs in their model change both species' growth rates and population sizes. However, these models consist of only one species, and hence do not consider interspecific interactions. Our model includes both DN-EFs and indirect species interactions (resource competition and facilitation via detoxification, see Fig. 1A), and thus we can analyze how DN-EFs coupling affect species diversity as in Fig. 6. In summary, here we have attempted to make the simplest choice to couple DN and EF in a transparent and biologically-relevant way.

### Appendix 3 Alternative environmental switching scenarios

In the main text, environmental switching affects only the resource supply (scenario 1) while the amount of toxin supply is fixed. Here, the results of other environmental switching scenarios are shown: In scenario 2, environmental switching affects only the toxin supply, while both resource and toxin supplies change and correlate negatively in scenario 3 (see Table 1).

In both scenarios 2 and 3, the difference in species 1's extinction probability $\Delta P\left(s_1(\sigma_{end}) = 0\right)$ is very similar to the negative value of the probability of exclusion of the fittest when the sign of $\Delta P\left(s_1(\sigma_{end}) = 0\right)$ is negative (Fig. A.7). This once again confirms that we can use one measure for the other.

In addition, in both scenarios, the probability of exclusion of the fittest are bi-modal across toxin sensitivities at very slow environmental switching $\nu = 10^{-5}$, but uni-modal at very fast environmental switching ($\nu = 10^3$) (Fig. A.7C, D). As explained in the main text, when $\nu \to 0$, there are no switches and the environmental state is randomly allocated to harsh or mild conditions at $t = 0$ with the same probability (the mean of $\xi$ is zero), yielding a bi-modal distribution at low switching rate. In the limit $\nu \to \infty$, there are so many switches that environmental noise averages out, i.e. $\xi$ is replaced by its mean (that is zero). This results in a uni-modal distribution when $\nu \gg 1$.

In scenario 2, however, we do not observe any non-monotonic changes over the switching rate (Fig. A.7C). This is because the critical toxin sensitivity under abundant toxin supply ($\delta = 0.3$) is close to that under mean toxin supply ($\delta = 0.4$). In contrast, we do observe non-monotonic effects of the switching rate in scenario 3 (Fig. A.7D): when $\delta = 0.2$, an intermediate switching rate ($\nu = 10^{-1}$) shows the minimum difference in extinction probability, and the maximum probability of competitive exclusion. Although in scenario 3 the same intermediate switching rate minimizes the probability of competitive exclusion at toxin sensitivity 0.6, the same non-monotonic effect is not observed in the difference in extinction probabilities (Fig. A.7B). Note that the critical toxin sensitivities under the mild environments in scenarios 2 (scarce toxin supply) and 3 (abundant resource supply and scarce toxin supply) are slightly larger than 1.0 (Fig. A.8) and therefore not visible in Fig. A.7D. Table A.2 shows the critical toxin sensitivities in each scenario.

### Appendix 4 Effects of resource supply

In this section, we again focus on environmental switching scenario 1 (changing only resource supply). We will see that the amount of resource supplies $R_1(\xi)$ changes the critical toxin sensitivities under scarce, mean, and abundant, resource supplies, affecting the likelihood of the non-monotonic effect of the environmental switching rate.

By increasing the abundant or decreasing the scarce resource supply (Figs. A.9A and D, respectively), the distance between the critical toxin sensitivities under scarce and mean (or mean and abundant) resource supplies becomes larger (Table A.2). Conversely, decreasing abundant or increasing scarce resource supply (Figs. A.9B and C, respectively), decreases the distance between the critical toxin sensitivities under scarce and mean (or mean and abundant) resource supplies (Table A.2). Once again, changes in competitive exclusion probability (A.9) match changes in species 2's effect on species 1 (Fig. A.10).

Analyzing the probability of exclusion of the fittest instead of the difference in extinction probability is valid only when the difference in extinction probability is negative: if it is positive, the second line in (A.2) cannot be ignored. The sign of the difference can, however, become positive at $\delta = 1.0$. Indeed, when $R_1^+ = 400$ and $\delta = 1.0$, species 2 has a positive effect on species 1 whose strength varies non-monotonically with the rate of environmental switching (A.11A). In this case, we analyze both (i) the probability of exclusion of the fittest (the first line in Eq (A.2), Fig. A.11B) and (ii) the difference in species 1's extinction probability in mono-culture and both species extinction in co-culture (the second line in Eq (A.2), Fig. A.11C). The effects of the environmental switching rate on (i) and (ii) are similar, leading to similar non-monotonic effects of species 2 on species 1. In sum, non-monotonic effects of environmental switching rates on species interactions can be observed whether these interactions are positive or negative. Although the main text explains why non-monotonic effects of environmental switching rates on species interactions happen when interactions are negative, it remains unclear why such non-monotonic changes happen when species interactions are positive.

### Appendix 5 Other forms of environmental fluctuations

In this section, we analyze environmental fluctuations other than symmetric switching between two states. Our goal is here to show that our main findings qualitatively still hold and can therefore traced back to the generic interdependence of EFs and DN rather than detail of their coupling.

As in the main text, we assume that the environmental fluctuations change only the resource supply while the toxin supply is constant. First, we investigate asymmetric switching between two resource supply conditions. Second, we increase the number of environmental states and introduce a cyclic change of the resource supply.

Under asymmetrically switching or cyclically fluctuating environments, we find similar patterns of how species interactions change over species' toxin sensitivities and a rate of environmental fluctuations.

#### Appendix 5.1 Asymmetric switching

In the main text and appendices other than this section, for the sake of simplicity, we assume symmetric switching rates between two states by Eq. (3) (see Taitelbaum et al. (2020)). Here, we relax this assumption and introduce asymmetric switching rates because perfectly symmetric switching environments are very unlikely in nature; in gut microbiota, for example, duration that their host is starving would be longer than that the host is eating food. We implemented asymmetric switching as follows:

$$\xi = 1 \xrightarrow{\nu_1} \xi = -1 \tag{A.23a}$$

$$\xi = -1 \xrightarrow{\nu_2} \xi = 1. \tag{A.23b}$$

Without loss of generality, we define the two switching rates as follows:

$$\nu_1 = \beta_1 \nu \tag{A.24a}$$

$$\nu_2 = \beta_2 \nu \tag{A.24b}$$

where $\nu$ is the basal switching rates. In extreme cases ($\beta_1 \gg \beta_2$), for asymmetric switching scenarios can correspond to systems with rare disturbances in nature. In this extremely asymmetric case, the sojourn time in the harsh environment exceeds greatly that in the mild environment, which results in a strong effect of DN. In contrast, DN effects are less important when $\beta_2 \gg \beta_1$ and the population experiences more frequently the mild than the harsh environment(Taitelbaum et al., 2020). We recover a symmetric environmental switching when $\beta_1 = \beta_2$.

Fig. A.12 summarizes the two species interactions under resource supply fluctuations with asymmetric environmental switching rates when the initial environmental condition is $\xi(0) = 1$ with probability of 0.5 (otherwise $\xi(0) = -1$). In Figs. A.12 A and C, $\nu_1 > \nu_2$ and therefore the sojourn time of $\xi = -1$ (an harsh environment) is longer than that of $\xi = 1$ (a mild environment). On the other hand, Figs. A.12 B and D shows the cases when the sojourn time the mild environment is longer than that of the harsh environment because $\nu_1 < \nu_2$. In both cases, species 1's difference in extinction probabilities and the probability of exclusion of the fittest show monotonically increasing, monotonically decreasing, or non-monotonic changing with a minimum or maximum value at an intermediate switching rate, although they quantitatively differ from Figs. 3 A and C.

#### Appendix 5.2 Cyclic changes

Here, we analyze cases when the number of environmental states in terms of resource supply is greater than two, but remains discrete and finite. This simply reflects that natural environments do not always fluctuate between two states. In this subsection, an environmental state is given by $\xi = 1, 2, \ldots, n$ and environments cyclically fluctuate with rate $\nu$ as follows:

$$\xi \xrightarrow{\nu} \begin{cases} \xi + 1 & \text{if } \xi = 1, \ldots, n-1 \\ 1 & \text{otherwise.} \end{cases} \tag{A.25}$$

This is a natural extension of Eq (3) by increasing the number of environmental conditions: $n = 2$ recovers a symmetrically switching environment between two conditions ($\xi = 1 \to 2 \to 1 \to \ldots$ although we use the notation $\xi = \pm 1 \to \mp 1 \to \pm 1 \to \ldots$ in the main text).

Fig. A.13 shows how species interactions between two species change when $n = 4$ and the resource supply fluctuates such that $R_1(\xi = 1) > R_1(\xi = 2) > R_1(\xi = 3) > R_1(\xi = 4)$. In this analysis, an initial environmental condition $\xi(0)$ is one of four conditions (probability of 0.25 for each). As in Figs. 3 A and C, the rate of cyclical environmental change affects species 1's extinction probability and the probability that species 2 excludes species 1. We frequently observe that the probability of exclusion of the fittest non-monotonically changes (toxin sensitivity: 0.1 – 0.6).

In this work, the fluctuating environment has been modeled as randomly switching between a finite number of environmental states $\xi$. This choice is particularly convenient as it allows us to deal with bounded noise, and hence $R_i$ and $T_j$ to always remain positive, and are straightforward to simulate using the standard Gillespie algorithm. The case of environmental noise varying continuously in time is also of great interest, as it allows $R_i$ and $T_j$ to take any values in a domain. For instance, the environmental noise can be an Ornstein-Uhlenbeck process ($\xi_{OU}$), e.g. by letting $R_i = R_i(\xi_{OU}) = \bar{R}_i(1 + k\xi_{OU})$, where $\bar{R}_i$ is constant, $\xi$ varies in time, and $k > 0$, see, e.g., Assaf et al. (2013). This poses a number of challenges since, $\xi_{OU}$ being unbounded, $R_i$ can take negative (unphysical) values. Furthermore, there are no general methods to simulate exactly birth-death processes subject to continuous external noise, see, e.g., Berríos-Caro and Galla (2020). Here, while the assumption of discrete environmental noise is a simplification of many real situations, we think that our main findings are generic and shall hold also under continuous environmental noise. In fact, since our results stem chiefly from the coupling of EFs and DN, a feature shared by discrete and continuous noise, they are expected to qualitatively hold also in the case of continuous external noise.

#### Appendix 6 Diversity in communities of increasing species number

In the main text, we show the distributions of beta diversity and species richness in communities when the initial number of species $N$ is two or ten. This section shows the results of intermediates values of $N = 4, 6, 8$. The number of initial species does not change how beta diversity changes over the environmental switching rate, except when the mean toxin sensitivity $\bar{\delta} = 0.4$ (Fig. A.18). These results indicate that beta diversity and the probability of competitive exclusion change similarly over the switching rate when the initial number of species is larger than two.

The initial number of species $N$ in a community affects the maximum values of species richness (Fig. A.19). However, how the switching rate and the mean toxin sensitivity affect the distribution of species richness was consistent for different values of $N$. In particular, increasing the mean toxin sensitivity decreases species richness. The effects of the environmental switching rate also depend on the mean toxin sensitivity: at mean toxin sensitivity 1.0, species in all cases are more likely to go extinct as the switching rate increases, while the likelihood of all species going extinct consistently shows a humped shape at toxin sensitivity 0.4 or 0.6.

#### Appendix 7 Quantified the similarity between the exclusion of fittest and beta diversity

In this section, we first quantify the similarity between the exclusion of the fittest and beta diversity in two-species communities. Then, we investigate how many species pairs we should analyze to predict the patterns of beta diversity in ten-species communities.

In the main text, we show that the probability of exclusion of the fittest and beta diversity exhibit similar patterns (see columns A and B in Fig. 6). We quantified the similarities with Pearson's correlation coefficients. Fig. A.15 shows the distributions of the correlation coefficients of 100 two-species communities at each mean toxin sensitivity (see also Appendix 1.3). Except for the case that mean toxin sensitivity is 0.4 (where the probability of exclusion of the fittest and beta diversity do not match), the correlation coefficients are large positive ($> 0.6$). Therefore, a large correlation coefficients indicates the similarity between the probability of the exclusion of the fittest and beta diversity.

We continued the analysis in ten-species communities. In these cases, we have 45 pairs of species in each community and we calculated the probabilities of the exclusion of the fittest by running 100 replicates in each species pair of thirty ten-species communities (six communities at five mean toxin sensitivities). Fig. A.16 shows that the exclusion of the fittest in some pairs match the patterns of beta diversity of whole communities but other pairs do not. Then, we investigated the number of species pairs $m$ that is necessary to predict a pattern of beta diversity over the environmental switching rate (i.e., a large correlation between probability of exclusion of the fittest and beta diversity). When $m \geq 2$, we calculated Pearson's correlation coefficient between beta diversity and mean probability of exclusion of the fittest within $m$ pairs. However, we have many possible choices of $m$ pairs when $1 < m < 45$. In these cases, we randomly chose 300 sets of $m$ pairs in each of community and thus we obtained 1800 correlation coefficients at each mean toxin sensitivities. When $m = 1$ or 45, we analyzed all possible choices of $m$ species pairs and we obtained 45 or 1 correlation coefficient(s) in each community, respectively. Fig. A.17 indicates that the correlation coefficients can be large even when $m = 1$ or 2, but larger $m$ increases correlation coefficients. We suggest that $m = 5$ is the best because the $m = 5$ and $m = 9$ show little difference in the distributions of the correlation coefficients and we are very likely to obtain a large correlation coefficient with $m = 5$. In conclusion, we do not have to analyze the exclusion of the fittest for all species pairs to predict how the environmental switching rate affect beta diversity.

#### References

Assaf, M., Mobilia, M., and Roberts, E. Cooperation dilemma in finite populations under fluctuating environments. Physical Review Letters, 111(23):1–7, 2013. ISSN 00319007. doi: 10.1103/PhysRevLett.111.238101.

Bena, I. Dichotomous Markov noise: Exact results for out-of-equilibrium systems. A review. International Journal of Modern Physics B, 20(20):2825–2888, 6 2006. ISSN 02179792. doi: 10.1142/S0217979206034881. URL http://arxiv.org/abs/cond-mat/0606116.

Berríos-Caro, E. and Galla, T. Beyond the adiabatic limit in systems with fast environments: a $\tau$-leaping algorithm. arXiv, pages 1–22, 2020. URL http://arxiv.org/abs/2011.10748.

Chisholm, R. A., Condit, R., Rahman, K. A., Baker, P. J., Bunyavejchewin, S., Chen, Y.-Y., Chuyong, G., Dattaraja, H. S., Davies, S., Ewango, C. E. N., Gunatilleke, C. V. S., Nimal Gunatilleke, I. A. U., Hubbell, S., Kenfack, D., Kiratiprayoon, S., Lin, Y., Makana, J.-R., Pongpattananurak, N., Pulla, S., Punchi-Manage, R., Sukumar, R., Su, S.-H., Sun, I.-F., Suresh, H. S., Tan, S., Thomas, D., and Yap, S. Temporal variability of forest communities: empirical estimates of population change in 4000 tree species. Ecology Letters, 17(7): 855–865, 7 2014. ISSN 1461023X. doi: 10.1111/ele.12296. URL http://doi.wiley.com/10.1111/ele.12296.

Davis, M. H. A. Piecewise-Deterministic Markov Processes: A General Class of Non-Diffusion Stochastic Models. Journal of the Royal Statistical Society: Series B (Methodological), 46(3):353–376, 1984. doi: 10.1111/j.2517-6161.1984.tb01308.x.

Engen, S. and Lande, R. Population Dynamic Models Generating Species Abundance Distributions of the Gamma Type. Journal of Theoretical Biology, 178(3):325–331, 2 1996. ISSN 00225193. doi: 10.1006/jtbi.1996.0028. URL https://linkinghub.elsevier.com/retrieve/pii/S0022519396900284.

Fung, T., Villain, L., and Chisholm, R. A. Analytical formulae for computing dominance from species-abundance distributions. Journal of Theoretical Biology, 386:147–158, 2015. ISSN 10958541. doi: 10.1016/j.jtbi.2015.09.011. URL http://dx.doi.org/10.1016/j.jtbi.2015.09.011.

Horsthemke, W. and Lefever, R. Noise-Induced Transitions, volume 15 of Springer Series in Synergetics. Springer Berlin Heidelberg, 2nd edition, 4 2006. ISBN 978-3-540-11359-1. doi: 10.1007/3-540-36852-3. URL http://link.springer.com/10.1007/3-540-36852-3.

Kalyuzhny, M., Kadmon, R., and Shnerb, N. M. A neutral theory with environmental stochasticity explains static and dynamic properties of ecological communities. Ecology Letters, 18(6):572–580, 6 2015. ISSN 1461023X. doi: 10.1111/ele.12439. URL http://doi.wiley.com/10.1111/ele.12439.

Kamenev, A., Meerson, B., and Shklovskii, B. How Colored Environmental Noise Affects Population Extinction. Physical Review Letters, 101(26):268103, 12 2008. ISSN 0031-9007. doi: 10.1103/PhysRevLett.101.268103. URL https://link.aps.org/doi/10.1103/PhysRevLett.101.268103.

Leigh, E. G. The average lifetime of a population in a varying environment. Journal of Theoretical Biology, 90(2):213–239, 5 1981. ISSN 00225193. doi: 10.1016/0022-5193(81)90044-8. URL https://linkinghub.elsevier.com/retrieve/pii/0022519381900448.

Taitelbaum, A., West, R., Assaf, M., and Mobilia, M. Population Dynamics in a Changing Environment: Random versus Periodic Switching. Physical Review Letters, 125(4):048105, 7 2020. ISSN 0031-9007. doi: 10.1103/PhysRevLett.125.048105. URL https://link.aps.org/doi/10.1103/PhysRevLett.125.048105.

West, R. and Mobilia, M. Fixation properties of rock-paper-scissors games in fluctuating populations. Journal of Theoretical Biology, 491:110135, 4 2020. ISSN 00225193. doi: 10.1016/j.jtbi.2019.110135. URL http://dx.doi.org/10.1016/j.jtbi.2019.110135.

Wienand, K., Frey, E., and Mobilia, M. Evolution of a Fluctuating Population in a Randomly Switching Environment. Physical Review Letters, 119(15):158301, 10 2017. ISSN 0031-9007. doi: 10.1103/PhysRevLett.119.158301. URL https://link.aps.org/doi/10.1103/PhysRevLett.119.158301.

Wienand, K., Frey, E., and Mobilia, M. Eco-evolutionary dynamics of a population with randomly switching carrying capacity. Journal of the Royal Society Interface, 15(145):20180343, 8 2018. ISSN 17425662. doi: 10.1098/rsif.2018.0343. URL https://royalsocietypublishing.org/doi/10.1098/rsif.2018.0343.

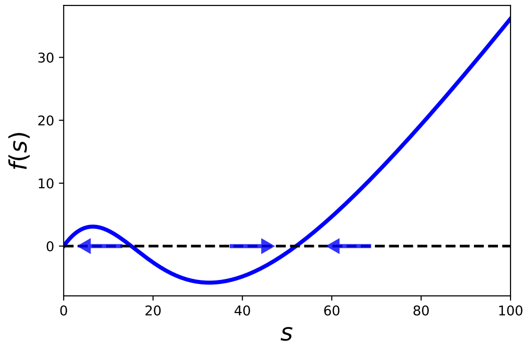


Figure A.1: Feasible and stable equilibrium state

An example of $f(s)$ in Eq (A.13) and equilibrium states. In this example, $f(s)$ has three roots: $s = 0$, and two feasible equilibria. The blue arrows indicate that $s$ decreases or increases when $f(s)$ is positive or negative, respectively. As $df/ds$ is negative at the left equilibrium state, this equilibrium state is unstable. On the other hand, the right feasible equilibrium has a positive $df/ds$ and thus this equilibrium can be stable. Note that the equilibrium state corresponding to $s = 0$ can be also stable. Parameter values are $\delta_{1k} = 1.2$, $R_1 = 200$, $T_1 = 125$ and as in Table A.1 otherwise.

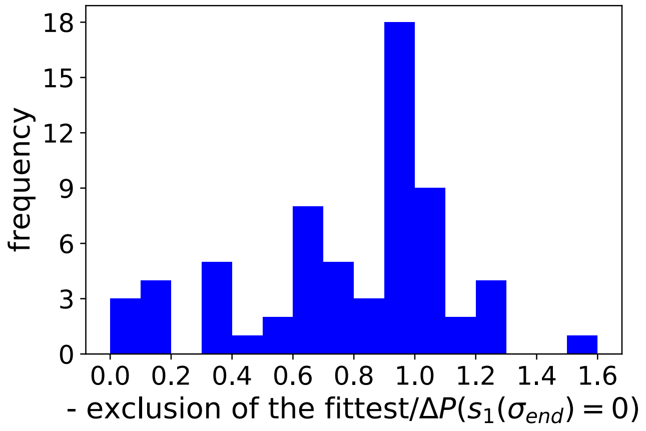


Figure A.2: Probability of exclusion of the fittest relative to the difference in extinction probability of species 1

A histogram of the ratio of the probability of exclusion of the fittest and the difference in species 1's extinction probabilities alone versus in the presence of species 2, showing the results of 81 sets of the environmental switching rate $\nu$ and the toxin sensitivity $\delta$ (9 values of $\nu = 10^{-5}, \ldots, 10^{3}$ and 9 values of $\delta = 0.1, \ldots, 0.9$). For each set of the parameter values, $10^5$ simulations were run to calculate the competitive exclusion probability and the difference in species 1's extinction probabilities in the presence/absence of species 2. In many of these 81 parameter sets, this ratio is close to 1, indicating that both measures yield similar results. As in the manuscript we focus on conditions leading to competition between the two species, we ignore toxin sensitivity $\delta = 1.0$ where species 2's effect on species 1 can be positive.

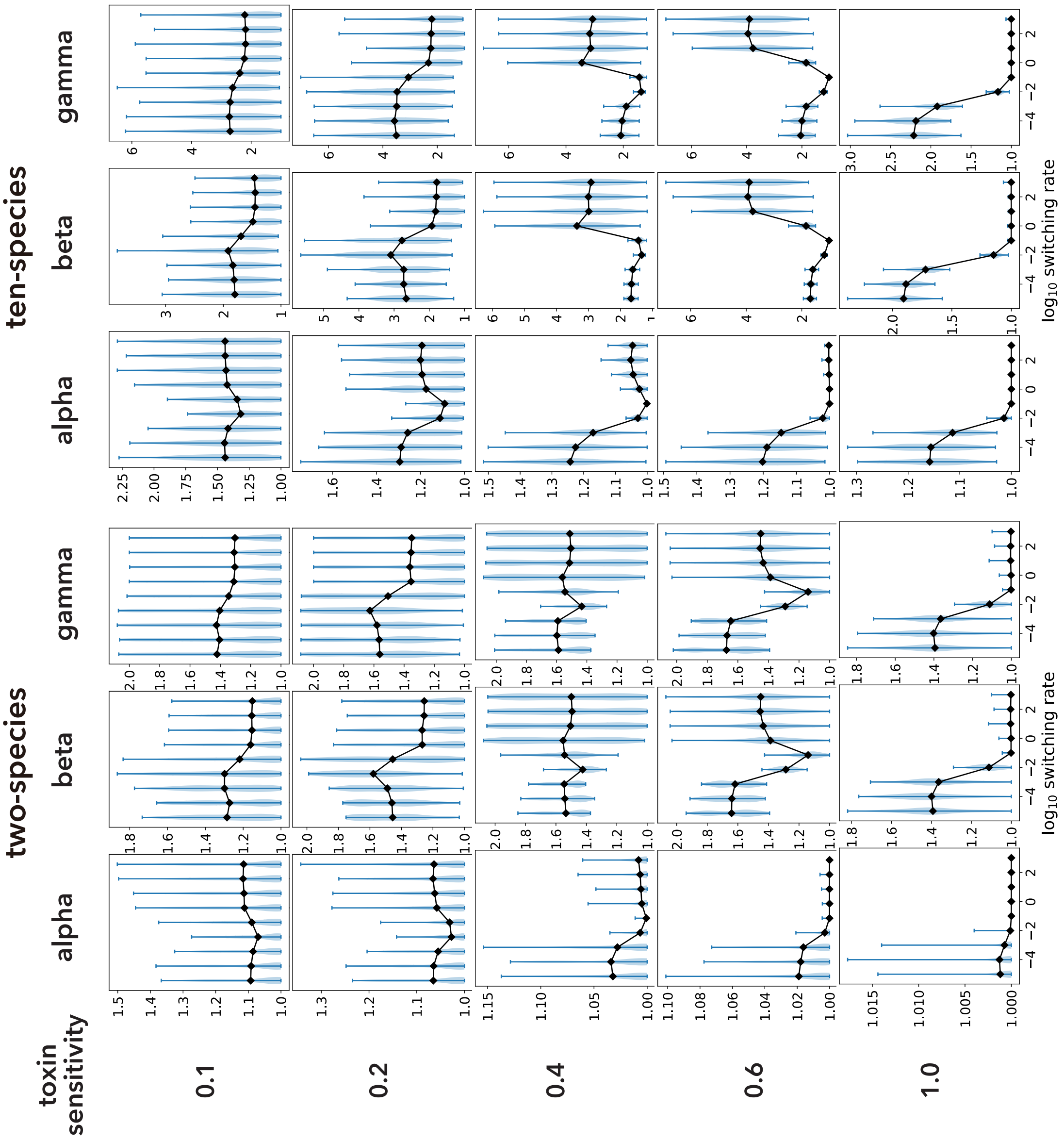


Figure A.3: Changes of alpha, beta, and gamma diversities

Alpha, beta, and gamma diversities over the switching rate and mean toxin sensitivity in two- and ten-species communities at the end of simulations. As alpha diversity is always closed to one, beta diversity and gamma diversity show similar trend. The black lines and blue areas represent the mean values and the probability distributions of the diversities calculated from 10’000 simulations. See Appendix 1.3 for more detail.

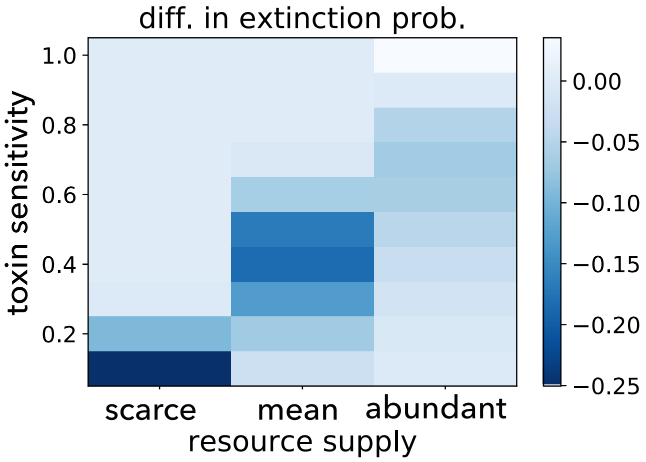


Figure A.4: Species 2's effect on species 1 in the absence of environmental switching

Species 2's effect on species 1 when the resource supply is fixed to be scarce ($R_1^-$), mean ($\langle R_1 \rangle$), or abundant ($R_1^+$). The toxin sensitivities that minimize species 2's effect on species 1 correspond to the peak sensitivities in Fig. 5.

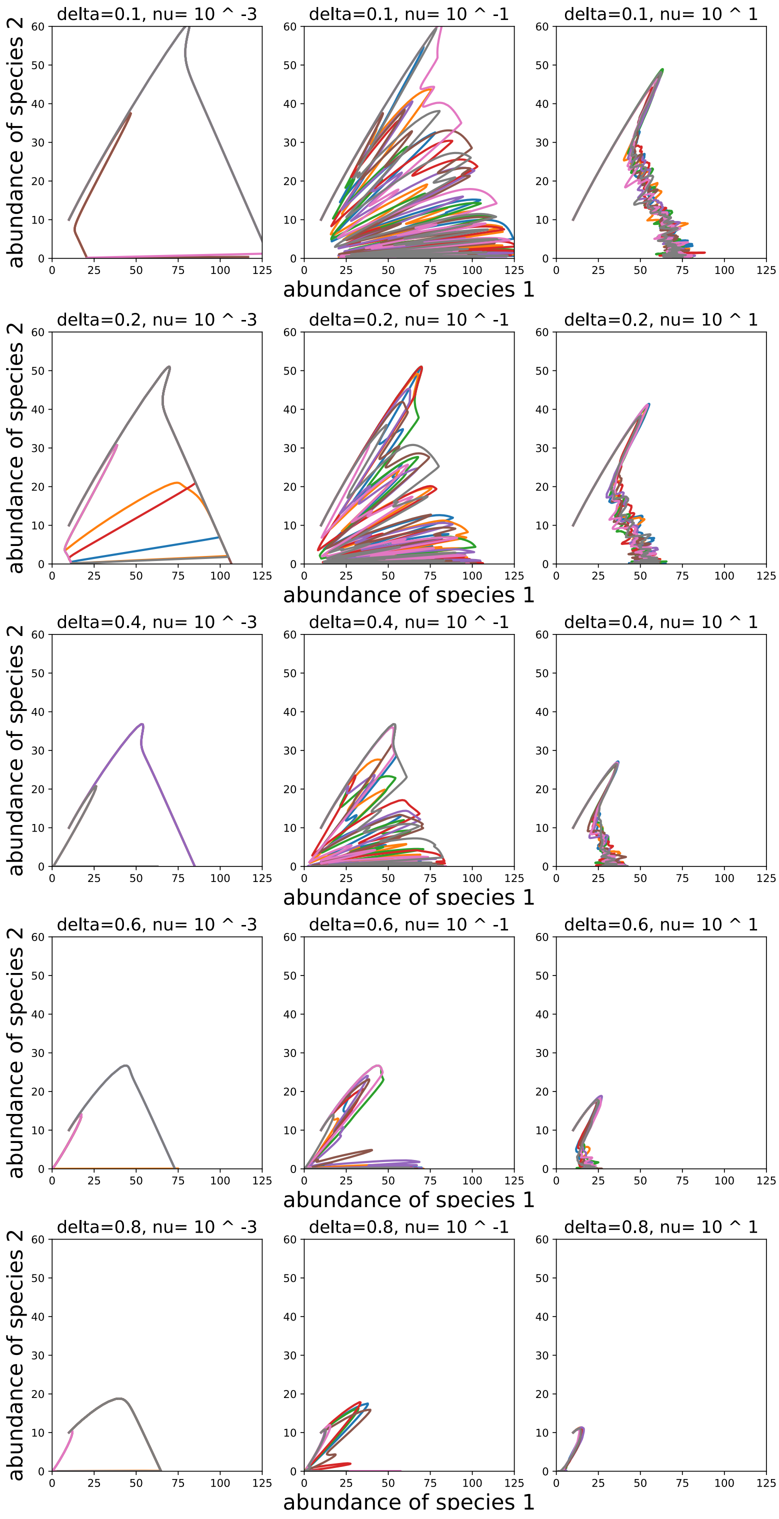


Figure A.5: Examples of the dynamics with only the environmental fluctuations

State transition of two species abundances $(s_1, s_2)$ in the absence of DN but the presence of the EFs are shown. Here, EFs switch resource supply (scenario 1 in Table 1) and the initial population abundances are $s_1(0) = s_2(0) = 10$. In this setting, species 2 always goes extinct ($\lim_{\sigma\to\infty} s_2(\sigma) = 0$) regardless of the values of $\delta$ and $\nu$. On the other hand, species 1 survives ($\lim_{\sigma\to\infty} s_1(\sigma) > 0$) if the environment is not too harsh. In each panel, different colors represent different samples of the dynamics with EFs alone. The values of $\nu$ and $\delta$ are shown on the top of each panel and the rest parameter values are shown in Table A.1.

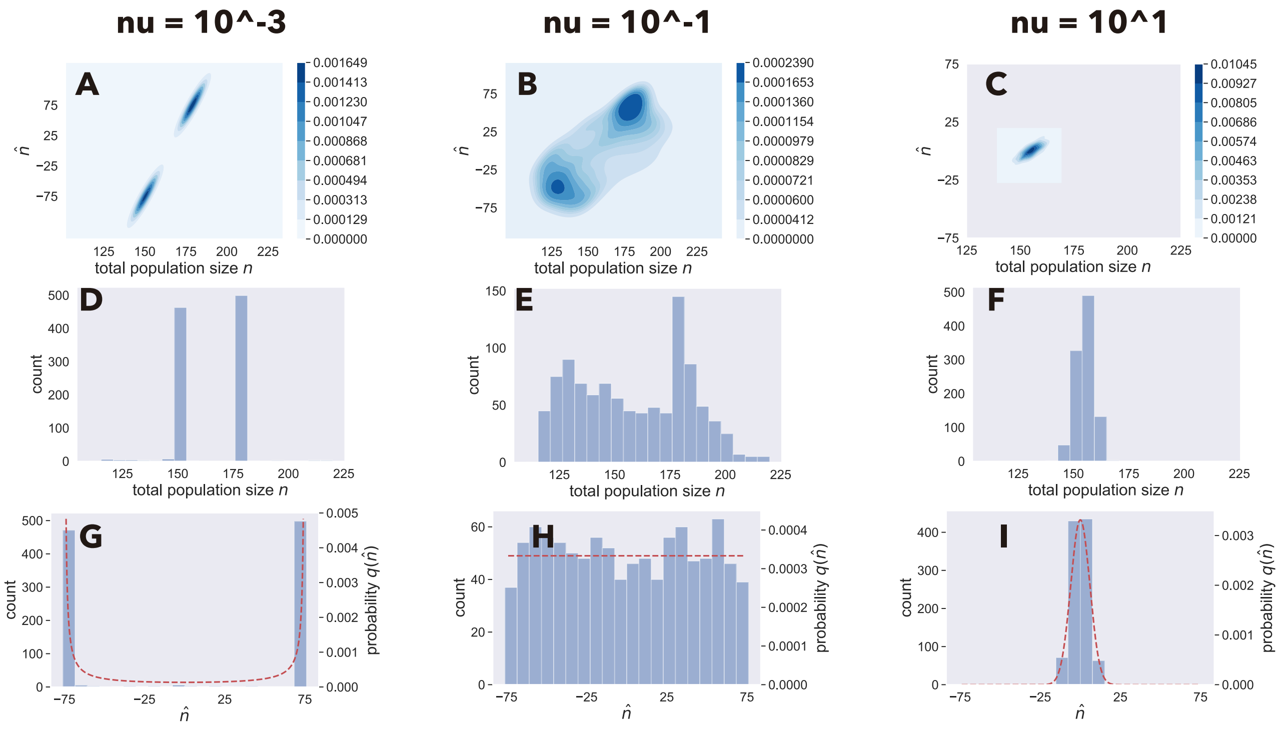


Figure A.6: Distributions of total population sizes

Probability distributions of total population size $n = r_1 + t_1 + s_1 + s_2$ and the auxiliary quantity $\hat{n} = r_1 - t_1 + s_1 + s_2$ obtained from 1,000 runs of Eqs (A.9a)-(A.9c) with environmental switching of scenario 1 in Table 1 at the slow $\nu = 10^{-3}$ (first column), intermediate $\nu = 10^{-1}$ (second column), or fast $\nu = 10^{1}$ (third column) switching rates. We collected the simulation data at time $\sigma_{end} = 200$. At the beginning of each simulation, $\xi = 1$ with 50 percents; otherwise $\xi = -1$. (A-C): the contour plots show the joint probability distributions of $n$ and $\hat{n}$: large $n$ corresponds to large $\hat{n}$. (D-F): the histograms show the distributions of the total population size $n$. (G-I): the histograms of $\hat{n}$ (left y-axis) and its theoretical probability distributions $q_{\nu/\alpha}(\hat{n})$ (right y-axis), which is given by Eq(A.22), are shown in blue bars and red dashed lines, respectively. We used $\delta_{1,1} = \delta_{1,2} = 0.2$ and all other parameter values are shown in Table A.1, and thus $\nu/\alpha = 0.01$ corresponds to the slow switching rate (first column), $\nu/\alpha = 1$ corresponds to the intermediate switching rate (second column), and $\nu/\alpha = 100$ corresponds to the fast switching rate (third column), respectively, see Appendix 2.2. Exceptionally, we used $\nu/\alpha = 80$ instead of $\nu/\alpha = 100$ to show $q_{\nu/\alpha}(\hat{n})$ in the third column because $\nu/\alpha = 100$ causes overflow during the calculation of $q_{\nu/\alpha}(\hat{n})$, but this modification does not change the qualitative feature of $q_{\nu/\alpha}(\hat{n})$.

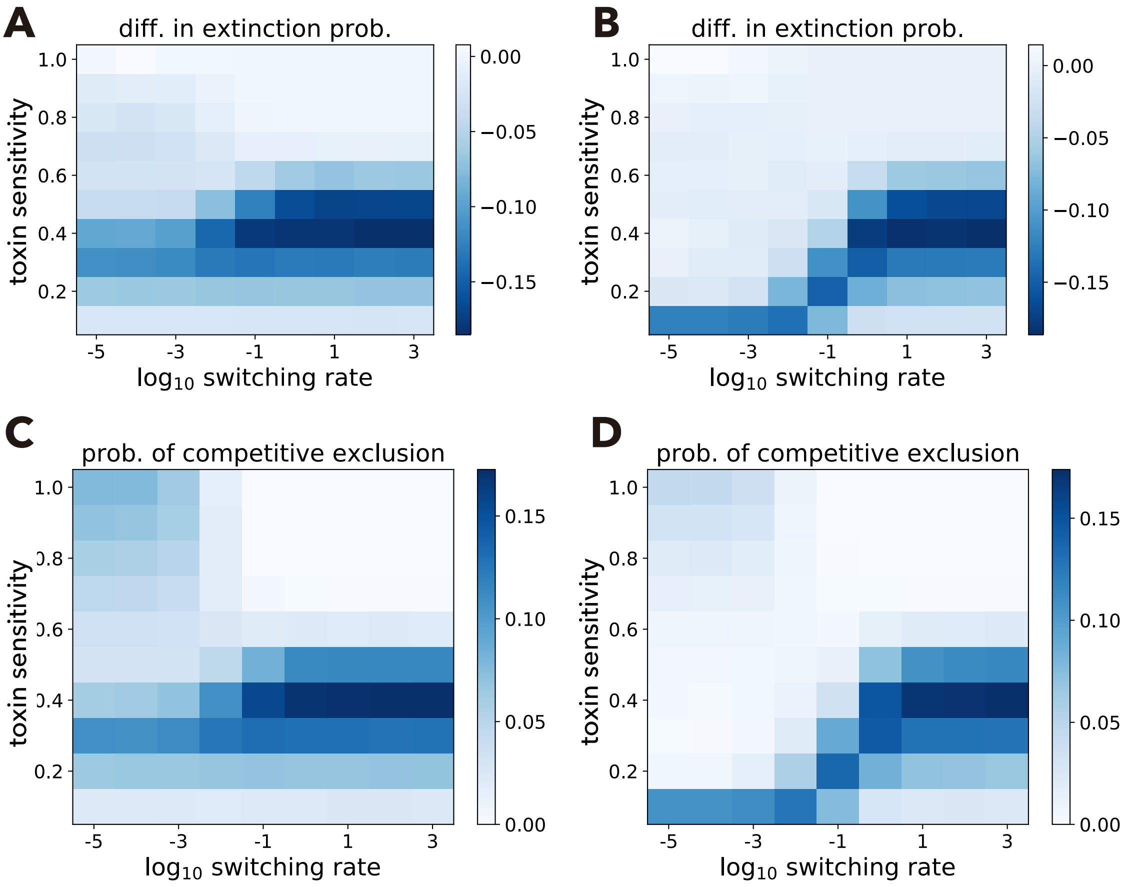


Figure A.7: Effects of the environmental switching rate in alternative scenarios

Examples of the effect of switching rate in alternative scenarios. In the left column (A and C), toxin supply is switching (scenario 2), while both resource and toxin supplies switch and are negatively correlated (scenario 3) in the right column (B, D). A and B: difference between extinction probabilities in absence and presence of species 2. C and D: competitive exclusion probability. Parameter values: $R_1^+ = 200$, $R_1^- = 50$, $T_1^+ = 200$, and $T_1^- = 50$.

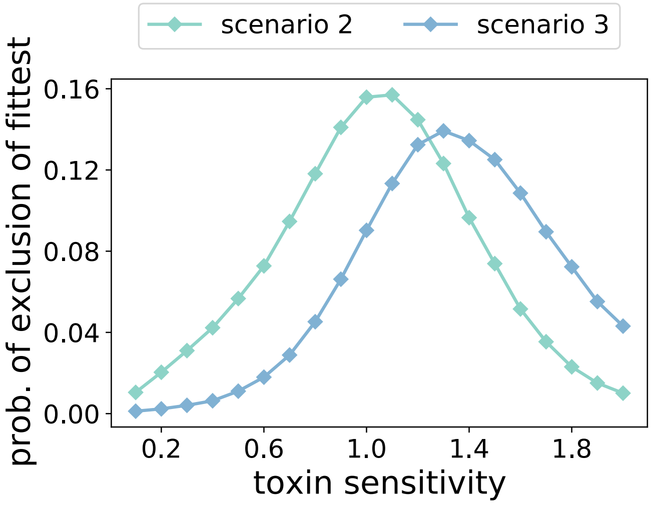


Figure A.8: Critical toxin sensitivities under mild environments

The critical toxin sensitivities (i.e., toxin sensitivity that maximizes the probability of exclusion of the fittest in the absence of environmental switching) under the mild environments (scenario 2 :scarce toxin supply $T_1^- = 50$, and scenario 3: abundant resource supply $R_1^+ = 200$ and scarce toxin supply) are $> 1$.

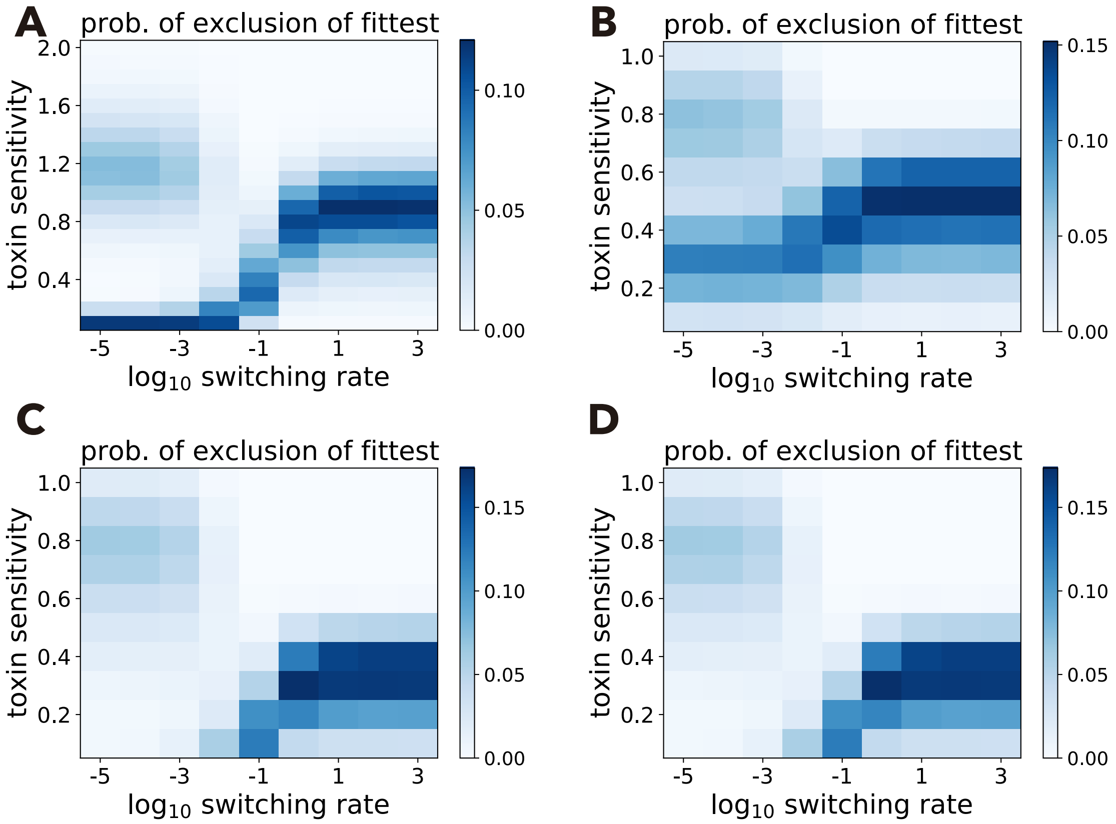


Figure A.9: Effects of resource supplies on exclusion of the fittest

Top: abundant resource supply becomes twice (A) or half (B) of $R_1^+ = 200$, i.e. $R_1^+ = 400$ in panel (A) and $R_1^+ = 100$ in panel (B). Bottom: scarce resource supply becomes twice (C) or half (D) of $R_1^- = 50$, i.e. $R_1^- = 100$ in panel (C) and $R_1^- = 25$ in panel (D).

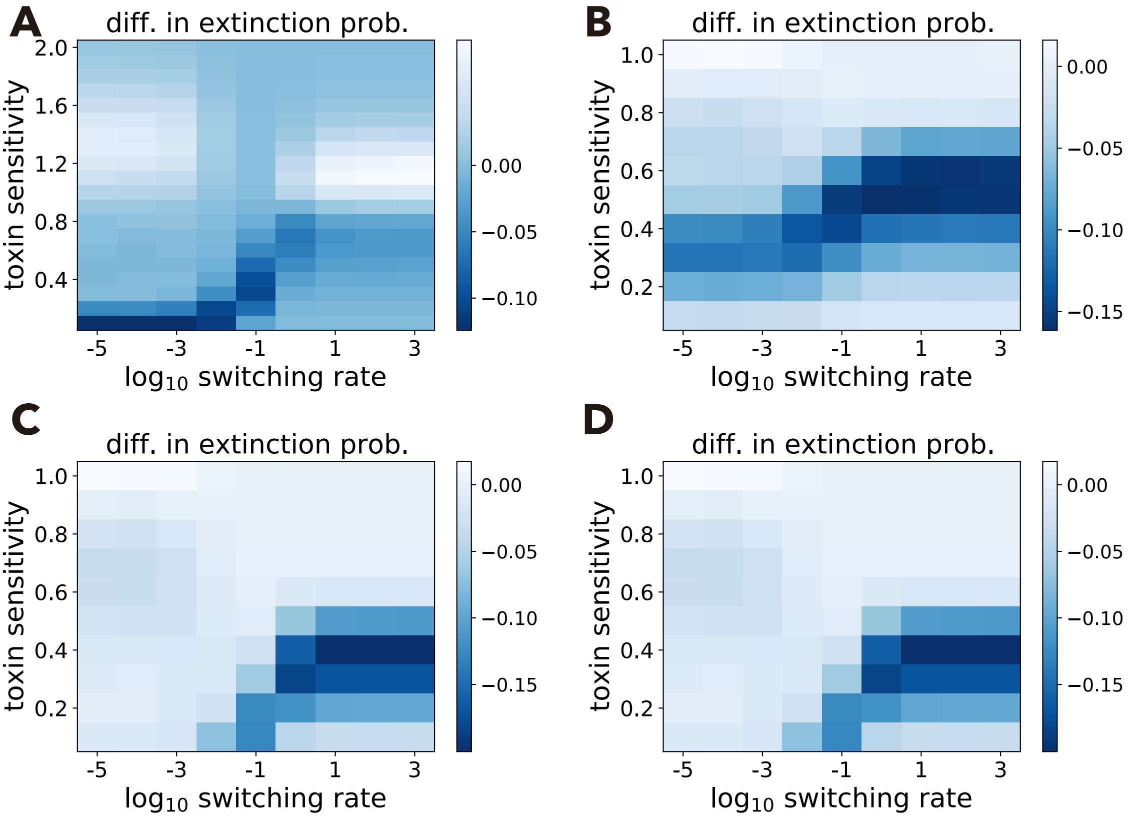


Figure A.10: Effects of resource supplies on difference in extinction probability

Similar to Fig. A.9 but showing species 2's effect on species 1's extinction probability. Top: abundant resource supply becomes twice (A) or half (B) of $R_1^+ = 200$, i.e. $R_1^+ = 400$ in panel (A) and $R_1^+ = 100$ in panel (B). Bottom: the scarce resource supply becomes twice (C) or half (D) of $R_1^- = 50$, i.e. $R_1^- = 100$ in panel (C) and $R_1^- = 25$ in panel (D). We plotted toxin sensitivity from 0.1 to 2.0 in panel A to see non-monotonic positive species interactions (see also Fig.A.11 )

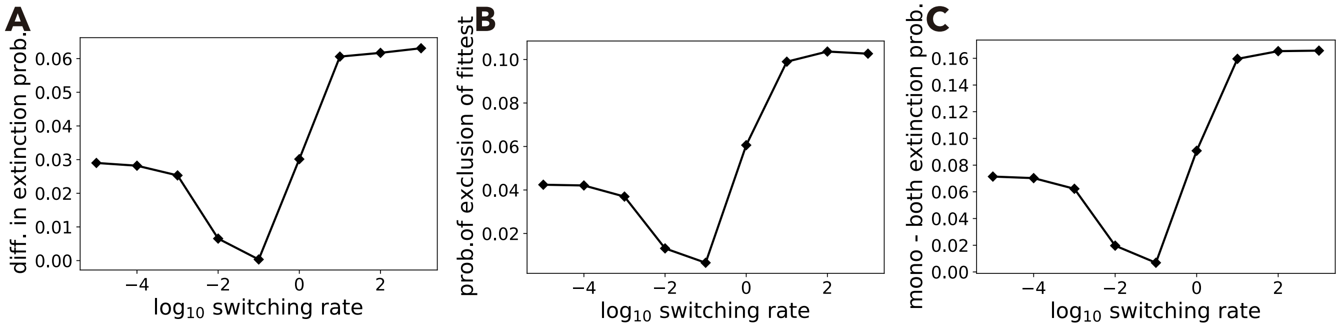


Figure A.11: Positive interaction strength varies non-monotonically with switching rate

A: Difference in species 1's extinction probability with positive sign at $\delta = 1.0$ and $R_1^+ = 400$, showing a non-monotonic effect of the environmental switching rate. B: Probability of exclusion of the fittest. C: Difference between the probability of species 1 going extinct alone and both species going extinct.

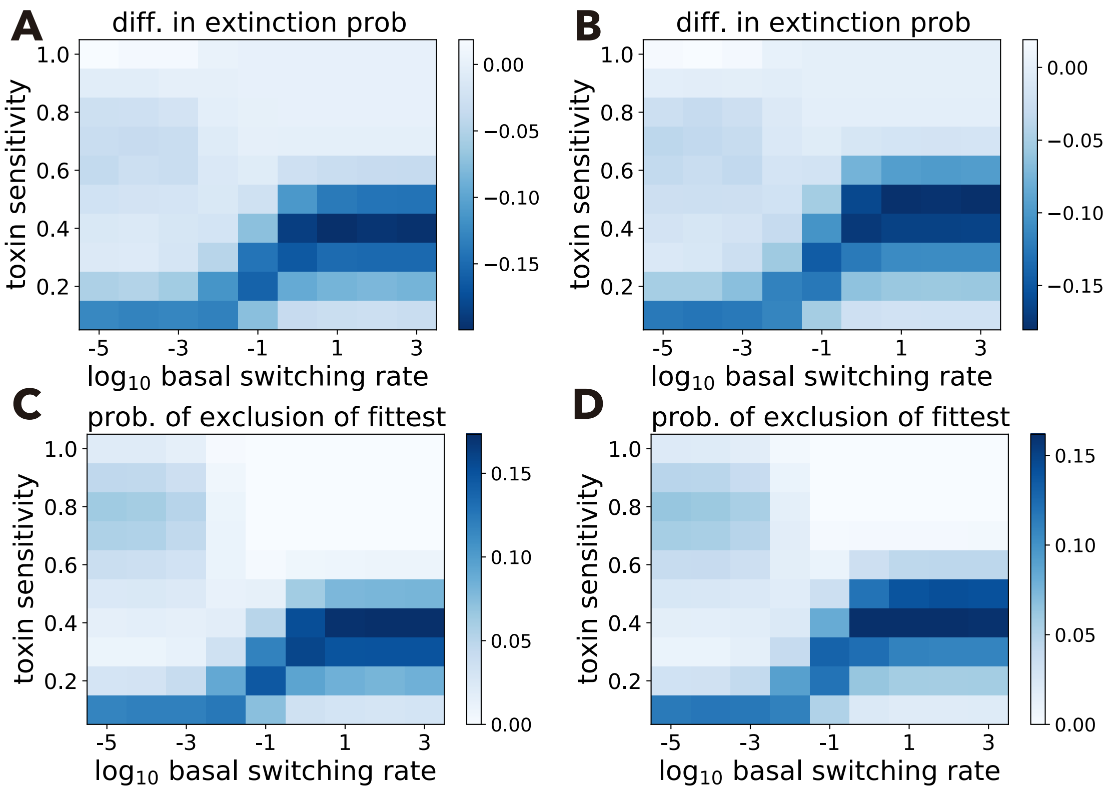


Figure A.12: Extinction probabilities under asymmetrically switching environments

Difference of species 1's extinction probability and the probability of exclusion of the fittest under asymmetric switching rates over the baseline of the switching rate $\nu$ and toxin sensitivity. Here, environmental fluctuations change the amounts of resource supplies. A and B: Difference of species 1's extinction probabilities in mono-culture minus co-culture wit species 2 when (A) the harsh environment continues longer than the mild environment ($\beta_1 = 1.2$ and $\beta_2 = 1$), or when (B) the mild environment continues longer than the harsh environment ($\beta_1 = 1$ and $\beta_2 = 1.2$), respectively. C and D: Probabilities that species 2 excludes species 1 when the environmental switching rates are identical to panels A and B, respectively. Other parameter values are shown in Table A.1.

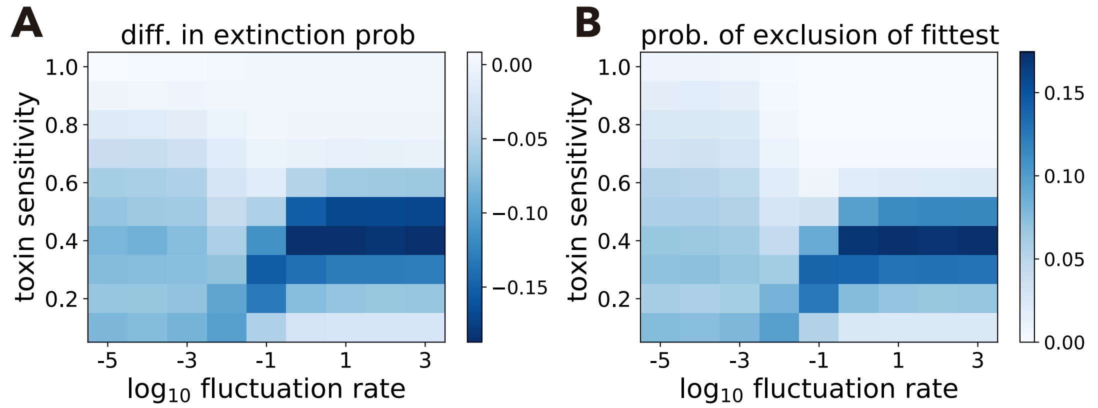


Figure A.13: Extinction probabilities under cyclic environmental changes

Difference in species 1's extinction probability and the probability of exclusion of the fittest under under cyclically fluctuating resource supplies among four states: $R_1\,(\xi = 1) = 200$, $R_1\,(\xi = 2) = 150$, $R_1\,(\xi = 3) = 100$, and $R_1\,(\xi = 4) = 50$. A: Difference of species 1's extinction probabilities in mono-culture minus co-culture with species 2. B: Probability that species 1 goes extinct but species 2 survives. Other parameter values are shown in Table A.1.

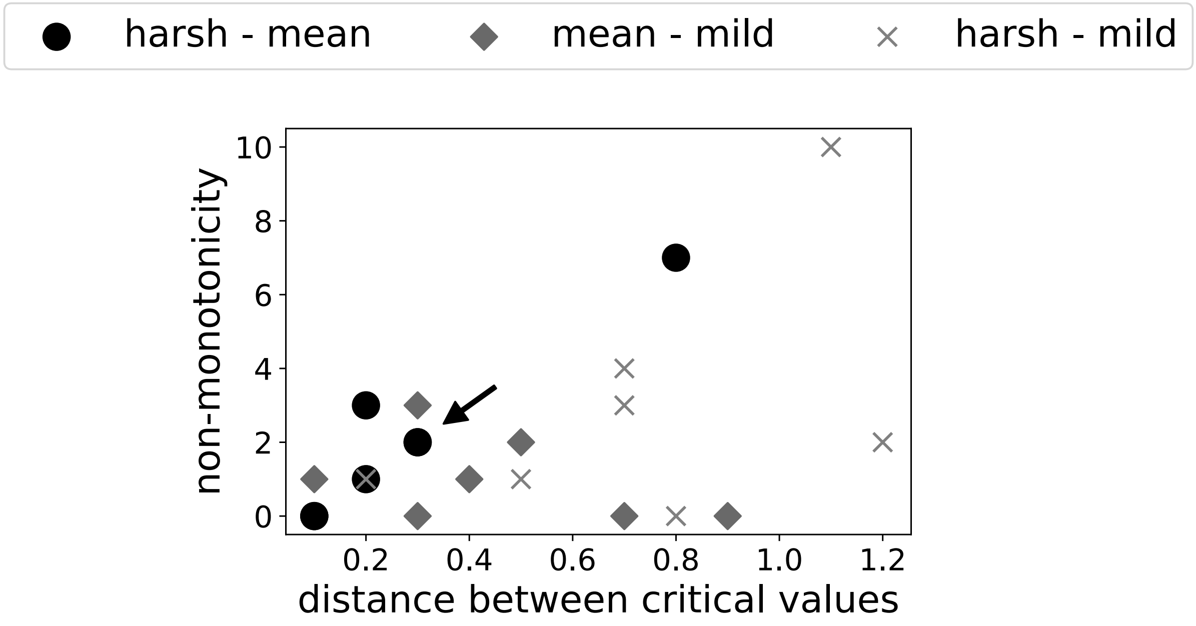


Figure A.14: Non-monotonicity and critical toxin sensitivities

The number of times we observe non-monotonic changes of species 1's difference in extinction probability across the explored parameter range varies with the distance between the two critical toxin sensitivities, depending on where distances are measured (between harsh and mean environments: dots, between mean and mild: environment diamonds, and between harsh and mild environments: crosses). These three distances were measured in each of the following seven scenarios: three different scenarios of environmental switching (Table 1 and Appendix 3) and four environmental switching scenario 1s with changing amounts of resource supplies (Appendix 4). The correlation is only significantly positive for the distance between scarce resource or abundant toxin supplies (i.e., harsh environments) and mean resource/toxin supplies (Spearman's $\rho = 0.77$, P-value: 0.043). The dot indicated by the arrow corresponds to the scenario analyzed in the main text.

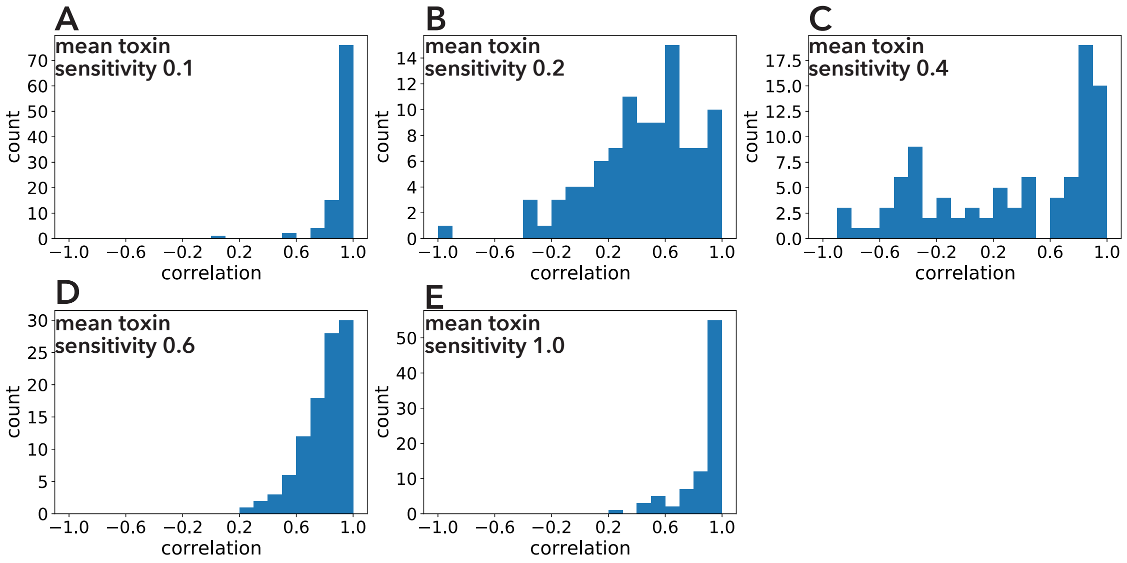


Figure A.15: Correlations between exclusion of the fittest and beta diversity in two-species communities

The Pearson correlation coefficients between probability of exclusion of the fittest (first column of Fig. 6) and beta diversity (second column of Fig. 6) in two-species communities are shown. Each panel differs in mean toxin sensitivity (A: 0.1, B: 0.2, C: 0.4, D: 0.6, and E: 1.0) and we analyzed 100 two-species communities in each case.

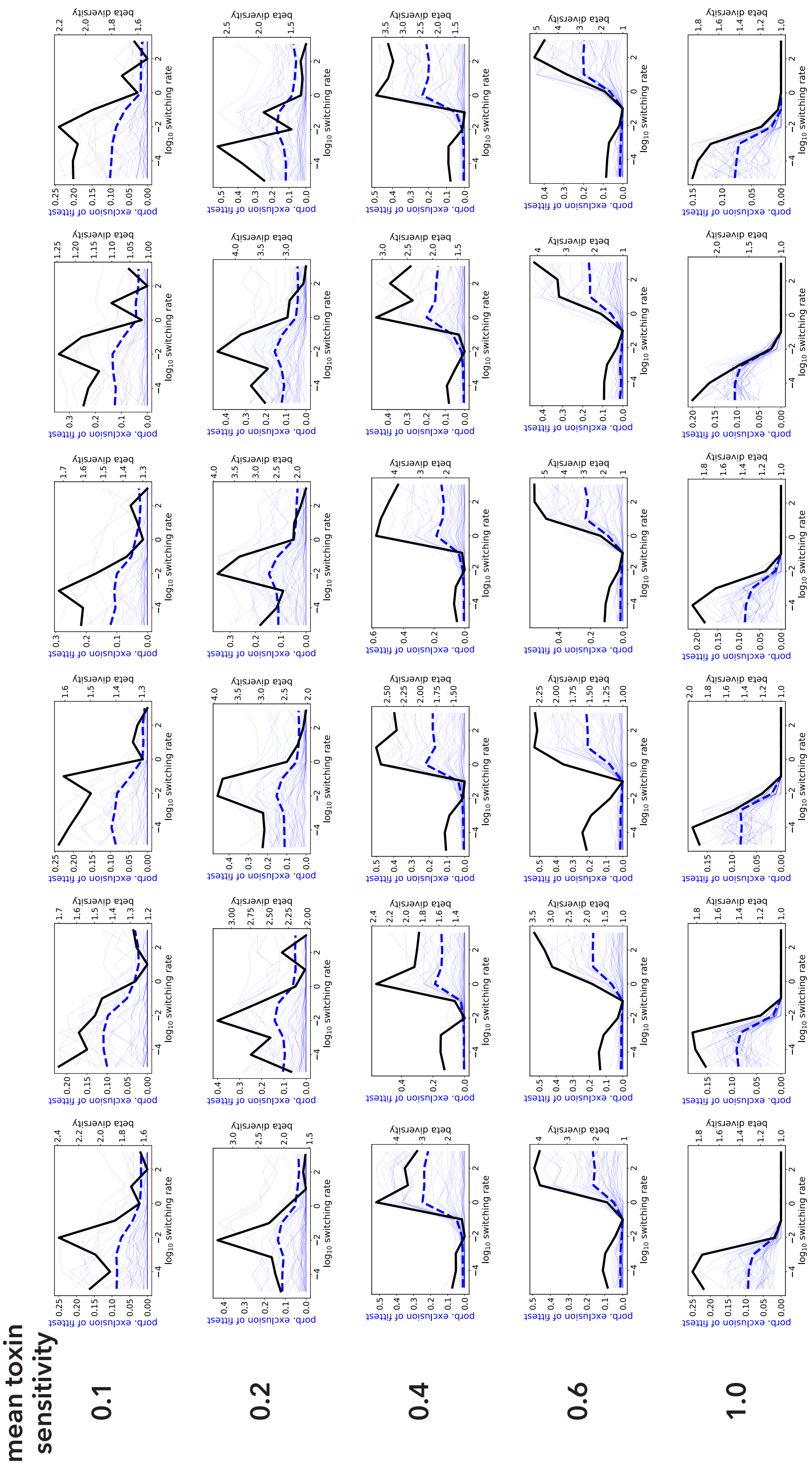


Figure A.16: Beta diversity and exclusion of the fittest in ten-species communities

Each panel represents the beta diversity of a ten-species community (black lines), probabilities of exclusion of the fittest in 45 species pairs (solid blue lines) and the mean probability of exclusion of the fittest over the 45 pairs (dashed blue lines). Each row corresponds to mean toxin sensitivity $\bar{\delta}$. One can see six examples of ten-species communities at each mean toxin sensitivity.

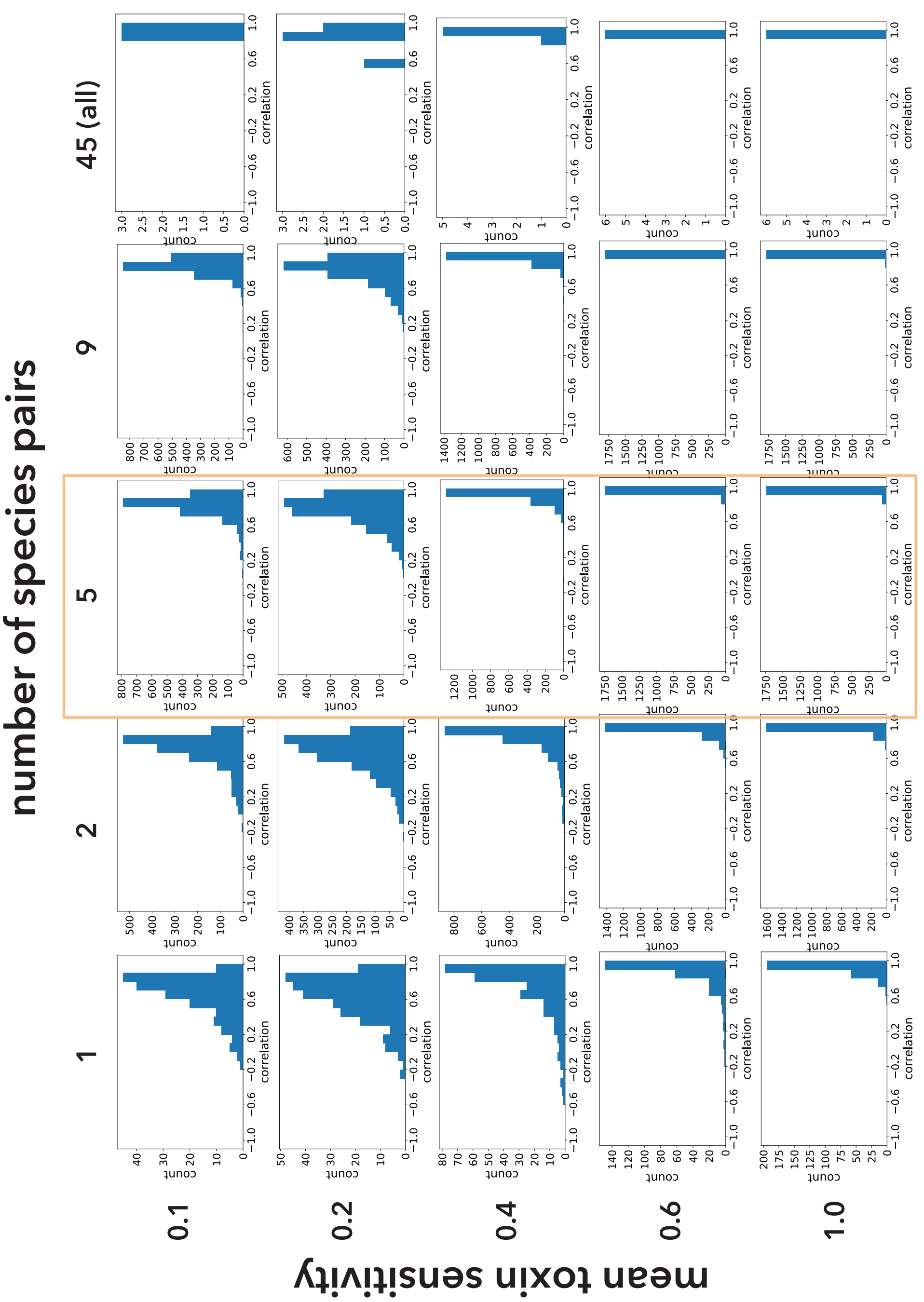


Figure A.17: Correlations between exclusion of the fittest and beta diversity in ten-species communities

The Pearson correlation coefficients between mean probability of exclusion of the fittest and beta diversity (Fig. A.16) in ten-species communities are shown. Each row represents the different value of mean toxin sensitivity $\bar{\delta}$ while each column represents the different number of sampled species pairs $m$.

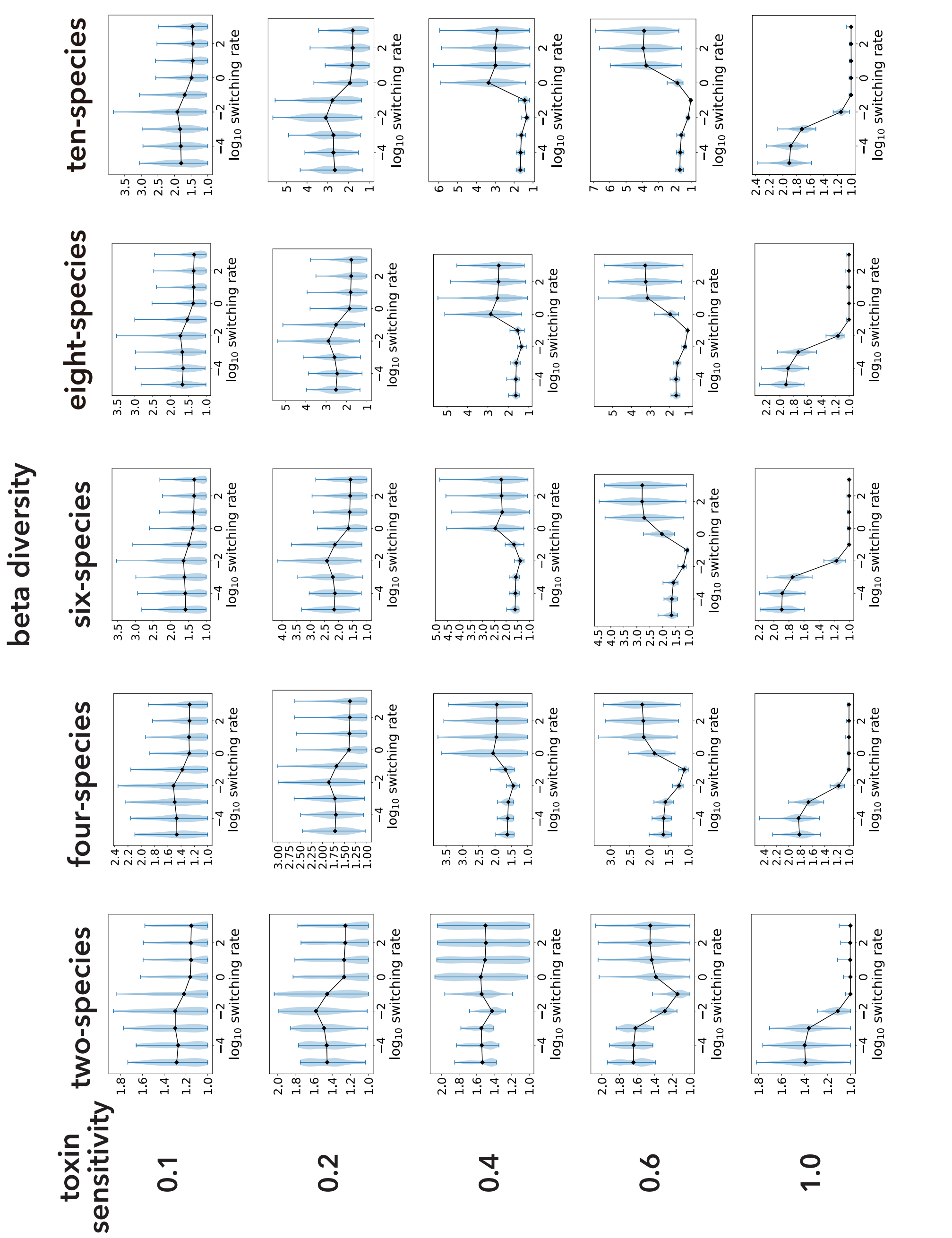


Figure A.18: Beta diversity with increasing initial number of species in a community

Beta diversities with increasing initial numbers of species $N$ and mean toxin sensitivities $\bar{\delta}$. The black lines show the means and blue areas represent the probability distributions calculated by 10'000 simulations (100 beta diversity measurements using different parameter sets, each from 100 replicate runs).

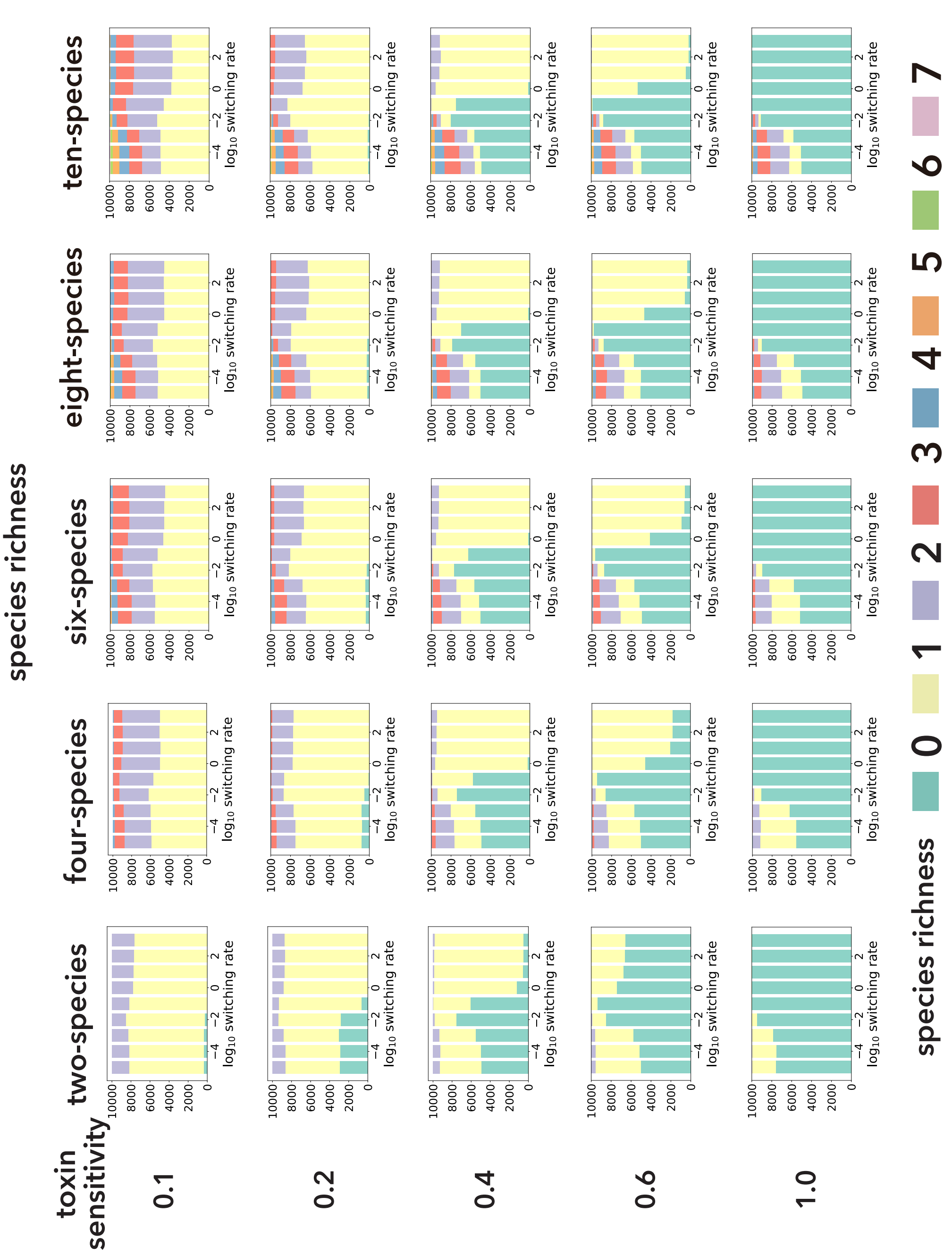


Figure A.19: Species richness with increasing initial number of species in a community

Species richness with increasing initial numbers of species $N$ and mean toxin sensitivities $\bar{\delta}$. Each bar plot represents results of 10'000 simulations.

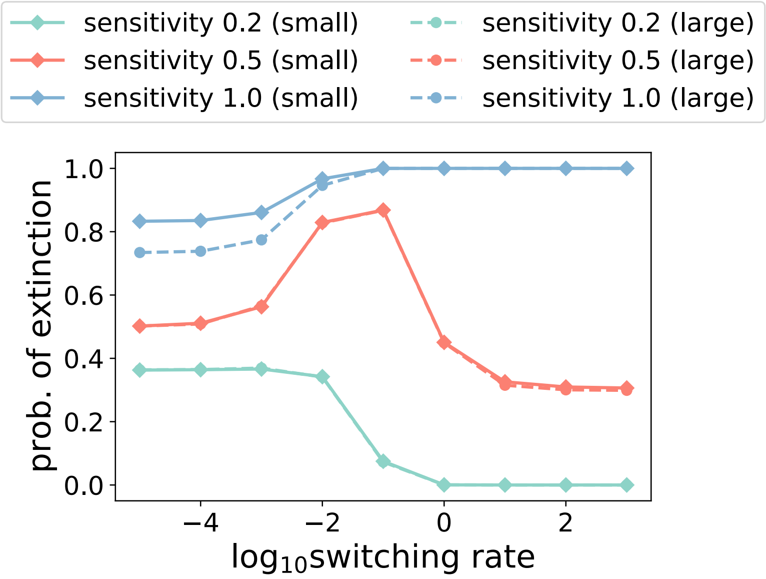


Figure A.20: Initial Population size's effect on extinction probabilities

Extinction probability of species 1 in mono-culture when the initial species abundance is default (small: $s_1(0) = 10$) or larger (large: $s_1(0) = 20$). When the switching rate is small and the toxin sensitivity is 1.0, the extinction probability is lower with the larger initial species abundance. In the rest cases, the extinction probabilities are not affected by the initial species abundance. The parameter values are as shown by Table A.1.

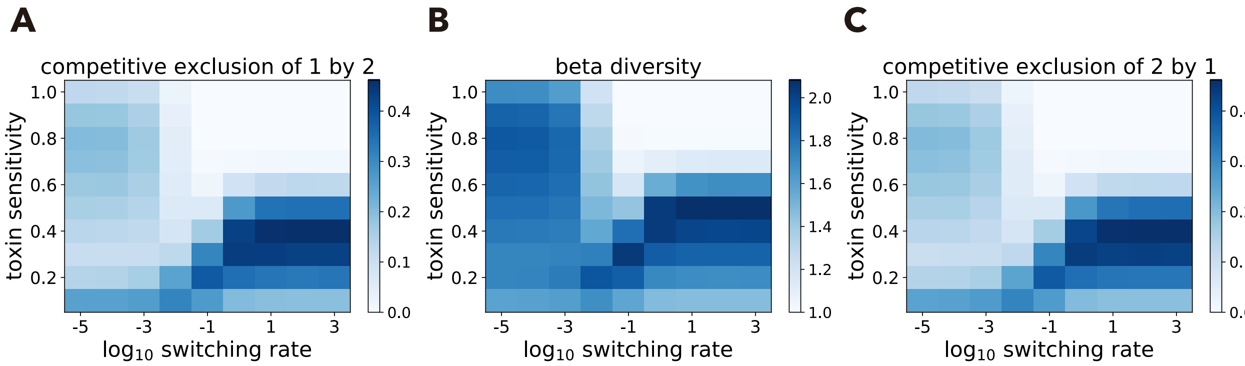


Figure A.21: Exclusion probabilities and beta diversity in neutral cases

Two species dynamics under neutral cases (i.e., two species differ only in their labels). In this case, the probabilities that species 2 outcompete species 1 (A) are identical to those that species 1 outcompete species 2 (C). Without loss of generality, we can call exclusion of either species as exclusion of the fittest in the neutral scenarios. As in Fig. 6, there are similarities between how the exclusion of the fittest (A or C) and beta diversity (B) changes over the switching rate at each toxin sensitivity. The parameter values are as shown by Table A.1.

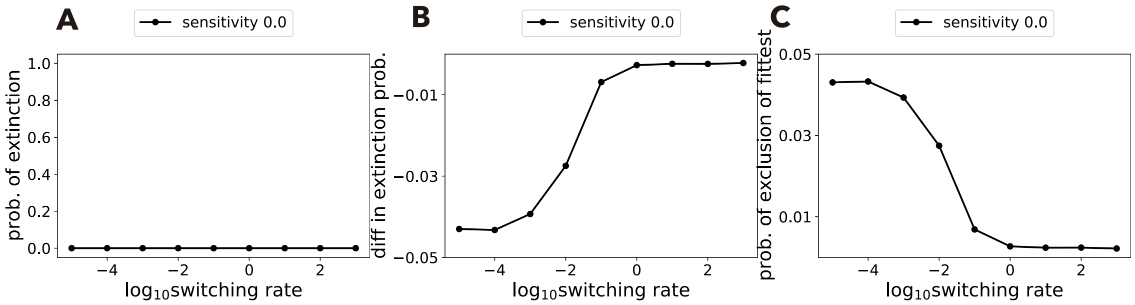


Figure A.22: Extinction probabilities with zero toxin sensitivity

When species's toxin sensitivities are zero, the dynamics are equivalent with the case without toxins because species die only due to dilution. A: Species 1 does not go extinct in mono-culture. B: Difference of species 1's extinction probabilities in mono-culture minus co-culture. C: Probability that species 2 outcompetes species 1 in co-culture. Panels B and C show qualitatively similar results with the case of toxin sensitivity 0.1, see Figs. 3A and C because species 2 can outcompete species 1 under the harsh environment with low switching rates, due to DN.

Table A.1: List of fixed parameters in the interaction analysis

| Symbol | Value | Description |
|---|---|---|
| $\alpha$ | 0.1 | dilution rate of the chemostat |
| $R_1^{\pm}$ | $R_1^+ = 200, R_1^- = 50$ | abundant or scarce resource supply concentration |
| $T_1^{\pm}$ | $T_1^+ = 200, T_1^- = 50$ | abundant or scarce toxin supply concentration |
| $Y_{1k}^r$ | $Y_{1k}^r = 1$ for $k = 1, 2$ | species $k$'s biomass yields of resource |
| $Y_{1k}^t$ | $Y_{1k}^t = 1$ for $k = 1, 2$ | species $k$'s biomass yields of toxin |
| $\mu_{11}$ | 1.0 | maximum growth rate of species 1 on resource 1 |
| $\mu_{12}$ | 0.91 | maximum growth rate of species 2 on resource 1 |
| $\delta_{1k}$ | $[0.1, \ldots, 1.0]$ and $\delta = \delta_{11} = \delta_{12}$ | sensitivity of species $k$ to toxin 1. |
| $K_{1k}^r$ | 100 | amount of resource 1 that gives half-max growth rate of species $k$ |
| $K_{1k}^t$ | 100 | amount of toxin 1 that gives half-max death rate of species $k$ |

Table A.2: Summary of critical toxin sensitivities and number of times non-monotonicity is observed

| Switching scenario | | 1 | 2 | 3 | 1 | | | |
|---|---|---|---|---|---|---|---|---|
| Amounts of supplies | | base line * | | | $\uparrow R_1^+$ | $\downarrow R_1^+$ | $\uparrow R_1^-$ | $\downarrow R_1^-$ |
| Critical toxin | harsh | 0.1 | 0.3 | 0.1 | 0.1 | 0.1 | 0.3 | 0.1 |
| sensitivities | mean | 0.4 | 0.4 | 0.4 | 0.9 | 0.2 | 0.5 | 0.3 |
| | mild | 0.8 | 1.1 | 1.3 | 1.2 | 0.3 | 0.8 | 0.8 |
| number of | mean and harsh | 2 | 0 | 2 | 7 | 0 | 1 | 3 |
| non-monotonic changes | mean and mild | 1 | 0 | 0 | 3 | 0 | 0 | 2 |
| observed between | mild and harsh | 3 | 0 | 2 | 10 | 0 | 1 | 4 † |

* For exact parameter values, see Table A.1.

† At the critical toxin sensitivity corresponding to the mean environment ($\delta = 0.3$), the species interaction non-monotonically changes over the switching rate: the frequency of non-monotonicity is $3 + 2 - 1 = 4$.
